# A dynamic 6,000-year genetic history of Eurasia’s Eastern Steppe

**DOI:** 10.1101/2020.03.25.008078

**Authors:** Choongwon Jeong, Ke Wang, Shevan Wilkin, William Timothy Treal Taylor, Bryan K. Miller, Sodnom Ulziibayar, Raphaela Stahl, Chelsea Chiovelli, Jan H. Bemmann, Florian Knolle, Nikolay Kradin, Bilikto A. Bazarov, Denis A. Miyagashev, Prokopiy B. Konovalov, Elena Zhambaltarova, Alicia Ventresca Miller, Wolfgang Haak, Stephan Schiffels, Johannes Krause, Nicole Boivin, Erdene Myagmar, Jessica Hendy, Christina Warinner

## Abstract

The Eastern Eurasian Steppe was home to historic empires of nomadic pastoralists, including the Xiongnu and the Mongols. However, little is known about the region’s population history. Here we reveal its dynamic genetic history by analyzing new genome-wide data for 214 ancient individuals spanning 6,000 years. We identify a pastoralist expansion into Mongolia ca. 3000 BCE, and by the Late Bronze Age, Mongolian populations were biogeographically structured into three distinct groups, all practicing dairy pastoralism regardless of ancestry. The Xiongnu emerged from the mixing of these populations and those from surrounding regions. By comparison, the Mongols exhibit much higher Eastern Eurasian ancestry, resembling present-day Mongolic-speaking populations. Our results illuminate the complex interplay between genetic, sociopolitical, and cultural changes on the Eastern Steppe.

## Introduction

Recent paleogenomic studies have revealed a dynamic population history on the Eurasian Steppe, with continental-scale migration events on the Western Steppe coinciding with Bronze Age transformations of Europe, the Near East, and the Caucasus (Allentoft et al., 2015; de Barros Damgaard et al., 2018; Damgaard et al., 2018; Haak et al., 2015; Mathieson et al., 2015; Wang et al., 2019). However, despite advances in understanding the genetic prehistory of the Western Steppe, the prehistoric population dynamics on the Eastern Steppe remain poorly understood (de Barros Damgaard et al., 2018; Jeong et al., 2018). The Eastern Steppe is a great expanse of grasslands, forest-steppe, and desert-steppe extending more than 2,500 km from the Altai-Sayan mountain range in the west to northeastern China in the east (Fig. 1). While also covering parts of modern-day China and Russia, most of the Eastern Steppe falls within the national boundaries of present-day Mongolia (Fig. S1). The Eastern Steppe has been occupied since the early Upper Paleolithic (ca. 34,000 cal BP) (Devièse et al., 2019), and recent paleogenomic studies suggest that the eastern Eurasian forest-steppe zone was genetically structured during the Pre-Bronze and Early Bronze Age periods, with a strong west-east admixture cline of ancestry stretching from Botai in central Kazakhstan to Lake Baikal in southern Siberia to Devil’s Gate Cave in the Russian Far East (de Barros Damgaard et al., 2018; Jeong et al., 2018; Sikora et al., 2019; Siska et al., 2017).

**Fig. 1.**
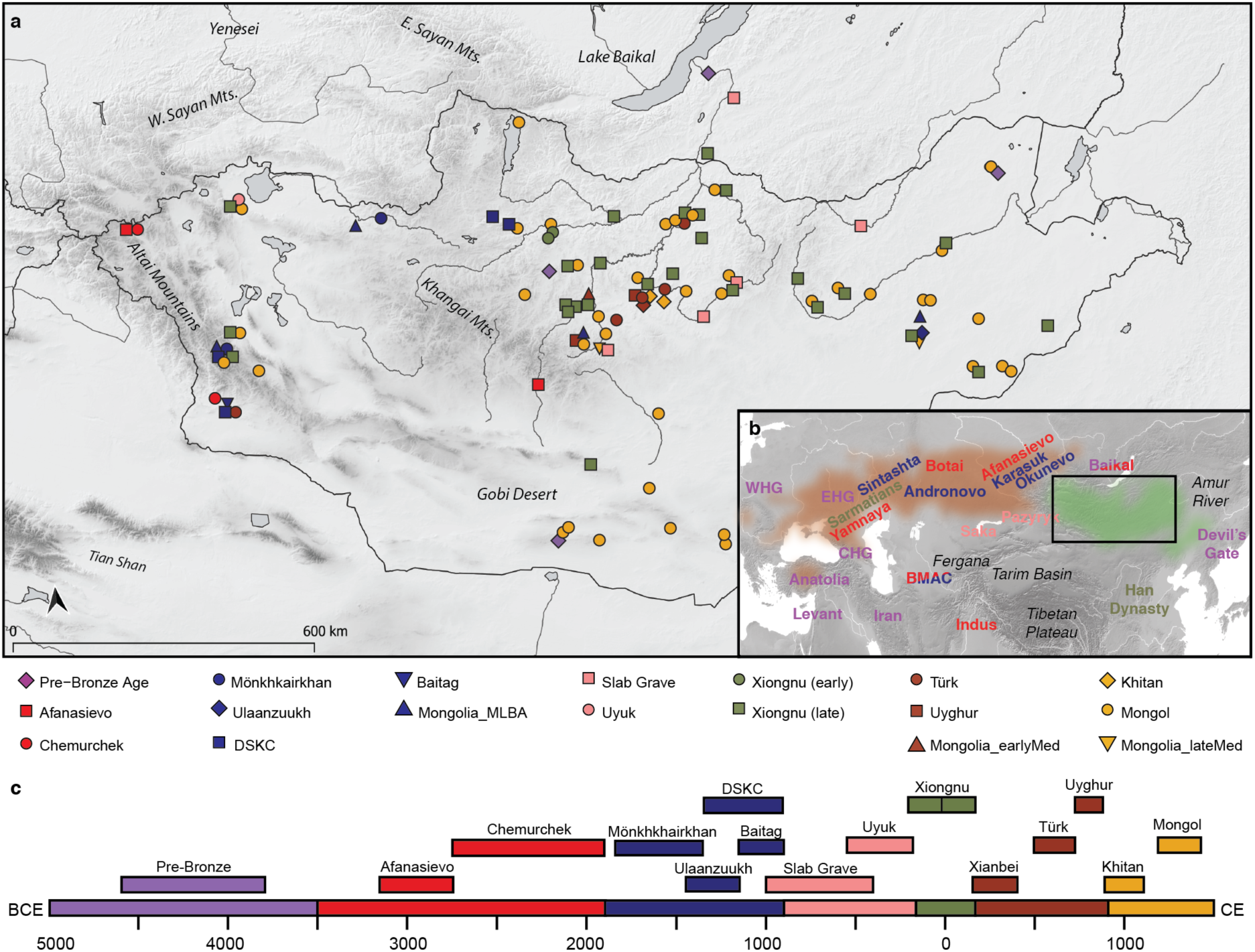
Overview of ancient populations and time periods. (**a**), Distribution of sites with their associated culture and time period indicated by color: Pre-Bronze, purple; Early Bronze, red; Middle/Late Bronze, blue; Early Iron, pink; Xiongnu, green; Early Medieval, brown; Late Medieval, gold (See supplementary materials). See Table S2 for site codes and labels. (**b**), Inset map of Eurasia indicating area of present study (box) and the locations of other ancient populations referenced in the text, colored by time period. The geographic extent of the Western/Central Steppe is indicated in light brown, while the Eastern Steppe is indicated in light green. (**c**) Timeline of major temporal periods and archaeological cultures in Mongolia. Site locations have been jittered to improve visibility of overlapping sites.

During the Bronze Age, the multi-phased introduction of pastoralism drastically changed lifeways and subsistence on the Eastern Steppe (Honeychurch, 2015; Kindstedt and Ser-Od, 2019). A recent large-scale paleoproteomic study has confirmed milk consumption in Mongolia prior to 2500 BC by individuals affiliated with the Afanasievo (3000 BCE) and Chemurchek (2750-1900 BCE) cultures (Wilkin et al., 2019). Although Afanasievo groups in the Upper Yenisei region have been genetically linked to the Yamnaya culture of the Pontic-Caspian steppe (ca. 3300-2200 BCE) (Allentoft et al., 2015; Morgunova and Khokhlova, 2013), the origins of the Chemurchek have been controversial (Kovalev, 2014). Once introduced, ruminant dairying became widespread by the Middle/Late Bronze Age (MLBA, here defined as 1900-900 BCE), being practiced in the west and north at sites associated with Deer Stone-Khirigsuur Complex (DSKC) and in the east in association with the Ulaanzuukh culture (Jeong et al., 2018; Wilkin et al., 2019). The relationships between DSKC and Ulaanzuukh groups are poorly understood, and little is known about other MLBA burial traditions in Mongolia, such as the Mönkhkhairkhan and Baitag (See supplementary materials). By the end of the second millennium BCE, the previously powerful MLBA cultures were in decline, and political power shifted during the Early Iron Age to the Slab Grave culture (ca. 1000-300 BCE), whose burials often incorporate uprooted materials from DSKC monuments (Honeychurch, 2015), and to the Uyuk culture (ca. 700-200 BCE) of the Sayan mountains to the northwest (also known as the Aldy-Bel culture), who had strong cultural ties to the Pazyryk (ca. 500-200 BCE) and Saka (ca. 900-200 BCE) cultures of the Altai and eastern Kazakhstan (Savinov, 2002; Tseveendorj, 2007).

From the late first millennium BC onwards, a series of hierarchical and centrally organized empires arose on the Eastern Steppe, notably the Xiongnu (209 BCE-98 CE), Türkic (552-742 CE), Uyghur (744-840 CE), and Khitan (916-1125 CE) empires. The Mongol empire, emerging in the 13^th^ century CE, was the last and most expansive of these regimes, eventually controlling vast territories and trade routes stretching from China to the Mediterranean. However, due to a lack of large-scale genetic studies, the origins and relationships of the people who formed these states, including both the ruling elites and local commoners, remain obscure (See supplementary materials).

To clarify the population dynamics on the Eastern Steppe since prehistory, we generated and analyzed genome-wide genetic datasets for 214 individuals from 85 Mongolian and 3 Russian sites spanning approximately 6,000 years of time (ca. 4600 BCE to 1400 CE) (Tables S1-S9 and Fig. S2-S8) (See supplementary materials). To this, we added recently published genomic data for 18 Bronze Age individuals from northern Mongolia (Jeong et al., 2018), as well as datasets from neighboring ancient populations in Russia and Kazakhstan (de Barros Damgaard et al., 2018; Damgaard et al., 2018; Narasimhan et al., 2019; Sikora et al., 2019; Unterländer et al., 2017) (Table S10), which we analyze together with worldwide modern reference populations (Table S11; Fig. S4). We also generated 24 new accelerator mass spectrometry dates (Table S12), supplementing 81 previously published radiocarbon dates from Mongolia (Jeong et al., 2018; Taylor et al., 2019), for a total of 105 directly dated individuals in this study (See supplementary materials).

## Results

### Pre-Bronze Age population structure and the arrival of pastoralism

Throughout the mid-Holocene and before the Bronze Age, the Eastern Steppe was sparsely populated by hunter-gatherers, whose lives and activities are recorded in petroglyphs throughout the region (Jacobson-Tepfer and Meacham, 2010; Kindstedt and Ser-Od, 2019). In this study, we analyzed six pre-Bronze Age individuals from three sites dating to the fifth and fourth millennia BCE: one from eastern Mongolia (SOU001, “eastMongolia_preBA”, 4686-4495 cal. BCE), one from central Mongolia (ERM003, “centralMongolia_preBA”, 3781-3643 cal. BCE), and four from the eastern Baikal region (“Fofonovo_EN”). By comparing these genomes to previously published ancient and modern data across Eurasia (Fig. 2) (see Methods and Materials), we found that they are most closely related to contemporaneous hunter-gatherers from the western Baikal region (“Baikal_EN”, 5200-4200 BCE) and the Russian Far East (“DevilsCave_N”, ca. 5700 BCE), filling in the geographic gap in the distribution of this genetic profile (Fig. 3a). We refer to this profile as “Ancient Northeast Asian” (ANA) to reflect its geographic distribution relative to another widespread mid-Holocene genetic profile known as “Ancient North Eurasian” (ANE), which is found among the Pleistocene hunter-gatherers of the Mal’ta (ca. 24500-24100 BP) and Afontova Gora (ca. 16900-16500 BP) sites in Siberia (Fu et al., 2016; Raghavan et al., 2015) and the horse-herders of Botai, Kazakhstan (ca. 3500-3300 BCE) (de Barros Damgaard et al., 2018). In PCA (Fig. 2), ancient ANA individuals fall close to the cluster of present-day Tungusic- and Nivkh-speaking populations from the lower Amur River region in northeast Asia, indicating that their genetic profile is still present in indigenous populations of the Far East today (Fig. S5). EastMongolia_preBA is genetically indistinguishable from DevilsCave_N (Fig. 3a; Table S13; Fig. S9-S10), whereas Fofonovo_EN and the slightly later centralMongolia_preBA both derive a minority (12-17%) of their ancestry from ANE-related (Botai-like) groups with the remainder of their ancestry (83-87%) characterized as ANA (Fig. 3a; Table S13). Reanalyzing published data from the western Baikal early Neolithic Kitoi culture (Baikal_EN) and the early Bronze Glazkovo culture (Baikal_EBA) (de Barros Damgaard et al., 2018), we find that they have similar ancestry profiles and a slight increase in ANE ancestry through time (from 6.4% to 20.1%) (Fig. 3a). Overall, these data suggest broader genetic shifts in hunter-gatherer ancestry at this time.

**Fig. 2.**
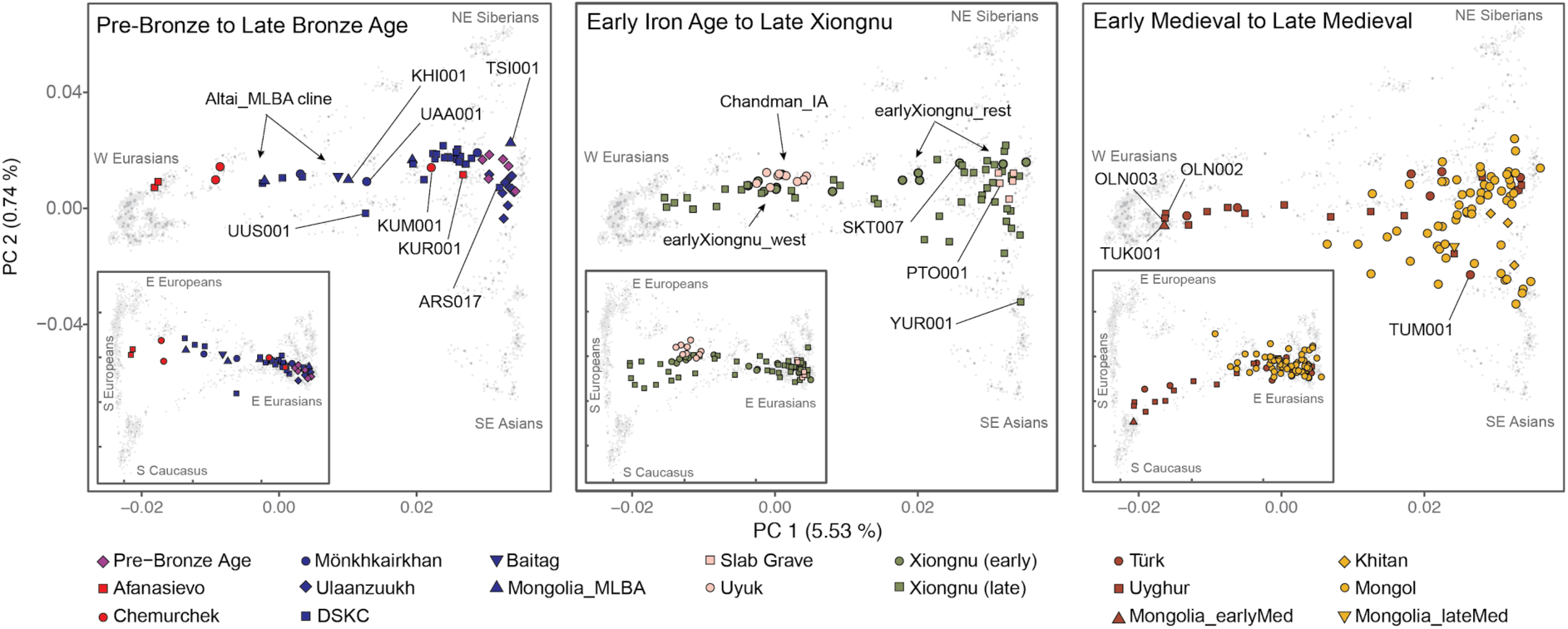
Genetic structure of Mongolia through time. Principal component analysis (PCA) of ancient individuals (n=214) from three major periods projected onto contemporary Eurasians (gray symbols). Main panels display PC1 vs PC2; insets display PC1 vs PC3. Inset tick marks for PC1 correspond to those for the main panels; PC3 accounts for 0.35% of variation. See Fig. S5 for population, sample, and axis labels, and Tables S2-S4 for further site and sample details.

**Fig. 3.**
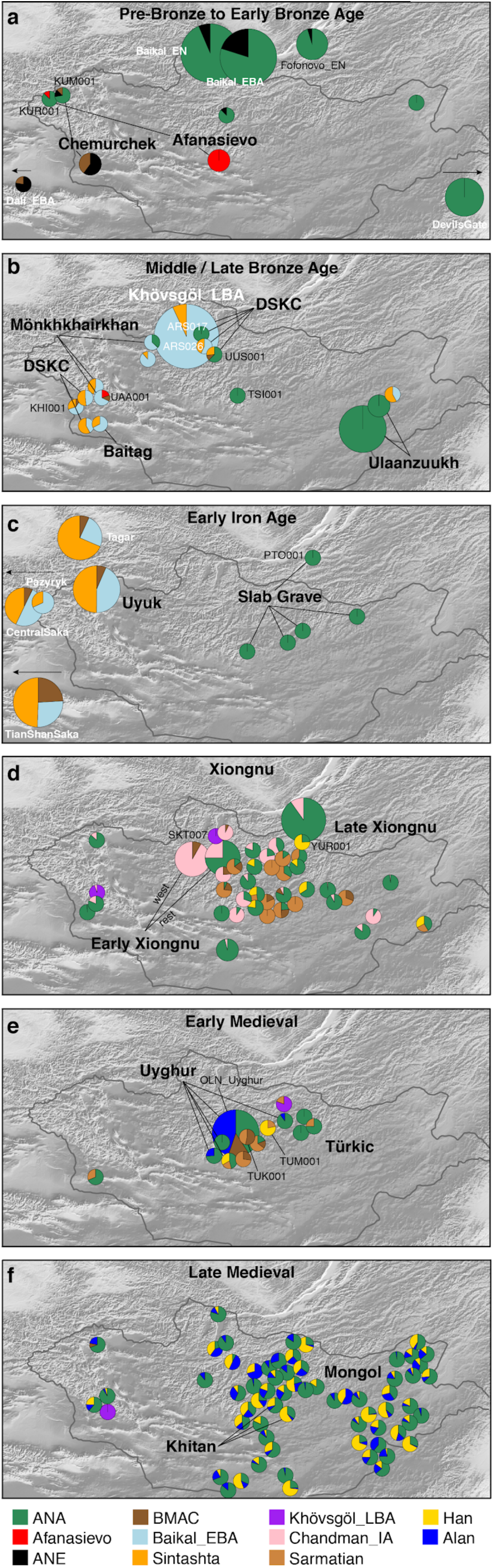
Genetic changes in the Eastern Steppe across time characterized by qpAdm. (**a**) Pre-Bronze through Early Bronze Age; (**b**) Middle/Late Bronze Age; (**c**) Early Iron Age; (**d**) Xiongnu period; (**e**) Early Medieval; (**f**) Late Medieval. Modeled ancestry proportions are indicated by sample size-scaled pie charts, with ancestry source populations shown below (See supplementary materials). For panels (**b-c**), Baikal_EBA is modeled as light blue; in (**d-f**), Khövsgöl_LBA (purple) and the Uyuk of Chandman_IA (pink) are modeled as new sources (Fig. S7). Cultural groups are indicated by bold text. For panels (**d-f**), individuals are Late Xiongnu, Türkic, and Mongol, respectively, unless otherwise noted. Previously published reference populations are noted with white text; all others are from this study. Populations beyond the map borders are indicated by arrows. Burial locations have been jittered to improve visibility of overlapping individuals. See Fig. S8 for additional labels.

Pastoralism in Mongolia is often assumed to have been introduced by the eastward expansion of Western Steppe cultures (e.g., Afanasievo) via either the Upper Yenisei and Sayan mountain region to the northwest of Mongolia or through the Altai mountains in the west (Janz et al., 2017). Although the vast majority of Afanasievo burials found to date are located in the Altai mountains and Upper Yenisei regions outside of Mongolia (Honeychurch, 2017), an Early Bronze Age (EBA) site in the southern Khangai Mountains of central Mongolia has yielded Afanasievo-style graves with proteomic evidence of ruminant milk consumption (Wilkin et al., 2019). Analyzing two of these individuals (Afanasievo_Mongolia, 3112-2917 cal. BCE), we find that their genetic profiles are indistinguishable from that of published Afanasievo individuals from the Yenisei region (Allentoft et al., 2015; Narasimhan et al., 2019) (Fig. 2, Fig. S11), and thus these two Afanasievo individuals confirm that the EBA expansion of Western Steppe herders (WSH) did not stop at the mountain ranges separating the Western and Eastern Steppes (Allentoft et al., 2015), but rather extended a further 1500 km eastwards into the heart of central Mongolia (Fig. 3a).

Although the Afanasievo-related expansion may have introduced dairy pastoralism into Mongolia (Janz et al., 2017), the people themselves do not seem to have left a long-lasting genetic footprint. An Afanasievo-style burial (Afanasievo_KUR001) excavated in the Mongolian Altai and dating to approximately 500 years later (2618-2487 cal. BCE), was found to have a distinct (Fig. 2) and mostly ANA (∼90%) derived ancestry (Fig. 3a), with only a minor component tracing back to the Afanasievo (∼10%; Table S14; Fig. S11).

The succeeding EBA Chemurchek culture (2750-1900 BCE), a ruminant dairying society (Wilkin et al., 2019) whose mortuary features include stone slabs and anthropomorphic stelae, has also been purportedly linked to WSH migrations (Kovalev and Erdenebaatar, 2009). Chemurchek graves are found throughout the Altai and in the Dzungar Basin in Xinjiang, China (Jia and Betts, 2010). We analyzed two Chemurchek individuals from the southern Altai site of Yagshiin Huduu and one from Khundii Gobi (KUM001) in the northern Altai. Compared to Afanasievo_Mongolia, the Yagshiin Huduu individuals also show a high degree of Western ancestry but are displaced in PCA (Fig. 2), having also a strong genetic affinity with ANE-related ancient individuals such as AfontovaGora3 (AG3), West_Siberia_N, and Botai (Fig. 3a; Fig. S9, S11). We find that these Chemurchek individuals (Chemurchek_Altai) are genetically similar to Dali_EBA (Fig. 3a), a contemporaneous individual from eastern Kazakhstan (Narasimhan et al., 2019). The genetic profiles of both the Yagshiin Huduu and Dali_EBA individuals are well fitted by two-way admixture models with Botai (60-78%) and groups with ancient Iranian-related ancestry, such as Gonur1_BA from Gonur Tepe, a key EBA site of the Bactria-Margiana Archaeological Complex (BMAC) (22-40%; Table S15; Fig. 3a), and ancient individuals from post-BMAC Bronze Age sites in southeastern Uzbekistan (Table S15). Although minor genetic contributions from the Afanasievo-related groups cannot be excluded, Iranian-related ancestry is required for all fitting models, and this admixture is estimated to have occurred 12±6 generations earlier (∼336±168 years; Fig. S12) when modeled using DATES (Narasimhan et al., 2019). Because all proxy source populations used in this modeling are quite distant in either time or space from the EBA Altai, the proximate populations contributing to the Chemurchek cannot be precisely identified, but future research in genetically unsampled regions such as Xinjiang may provide important clues.

In the northern Altai, the third Chemurchek individual in our study (KUM001) has additional northeast Asian ancestry, similar to levels seen in Afanasievo_KUR001 (Fig. 3a). KUM001 can be modeled as a mixture of Chemurchek-related (∼30%) and ANA ancestry (∼70%) (Table S15). The high proportions of local ANA ancestry in KUR001 and KUM001 suggest that the arrival of Afanasievo and later Chemurchek pastoralists did not necessarily displace local Altai peoples, despite the strong cultural and economic impact of these herders. Indeed, the cultural features that they first introduced, such as mortuary mound building and dairy pastoralism, continue to the present day. However, unlike in Europe, where migrating EBA steppe herders had a transformative and lasting genetic impact on local populations (Allentoft et al., 2015; Haak et al., 2015; Mathieson et al., 2018), we find that neither Afanasievo nor Chemurchek groups left significant or enduring genetic traces into the subsequent Middle and Late Bronze Ages (MLBA).

### Bronze Age emergence of a tripartite genetic structure

Previously, we reported a shared genetic profile among EBA western Baikal hunter-gatherers (Baikal_EBA) and Late Bronze Age (LBA) pastoralists in northern Mongolia (Khövsgöl_LBA) (Jeong et al., 2018). This genetic profile, composed of major and minor ANA and ANE ancestry components, respectively, is also shared with the earlier eastern Baikal (Fofonovo_EN) and Mongolian (centralMongolia_preBA) groups analyzed in this study (Fig. 3a,b), suggesting a regional persistence of this genetic profile for nearly three millennia. Centered in northern Mongolia, this genetic profile is distinct from that of other Bronze Age groups. Overall, we find three distinct and geographically structured gene pools in LBA Mongolia, with the Khövsgöl_LBA population representing one of them (Fig. 3b). The other two, which we refer to as “Altai_MLBA” and “Ulaanzuuk_SlabGrave”, are described below.

During the MLBA (1900-900 BCE), as grasslands expanded in response to climate change, new pastoralist cultures expanded out of inner-montane regions and across the Eastern Steppe (Kindstedt and Ser-Od, 2019). This period is also notable for the first regional evidence horse milking (ca. 1200 BCE; (Wilkin et al., 2019)), and a dramatic intensification of horse use, including the emergence of mounted horseback riding that would have substantially extended the accessibility of remote regions of the steppe. Today, horse milking is exclusively associated with alcohol production (Bat-Oyun et al., 2015), and this first appearance of horse milk may mark the origin of this social tradition on the Eastern Steppe. In the Altai-Sayan region, dairy pastoralists associated with the Mönkhkhairkhan, Deer Stone-Khirigsuur Complex (DSKC), and Baitag cultures (Altai_MLBA, n=7), all show clear genetic evidence of admixture between a Khövsgöl_LBA-related ancestry and a Sintashta-related WSH ancestry (Fig. 3b). Overall, they form an “Altai_MLBA” cline on PCA between Western Steppe groups and the Baikal_EBA/Khövsgöl_LBA cluster (Fig. 2), with their position varying on PC1 according to their level of Western ancestry (Table S16).

This is the first appearance on the Eastern Steppe of a Sintashta-like ancestry (frequently referred to as “steppe_MLBA” in previous studies), which is distinct from prior Western ancestries present in the Afanasievo and Chemurchek populations, and which instead shows a close affinity to European Corded-Ware populations and later Andronovo-associated groups, such as the Sintashta (Allentoft et al., 2015). In Khovd province, individuals belonging to Mönkhkhairkhan and DSKC cultures (SBG001 and BER002, respectively) have a similar genetic profile that is best modeled as an equal mixture of Khövsgöl_LBA and Sintashta (Fig. 3b; Table S16). This genetic profile matches that previously described for a genetic outlier in northern Mongolia that deviated from the Khövsgöl_LBA cluster in a previous study (ARS026; (Jeong et al., 2018)). An additional four Altai_MLBA individuals belonging to Baitag (ULI003), DSKC (ULI001), and an unclassified culture (BIL001, ULZ001) also fit this admixture model with varying admixture proportions (Table S16). Taken together, the Altai_MLBA cline reveals the ongoing mixture of two source populations: a Sintashta/Andronovo-related WSH population and a local population represented by Khövsgöl_LBA. The admixture is estimated to have occurred only 9±2 generations (∼250 years) before the individuals analyzed in this study, a finding consistent with their heterogeneous ancestry proportions (Fig. S12 and S13). Because the Sintashta culture (ca. 2800-1600 BCE) is associated with novel transportation technologies, such as horse-drawn chariots (Anthony, 2010), the appearance of this ancestry profile on the Eastern Steppe suggests that heightened mobility capabilities played an important role in linking diverse populations across the Eurasian Steppe.

Three MLBA individuals in our dataset present genetic profiles that cannot be fully explained by the Altai_MLBA cline. The Altai Mönkhkhairkhan individual UAA001 matches with 3-way admixture models using Khövsgöl_LBA/Baikal_EBA as the eastern Eurasian source, Afanasievo as the WSH source (but not Sintashta), and adding Gonur1_BA as a third source (Table S16). Thus, a minor Iranian-related ancestry component is necessary to model this individual’s genetic profile, as reflected by his displacement along PC2 in Fig. 2. We also find that the culturally unclassified Altai individual KHI001 may need additional ancestry from Gonur1_BA to improve the existing 2-way admixture model. Last, UUS001, despite coming from a DSKC context in Khövsgöl province, fits better with a 3-way model of using eastMongolia_preBA and Gonur1_BA with Sintashta as the WSH source. Although cultural differences may have existed among the major MLBA mortuary traditions of the Altai and northern Mongolia (Mönkhkhairkhan, DSKC, and Baitag), they do not form distinct genetic groups. Rather, these mortuary traditions encompass individuals of disparate and recently admixed ancestry, and the proliferation and variability of mortuary features at this time likely reflects this demographic dynamism.

The populations making up the heterogeneous Altai_MLBA cline left descendants in the Altai-Sayan region, who we later identify at the Uyuk site of Chandman Mountain (“Chandman_IA”, ca. 400-200 BCE) in northwestern Mongolia during the Early Iron Age (EIA). Nine Chandman_IA individuals form a tight cluster on PCA at the end of the previous Altai_MLBA cline away from Khövsgöl_LBA cluster (Fig. 2). During the EIA, the Uyuk (also known as the Aldy-Bel or Sagly-Bazhy culture) were pastoralists largely centered in the Upper Yenisei region of present-day Tuva, and together with the Pazyryk, with whom they share log chamber tombs and other features, they formed part of a broader Saka cultural phenomenon that stretched across the Western Steppe, the Tarim Basin, and the Upper Yenisei (Parzinger, 2006). In addition to dairying, many Saka groups, including EIA individuals at the Chandman Mountain site, also cultivated millet (Murphy et al., 2013; Ventresca Miller and Makarewicz, 2019; Wilkin et al.), a practice that had been earlier adopted by many southern steppe and Central Asian populations during the westward spread of millet from China to the Caucasus during the second millennium BCE (Ventresca Miller and Makarewicz, 2019).

We find that EIA Saka populations systematically deviate from the earlier Altai_MLBA cline, requiring a third ancestral component (Fig. 3c). The appearance of this ancestry, related to populations of Central Asia (Caucasus/Iranian Plateau/Transoxiana regions) including BMAC (Narasimhan et al., 2019), is clearly detected in the Iron Age groups such as Central Saka (Damgaard et al., 2018), TianShan Saka (Damgaard et al., 2018), Tagar (Damgaard et al., 2018), and Chandman_IA (Tables S17-S18), while absent in the earlier DSKC and Karasuk groups (Tables S16-S17). This third component makes up 6-24% of the ancestry in these Iron Age groups, and the date of admixture in Chandman_IA is estimated ∼17±4 generations earlier, ca. 750 BCE, which postdates the collapse of the BMAC ca. 1600 BCE and slightly predates the formation of the Persian Achaemenid empire ca. 550 BCE (Fig. S12, S13). We suggest that this Iranian-related genetic influx was mediated by increased contact and mixture with agropastoralist populations in the region of Transoxiana (Turan) and Fergana during the LBA to EIA transition. The widespread emergence of horseback riding during the late second and early first millennium BCE (Drews, 2004), and the increasing sophistication of horse transport thereafter, likely contributed to increased population contact and the dissemination of this Iranian-related ancestry onto the steppe. Our results do not exclude additional spheres of contact, such as increased mobility along Inner Asian Mountain Corridor, which could have also introduced this ancestry into the Altai via Xinjiang starting in the Bronze Age (Frachetti, 2012).

In contrast to the MLBA and EIA cultures of the Altai and northern Mongolia, different burial traditions are found in the eastern and southern regions of Mongolia (Honeychurch, 2015), notably the LBA Ulaanzuukh (1450-1150 BCE) and EIA Slab Grave (1000-300 BCE) cultures. In contrast to other contemporaneous Eastern Steppe populations, we find that individuals associated with these burial types show a clear northeastern-Eurasian (ANA-related) genetic profile lacking both ANE and WSH admixture (Fig. 2; Fig. 3c; Fig. S7). Both groups were ruminant pastoralists, and the EIA Slab Grave culture also milked horses (Wilkin et al., 2019). The genetic profiles of Ulaanzuukh and Slab Grave individuals are genetically indistinguishable (Fig. 2 and Table S16), consistent with the archaeological hypothesis that the Slab Grave tradition emerged out of the LBA Ulaanzuukh (Honeychurch, 2015; Khatanbaatar, 2019). Both groups are also indistinguishable from the earlier eastMongolia_preBA individual dating to ca. 4600 BCE, suggesting a long-term (>4,000 year) stability of this prehistoric eastern Mongolian gene pool (Table S16). In subsequent analyses, we merged Ulaanzuukh and Slab Grave into a single genetic group (“Ulaanzuukh_SlabGrave”). The Ulaanzuukh_SlabGrave genetic cluster is the likely source of the previously described DSKC eastern outlier from Khövsgöl province (ARS017) (Jeong et al., 2018), as well as a culturally unclassified individual (TSI001) from central Mongolia who dates to the LBA-EIA transition (Figs. 2 and 3c; Table S16). In addition, the Mönkhkhairkhan individual KHU001 from northwest Mongolia has a non-negligible amount of Ulaanzuukh_SlabGrave ancestry in addition to his otherwise Baikal_EBA ancestry (Fig. 3c; Table S16). While these three individuals attest to occasional long-distance contacts between northwestern and eastern Mongolia during the LBA, we find no evidence of Ulaanzuukh_SlabGrave ancestry in the Altai, and the overall frequency of the Ulaanzuukh_SlabGrave genetic profile outside of eastern and southern Mongolia during the MLBA is very low. During the EIA, the Slab Grave culture expanded northwards, often disrupting and uprooting former DSKC graves in their path (Honeychurch, 2015), and it ultimately reached as far north as the eastern Baikal region, which is reflected in the genetic profile of the Slab Grave individual PTO001 in this study (Fig. 3c). Overall, our findings reveal a strong east-west genetic division among Bronze Age Eastern Steppe populations through the end of the Early Iron Age.

### The Xiongnu Empire, the rise of the first imperial steppe polity

Arising from the prehistoric populations of the Eastern Steppe, large-scale polities began to develop during the late first millennium BCE. The Xiongnu was the first historically-documented empire founded by pastoralists, and its establishment is considered a watershed event in the sociopolitical history of the Eastern Steppe (Brosseder and Miller, 2011; Honeychurch, 2015). The Xiongnu held political dominance in East and Central Asia from the third century BCE through the first century CE. The cultural, linguistic and genetic make-up of the people who constituted the Xiongnu empire has been of great academic interest, as has their relationship to other contemporaneous and subsequent nomadic groups on the Eastern Steppe. Here we report genome-wide data for 60 Xiongnu-era individuals from across Mongolia and dating from 350 BCE to 130 CE, thus spanning the entire period of the Xiongnu empire. Although most individuals date to the late Xiongnu period (after 50 BCE), 13 individuals predate 100 BCE and include 12 individuals from the northern early Xiongnu frontier sites of Salkhityn Am (SKT) and Atsyn Gol (AST) and one individual from the early Xiongnu site of Jargalantyn Am (JAG) in eastern Mongolia.

We observe two distinct demographic processes that contributed to the formation of the early Xiongnu. First, half of the individuals (n=6) form a genetic cluster (earlyXiongnu_west) resembling that of Chandman_IA of the preceding Uyuk culture from the Altai-Sayan region (Fig. 2). They derive 91.8±1.9% of their ancestry from Chandman_IA with the remainder attributed to additional Iranian-related ancestry, which we model using BMAC as a proxy (Fig. 3d; Table S19). This suggests that the low-level Iranian-related gene flow identified among the Chandman Mountain Uyuk during the EIA likely continued during the second half of the first millennium BCE, spreading across western and northern Mongolia. Second, six individuals (“earlyXiongnu_rest”) fall intermediate between the earlyXiongnu_west and Ulaazuukh_SlabGrave clusters; four carry varying degrees of earlyXiongnu_west (39-75%) and Ulaazuukh_SlabGrave (25-61%) related ancestry, and two (SKT004, JAG001) are indistinguishable from the Ulaanzuuk_SlabGrave cluster (Fig. 3d; Tables S19-S20). This genetic cline linking the earlyXiongnu_west and Ulaanzuukh_SlabGrave gene pools signifies the unification of two deeply diverged and distinct lineages on the Eastern Steppe - between the descendants of the DSKC, Mönkhkhairkhan, Baitag, and Uyuk cultures in the west and the descendants of the Ulaanzuuk and Slab Grave cultures in the east - and even tracing back before the Bronze Age, when genetic differentiation is seen even among the northern/western and eastern hunter-gatherers. Overall, the low-level influx of Iranian-related gene flow continuing from the previous Uyuk culture and the sudden appearance of a novel east-west mixture uniting the gene pools of the Eastern Steppe are the two defining demographic processes associated with the rise of the Xiongnu.

Among late Xiongnu individuals, we find even higher genetic heterogeneity (Fig. 2), and their distribution on PC indicates that the two demographic processes evident among the early Xiongnu continued into the late Xiongnu period, but with the addition of new waves and complex directions of gene flow. Of the 47 late Xiongnu individuals, half (n=26) can be adequately modelled by the same admixture processes seen among the early Xiongnu: 22 as a mixture of Chandman_IA+Ulaanzuuk_SlabGrave, two (NAI002, TUK002) as a mixture of either Chandman_IA+BMAC or Chandman_IA+Ulaanzuuk_SlabGrave+BMAC, and two (TUK003, TAK001) as a mixture of either earlyXiongnu_west+Ulaanzuukh_SlabGrave or earlyXiongnu_west+Khovsgol_LBA (Fig 3d). A further two individuals (TEV002, BUR001) also likely derive their ancestry from early Xiongnu gene pool, although the *p*-value of their models is slightly lower than the 0.05 threshold (Table S20). However, a further 11 late Xiongnu with the highest proportions of western Eurasian affinity along PC1 cannot be modelled using BMAC or any other ancient Iranian-related population. Instead, they fall on a cluster of ancient Sarmatians from various locations in the western and central steppe (Fig. 2).

Admixture modeling confirms the presence of a Sarmatian-related gene pool among the late Xiongnu: three individuals (UGU010, TMI001, BUR003) are indistinguishable from Sarmatian, two individuals (DUU001, BUR002) are admixed between Sarmatian and BMAC, three individuals (UGU005, UGU006, BRL002) are admixed between Sarmatian and Ulaanzuuk_SlabGrave, and three individuals (NAI001, BUR004, HUD001) require Sarmatian, BMAC and Ulaanzuuk_SlabGrave (Fig. 3d; Table S20. In addition, eight individuals with the highest eastern Eurasian affinity along PC1 are distinct from both the Ulaanzuuk_SlabGrave and Khövsgöl_LBA genetic profiles, showing affinity along PC2 towards present-day people from East Asia further to the south (Fig. 2). Six of these individuals (EME002, ATS001, BAM001, SON001, TUH001, YUR001) are adequately modeled as a mixture of Ulaanzuuk_SlabGrave and Han (Table S19-S20), and YUR001 in particular exhibits a close genetic similarity to two previously published Han empire soldiers (Damgaard et al., 2018), whose genetic profile we refer to as “Han_2000BP” (Table S20). The remaining two individuals (BRU001, TUH002) are similar but also require the addition of Sarmatian ancestry (Fig. 3d, Table S20). The late Xiongnu are thus characterized by two additional demographic processes that distinguish them from the early Xiongnu: gene flow from a new Sarmatian-related Western ancestry source and intensified interaction and mixture with people of the contemporaneous Han empire of China. Together, these results match well with historical records documenting the political influence that the Xiongnu exercised over their neighbors, including the Silk Road kingdoms of Central Asia and Han Dynasty China, as well as purported migrations both in and out of Mongolia (Miller, 2014). Overall, the Xiongnu period can be characterized as one of expansive and extensive gene flow that began by uniting the gene pools of western and eastern Mongolia and ended by uniting the gene pools of western and eastern Asia.

### Fluctuating genetic heterogeneity in the post-Xiongnu polities

After the collapse of the Xiongnu empire ca. 100 CE, a succession of nomadic pastoralist regimes rose and fell over the next several centuries across the politically fragmented Eastern Steppe: Xianbei (ca. 100-250 CE), Rouran (ca. 300-550 CE), Türkic (552-742 CE), and Uyghur (744-840 CE). Although our sample representation for the Early Medieval period is uneven, consisting of one unclassified individual dating to the Xianbei or Rouran period (TUK001), 8 individuals from Türkic mortuary contexts, and 13 individuals from Uyghur cemeteries, it is clear that these individuals have genetic profiles that differ from the preceding Xiongnu period, suggesting new sources of gene flow into Mongolia at this time that displace them along PC3 (Fig. 2). Individual TUK001 (250-383 cal. CE), whose burial was an intrusion into an earlier Xiongnu cemetery, has the highest western Eurasian affinity. This ancestry is distinct from that of the Sarmatians, and closer to ancient populations with BMAC/Iranian-related ancestry (Fig. 2). Among the individuals with the highest eastern Eurasian affinity, two Türkic- and one Uyghur-period individual (ZAA004, ZAA002, OLN001.B) are indistinguishable from the Ulaanzuuk_SlabGrave cluster. Another individual (TUM001), who was recovered from the tomb ramp of an elite Türkic-era emissary of the Tang Dynasty, has a high proportion of Han-(78.1±1.5%) (Fig. 3e) and especially Han_2000BP-related ancestry (84±1.5%) (Table S21). This male, buried with two dogs, was likely a Chinese attendant sacrificed to guard the tomb entrance (Ochir et al., 2013). The remaining 17 Türkic and Uyghur individuals show intermediate genetic profiles (Fig. 3e).

The high genetic heterogeneity of the Early Medieval period is vividly exemplified by twelve individuals from the Uyghur period cemetery of Olon Dov (OLN; Fig. 2), in the vicinity of the Uyghur capital of Ordu-Baliq. Six of these individuals came from a single tomb (grave 19), of whom only two are related (OLN002 and OLN003, second-degree; Table S7); the absence of closer kinship ties raises questions about the function of such tombs and the social relationships of those buried within them. Most Uyghur period individuals exhibit a high, but variable, degree of west Eurasian ancestry - best modeled as a mixture of Alans (a historic nomadic pastoral group likely descended from the Sarmatians and contemporaries of the Huns (Bachrach, 1973)) and an Iranian-related (BMAC-related) ancestry - together with Ulaanzuukh_SlabGrave (ANA-related) ancestry (Fig. 3e). The admixture dates estimated for the ancient Türkic and Uyghur individuals in this study correspond to ca. 500 CE: 8±2 generations before the Türkic individuals and 12±2 generations before the Uyghur individuals (represented by ZAA001 and Olon Dov individuals).

### Rise of the Mongol empire

After the fall of the Uyghur empire in the mid-ninth century, the Khitans of northeast China established the powerful Liao Dynasty in 916 CE. Although few Khitan sites are known within Mongolia, the Khitans controlled large areas of the Eastern Steppe and are recorded to have relocated people within their conquered territories (Kradin and Ivliev, 2008). Our study includes three Khitan individuals (ZAA003, ZAA005, ULA001) from Bulgan province, all of whom have a strongly eastern Eurasian (ANA and Han_2000BP-related) genetic profile (Fig. 2), with <10% west Eurasian ancestry (Fig. 3f; Table S22). This may reflect the northeastern Asian origin of the Mongolic-speaking Khitan, but a larger sample size is required to adequately characterize the genetic profile of Khitan populations within Mongolia. In 1125 CE, the Khitan empire fell to the Jurchen’s Jin Dynasty, which was then conquered in turn by the Mongols in 1234 CE.

At its greatest extent, the Mongol empire (1206-1368 CE) spanned nearly two thirds of the Eurasian continent. It was the world’s largest contiguous land empire, and the cosmopolitan entity comprised diverse populations that flowed into the steppe heartland. We analyzed 62 Mongol-era individuals whose burials are consistent with those of low-level, local elites. No royal or regional elite burials were included, nor were individuals from the cosmopolitan capital of Karakorum. Although we find that Mongol-era individuals were diverse, they exhibit a much lower genetic heterogeneity compared to Xiongnu-era individuals (Fig. 2), and they almost entirely lack the residual ANE-related ancestry (in the form of Chandman_IA and Khövsgöl_LBA) that had been present among the Xiongnu and earlier northern/western MLBA cultures. On average, Mongol period individuals have a much higher eastern Eurasian affinity than previous empires, and this period marks the beginning of the formation of the modern Mongolian gene pool. We find that most historic Mongols are well-fitted by a three-way admixture model with the following ancestry proxies: Ulaanzuuk_SlabGrave, Han, and Alans. Consistent with their PCA location (Fig. 2), Mongol era individuals as a group can be modeled with only 15-18% of western Steppe ancestry (Alan or Sarmatian), but require 55-64% of Ulaanzuuk_SlabGrave and 21-27% of Han-related ancestry (Table S22). Applying the same model to each individual separately, this three-source model adequately explains 56 out of 61 ancient Mongols (based on *p*-value at threshold of 0.05), as well as one unclassified Late Medieval individual dating to around the beginning of the Mongol empire (SHU002) (Table S23).

Since the fall of the Mongol empire in 1368 CE, the genetic profile of the Mongolian populations has not substantially changed. The genetic structure established during the Mongol empire continues to characterize present-day Mongolic-speaking populations living in both Mongolia and Russia. We examined the genetic cladality between the historic Mongols and seven present-day Mongolic-speaking groups (Mongols, Kalmyk, Buryat, Khamnegan, Daur, Tu and Mongola) using an individual-based qpWave analysis {Sample; Mongolic group}. Within the resolution of current data, 34/61 historic Mongols are genetically cladal with at least one modern Mongolic-speaking population (Fig. S15). In addition, nearly a third of historic Mongol males (12/38) have Y haplogroup C2b, which is also widespread among modern Mongolians (Table S6, Fig. S3); C2b is the presumed patrilineage of Genghis Khan (Zerjal et al., 2003). The Mongol empire had a profound impact on restructuring the political and genetic landscape of the Eastern Steppe, and these effects endured long after the decline of the empire and are still evident in Mongolia today.

### Functional and gendered aspects of recurrent admixture in the Eastern Steppe

To investigate the functional aspects of recurrent admixture on the Eastern Steppe, we estimated the population allele frequency of five SNPs associated with functional or evolutionary aspects of lactose digestion (*LCT/MCM6*), dental morphology (*EDAR*), pigmentation (*OCA2, SLC24A5*), and alcohol metabolism (*ADH1B*) (Fig. 4a). First, we find that despite a pastoralist lifestyle with widespread direct evidence for milk consumption (Jeong et al., 2018; Wilkin et al., 2019), the MLBA and EIA individuals of the Eastern Steppe did not have any derived mutations conferring lactase persistence (LP). Individuals from subsequent periods did have the derived mutation that is today widespread in Europe (rs4988235), but at negligibly low frequency (∼5%) and with no increase in frequency over time (Fig. 4a). This is somewhat remarkable given that, in addition to other dairy products, contemporary Mongolian herders consume up to 4-10 liters of *airag* (fermented mare’s milk, ∼2.5% lactose) per day during the summer months (Bat-Oyun et al., 2015), resulting in a daily intake of 100-250g of lactose sugar. Petroglyph depictions of airag production date back to the EIA in the Yenisei Basin (Dėvlet, 1976), and accounts of the historic Mongols record abundant and frequent consumption of airag, as well as a wide range of additional liquid and solid ruminant dairy products (Bayarsaikhan, 2016; Onon, 2005), which has been additionally confirmed by ancient proteomic evidence (Jeong et al., 2018; Wilkin et al., 2019). How Mongolians have been able to digest such large quantities of lactose for millennia in the absence of LP is unknown, but it may be related to their reportedly unusual gut microbiome structure, which today is highly enriched in lactose-digesting *Bifidobacteria* spp. (Liu et al., 2016).

**Fig. 4.**
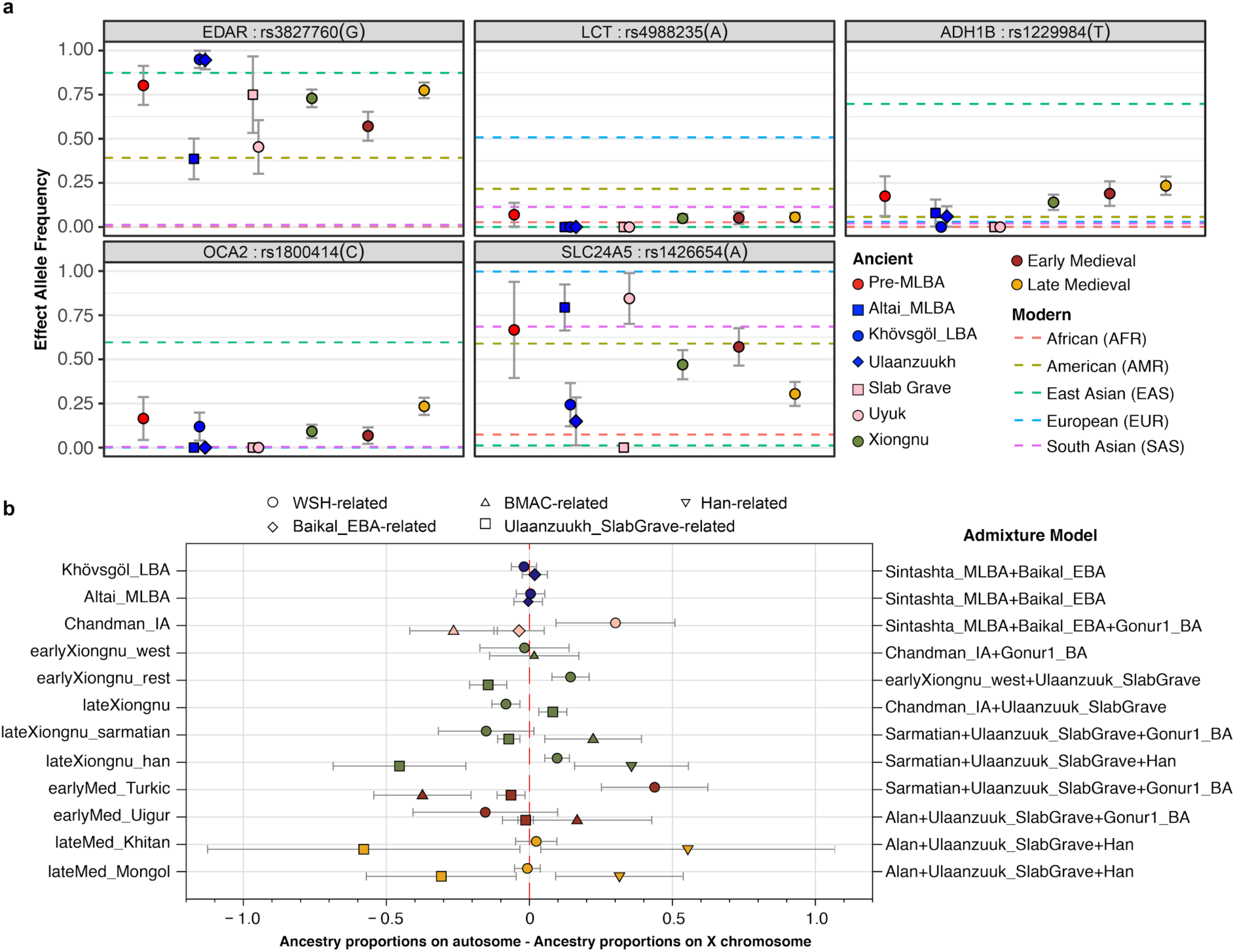
Functional allele frequencies and sex-biased patterns of genetic admixture. (**a**) Allele frequencies of five phenotypic SNP changes through time. For the effective allele, we show maximum likelihood frequency estimates and one standard error bars for each ancient group. The pre-MLBA category corresponds to the sum of all ancient groups before Mönkhkhairkhan. Xiongnu, early Medieval, late Medieval correspond to the sum of all ancient groups in each period correspondingly. Horizontal dashed lines show allele frequency information from the 1000 Genomes Project five super populations. (**b**) Sex-biased patterns of genetic admixture by period and population. We calculated Z-scores for every ancient individual who has genetic admixture with WSH-/Iranian-/Han-related ancestry. Positive scores suggest more WSH-/Iranian-/Han-related ancestry on the autosomes, i.e., male-driven admixture. See Fig. S6 for individual Z scores.

Genetic markers that underwent regional selective sweeps show allele frequency changes that correlate with changes in the genome-wide ancestry profile (Fig. 4a). For example, rs3827760 in *EDAR* (ectodysplasin A receptor) and rs1426654 in *SLC24A5* (solute carrier family 24 member 5) are well-known targets of positive selection in East Asians and Western Eurasians, respectively (Sabeti et al., 2007). Our MLBA and EIA populations show a strong population differentiation in the allele frequencies of these two SNPs: rs3827760 frequency is much higher in groups with higher eastern Eurasian affinity (Khovsgol_LBA, Ulaanzuuk_SlabGrave) while rs1426654 is higher in Altai_MLBA and Chandman_IA (Table S8). We find that two SNPs that have undergone more recent positive selection (Donnelly et al., 2012; Li et al., 2011) among East Asians, rs1229984 in *ADH1B* (aldehyde dehydrogenase 1B) and rs1800414 in *OCA2* (oculocutaneous albinism II), were absent or in extremely low frequency during the MLBA and EIA, when the eastern Eurasian ancestry was primarily ANA-related, but increased in frequency over time as the proportion of East Asian ancestry increased through interactions with China and other groups (Table S8).

Finally, we investigated gendered dimensions of the population history of the Eastern Steppe. Sex-biased patterns of genetic admixture can be informative about gendered aspects of migration, social kinship, and family structure. We observed an especially strong male-biased admixture from WSH groups during the EIA and Early Medieval periods. To estimate this sex-biased admixture, we compared the ancestry proportions in the autosomes and X chromosome for each component, and calculated the statistical significance of their difference following a published strategy (Mathieson et al., 2018). When the proportion of an ancestry is higher in the autosomes than in the X-chromosome, the admixture process involves male ancestors harboring this ancestry in higher proportion than female ancestors. As shown in Fig. 4b, we observe a clear signal of male-biased WSH admixture among the EIA Uyuk and during the Türkic period (i.e., more positive Z scores), which also corresponds to the decline in the Y chromosome lineage Q1a and the concomitant rise of the western Eurasian lineages such as R and J (Fig. S3). During the later Khitan and Mongol empires, sex-bias Z scores shift to indicate a prominant male bias for East Asian-related ancestry (Fig. S6), which can also be seen from the rise in frequency of Y-chromosomal lineage O2a (Fig. S3). The Xiongnu period exhibits the most complex pattern of male-biased admixture, whereby different genetic subsets of the population exhibit evidence of different sources of male-biased admixture (Fig. S6). We also detect ten genetic relative pairs among the Xiongnu individuals in this study, including a father-daughter pair buried in the same grave (JAG001 and JAA001) at Jargalantyn Am, as well as a mother-son pair (IMA002 and IMA005) at Il’movaya Pad, a brother-sister pair (TMI001 and BUR003) at Tamiryn Ulaan Khoshuu, and a brother-brother pair (SKT002 and SKT006) at Salkhityn Am (Table S7). Of the remaining six pairs, three are female-female relative pairs buried within the same site, suggesting the presence of extended female kinship within Xiongnu groups. These relationships, when combined with mortuary features, offer the first clues to local lineage and kinship structures within the Xiongnu empire, which are otherwise poorly understood.

## Discussion

The population history of the Eastern Steppe is one marked by the repeated mixing of diverse eastern and western Eurasian gene pools. However, rather than simple waves of migration, demographic events on the Eastern Steppe have been complex and variable. Generating more than 200 genome-wide ancient datasets, we have presented the first genetic evidence of this dynamic population history, from ca. 4600 BCE through the end of the Mongol empire. We found that the Eastern Steppe was sparsely populated by hunter gatherers of ANA and ANE ancestry during the mid-Holocene, and then shifted to a dairy pastoralist economy during the Bronze Age. Migrating Yamnaya/Afanasievo steppe herders, equipped with carts and domestic livestock (Kovalev and Erdenebaatar, 2009), appear to have first introduced ruminant dairy pastoralism ca. 3000 BCE (Wilkin et al., 2019), but surprisingly had little lasting genetic impact, unlike in Europe (Allentoft et al., 2015; Haak et al., 2015; Mathieson et al., 2015). Subsequent Chemurchek pastoralists in the Altai were confirmed in this study to represent a separate migration of dairy pastoralists with ANE and Iranian-related ancestry, who possibly migrated into the Altai region from the south, likely via Xinjiang and/or mountainous Central Asia due to the concentration of Chemurchek burials in this region (Jia and Betts, 2010). By the MLBA, ruminant dairy pastoralism had been adopted by populations throughout the Eastern Steppe (Wilkin et al., 2019), regardless of ancestry, and this subsistence has continued, with the additions of horse milking in the LBA and camel milking in the Mongol period (Wilkin et al., 2019), to the present day (Bat-Oyun et al., 2015; Kindstedt and Ser-Od, 2019). Puzzlingly, however, there is no evidence of selection for lactase persistence over this 5,000-year history, despite the repeated introduction of this genetic trait by subsequent migrations of groups from the west. This suggests a different trajectory of lactose adaptation in Asia that to date remains unexplained.

During the MLBA, we observed the formation of a tripartite genetic structure on the Eastern Steppe, characterized by the continuation of pre-Bronze ANA ancestry in the east and a cline of genetic variation between pre-Bronze Age ANA-ANE ancestry in the north and increasing proportions of a new Sintashta-related WSH ancestry in the west. The Sintashta, a western forest steppe culture with genetic links to the European Corded Ware cultures (Mathieson et al., 2015), were masters of bronze metallurgy and chariotry (Anthony, 2010), and the appearance of this ancestry on the Eastern Steppe may be linked to the introduction of new (especially horse-related) technologies. DSKC sites in particular show widespread evidence for horse use in transport, and perhaps even riding (Taylor et al. 2015), and genetic analysis has demonstrated a close link between these animals and the Sintashta chariot horses (Fages et al., 2019). The strong east-west genetic division among Bronze Age Eastern Steppe populations at this time was maintained for more than a millennium and through the end of the EIA, when the first clear evidence for widespread horseback riding appears (Drews, 2004) and the heightened mobility of some groups, notably the eastern Slab Grave culture (Honeychurch, 2015), began to disrupt this structure. Eventually, the three major ancestries met and mixed, and this was contemporaneous with the emergence of the Xiongnu empire. The Xiongnu are characterized by extreme levels of genetic heterogeneity and increased diversity as new and additional ancestries from China, Central Asia and the Western Steppe (Sarmatian-related) rapidly entered the gene pool.

Genetic data for the subsequent early medieval period are relatively sparse and uneven, and few Xianbei or Rouran sites have yet been identified during the 400-year gap between the Xiongnu and Türkic periods. We observed high genetic heterogeneity and diversity during the Türkic and Uyghur periods, with slight shifts in their Sarmatian-like western ancestry component towards that seen among the contemporaneous Alans, a Sarmatian-descendent group known for invading the Roman empire and warring with the Germanic tribes of Europe (Bachrach, 1973). Following the collapse of the Uyghur empire, we documented a final major genetic shift during the late medieval period towards greater eastern Eurasian ancestry, which is consistent with historically documented expansions of Tungusic- (Jurchen) and Mongolic-(Khitan and Mongol) speaking groups from the northeast into the Eastern Steppe (Biran, 2012). We also observed that this East Asian-related ancestry was brought into the Late Medieval populations more by male than female ancestors, and we observed a corresponding increase in Y-haplogroups associated with southeast Asians and the purported patriline of Ghenghis Khan (O2a and C2b, respectively). By the end of the Mongol period the genetic make-up of the Eastern Steppe had dramatically changed, retaining little of the ANE ancestry that had been a prominent feature during its prehistory. Today, ANE ancestry survives in appreciable amounts only in isolated Siberian groups and among the indigenous peoples of the Americas (Jeong et al., 2019). The genetic profile of the historic Mongols is still reflected among contemporary Mongolians, suggesting a relative stability of this gene pool over the last ∼700 years.

Having documented key periods of genetic shifts in the Eastern steppe, future work may be able to explore whether these shifts are also linked to cultural and technological innovations and how these innovations may have influenced the political landscape. Integrating these findings with research on changes in horse technology and herding practices, as well as shifts in livestock traits and breeds, may prove particularly illuminating. This study represents the first large-scale paleogenomic investigation of the eastern Eurasian Steppe and it sheds light on the remarkably complex and dynamic genetic diversity of the region. Despite this progress, there is still a great need for further genetic research in central and eastern Eurasia, and particularly in northeastern China, the Tarim Basin, and the eastern Kazakh steppe, in order to fully reveal the population history of the Eurasian Steppe and its pivotal role in world prehistory.

## Acknowledgments

We thank D. Navaan, Z. Batsaikhan, M. Erdene, D. Tumen, S. Ulziibayar, D. Khatanbaatar, D. Erdenebaatar, U. Erdenebat, A. Ochir, G. Ankhsanaa, A. Kovalev, V. Volkow, D. Tseveendorj, L. Erdenebold, M. Horton, Sh. Uranchimeg, Ch. Vanchigdash, B. Naran, B. Ochir, N. Ser-Odjav, P. Konovalov, Kh. Lhagvasuren, N. Mamonova, E. Mijiddorj, Ch. Munkhbayar, Namsrainaidan, A. Okladnikov, Kh. Perlee, S. Danilov, and Burayev for contributing archaeological material to the study. We thank R. Flad for comments on early manuscript drafts, M. Bleasdale and S. Gankhuyg for assistance with sampling, S. Nagel and M. Meyer for assistance with ssDNA library preparation, S.Nakagome for sharing a script to estimate population allele frequency from low-coverage sequence data, and P. Moorjani for granting access to the DATES program prior to publication.

## Funding

This research was supported by the Max Planck Society, the U.S. National Science Foundation (BCS-1523264 to C.W.), the Deutsche Forschungsgemeinschaft (SFB 1167 No. 257731206 to J.B.), the Ministry of Education and Science of the Russian Federation (Grant #14.W03.31.0016 to N.K., B.B., D.M., and P.K.), and the European Research Council under the European Union’s Horizon 2020 research and innovation programme under grant agreement numbers 771234-PALEORIDER (W.H.) and 804884-DAIRYCULTURES (C.W.).

## Author contributions

C.W., C.J., E.M., and N.B. designed the research; C.W. and C.J. supervised the research; E.M., S.W., J.H., J.B., S.U., W.H., N.K., B.A.B., D.A.M, P.B.K., E.Z., A.V.M., N.B., and C.W. provided materials and resources; B.M., W.T.T.T., J.B., T.T., and E.M. performed archaeological data analysis; R.S., and C.C.. performed genetic laboratory work; K.W., C.J., F.K., and S.S. performed genetic data analysis; B.M., K.W., W.T.T.T, C.J., and C.W. integrated the archaeological and genetic data; K.W., C.J., and C.W. wrote the paper, with contributions from B.M., W.T.T.T., J.B., and the other coauthors.

## Declaration of interests

The authors declare no competing interests.

## STAR Methods Text

### LEAD CONTACT AND MATERIALS AVAILABILITY

Further information and requests for resources and reagents should be directed to and will be fulfilled by the Lead Contact, Christina Warinner.

### EXPERIMENTAL MODEL AND SUBJECT DETAILS

Here we present new genome-wide data for 213 ancient individuals from Mongolia and 13 individuals from Buryatia, Russia, which we analyze together with 21 previously published ancient Mongolian individuals (Jeong et al., 2018), for a total of 247 individuals. All new Mongolian individuals, except ERM, were sampled from the physical anthropology collections at the National University of Mongolia and the Institute for Archaeology and Ethnology in Ulaanbaatar, Mongolia. ERM001/002/003 was provided by Jan Bemmann. Russian samples were collected from the Institute for Mongolian, Buddhist, and Tibetan Research as well as the Buryat Scientific Center, Russian Academy of Sciences (RAS).

Together, this ancient Eastern Steppe dataset of 247 individuals originates from 89 archaeological sites (Fig. 1; Table S1) and spans approximately 6,000 years of time (Data Tables S1, S2, S6). High quality genetic data was successfully generated for 214 individuals and was used for population genetic analysis (Table S4). Subsistence information inferred from proteomic analysis of dental calculus has been recently published for a subset of these individuals (n=32; (Wilkin et al., 2019)), and stable isotope analysis of bone collagen and enamel (n=137) is also in progress (Wilkin et al.); together, these data allow direct comparison between the biological ancestry of specific archaeological cultures and their diets, particularly with respect to their dairy and millet consumption.

### METHOD DETAILS

#### Radiocarbon dating of sample materials

A total of 24 new radiocarbon dates were obtained by accelerator mass spectrometry (AMS) of bone and tooth material at the Curt-Engelhorn-Zentrum Archäometrie (CEZA) in Mannheim, Germany (n=22) and the University of Cologne Centre for Accelerator Mass Spectrometry (CologneAMS) (n=2). Selection for radiocarbon dating was made for all burials with ambiguous or unusual burial context and for all individuals appearing as genetic outliers for their assigned period. Uncalibrated direct carbon dates were successfully obtained for all bone and tooth samples (Table S12). An additional 81 previously published radiocarbon dates for individuals in this study were also compiled and analyzed, making the total number of directly dated individuals in this study 105. Dates were calibrated using OxCal v.4.3.2 (Ramsey, 2017) with the r:5 IntCal13 atmospheric curve (Reimer et al., 2013).

Of the 105 total radiocarbon dates analyzed in this study, 27 conflicted with archaeological period designations reported in excavation field notes or previous publications (Table S12). Four burials of uncertain cultural context were successfully assigned to the Middle/Late Bronze Age (BIL001, MIT001) and Late Medieval periods (UUS002, ZAA003). One burial originally assigned to the Late Medieval period was reassigned to the pre-Bronze Age following radiocarbon dating (ERM001), and three burials originally assigned to the Middle/Late Bronze Age were similarly reassigned to the Early Bronze Age (KUM001, KUR001, IAG001). This suggests that early burials may be underreported in the literature because they are mistaken for later graves. Likewise three burials originally classified as Late Medieval were found to be hundreds or thousands of years older, dating to the Early Medieval (TSB001) and Middle/Late Bronze Age (ULZ001, TSI001) periods. Although some highly differentiated burial forms can be characteristic of specific locations and time periods, simple burial mounds also exist for all periods and - lacking distinctive features - they can be difficult if not impossible to date without radiometric assistance.

In addition to early burials being mistaken for later ones, late burials were also misassigned to earlier periods. For example, three burials originally assigned to the Middle/Late Bronze Age were determined to date to the Early Iron Age (DAR001), Xiongnu (ULN004), and Late Medieval (SHU001) periods, and two Early Iron Age (CHN010, CHN014), six Xiongnu (TUK001, UGU001, DUU002, BRL001, BAU001, DEE001), and two Early Medieval (ULA001, ZAA005) graves were likewise reassigned to later periods following radiocarbon dating. Part of the difficulty in correctly assigning archaeological period to later burials relates to the frequent reuse of earlier graves and cemeteries by populations from later periods. The site reports of several Xiongnu excavations noted burial intrusions, displaced burials, and other indications of burial disturbance and reuse. However, evidence of burial reuse may also be subtle and easily overlooked. As such, we recommend great care in making cultural or temporal assignments at multi-period cemeteries or for any burials showing evidence of disturbance.

#### Sampling for ancient DNA recovery and sequencing

Sampling was performed on a total of 169 teeth and 75 petrosal bones from fragmented crania originating from 225 individuals (Table S3). For 14 individuals, both a tooth and a petrosal bone were sampled (Table S5). For three individuals, two teeth were sampled, and for one individual, two teeth and one petrosal bone were sampled (Table S5). For Mongolian material, whole teeth and petrosal bone (except ERM) were collected at the physical anthropology collections of the National University of Mongolia and the Institute for Archaeology and Ethnology under the guidance and supervision of Erdene Myagmar and S. Ulziibayar. Petrosal and tooth material from ERM were provided by Jan Bemmann. For Russian material, whole teeth alongside petrosal bone or bone were collected from the Institute for Mongolian, Buddhist, and Tibetan Research as well as the Buryat Scientific Center, Russian Academy of Sciences (RAS). After collection, the selected human skeletal material was transferred to the Max Planck Institute for the Science of Human History (MPI-SHH) for genetic analysis.

#### Laboratory procedures for genetic data generation

Genomic DNA extraction and Illumina double-stranded DNA (dsDNA) sequencing library preparation were performed for all samples in a dedicated ancient DNA clean room facility at the MPI-SHH, following published protocols (Dabney et al., 2013) with slight modifications (Mann et al., 2018). We applied a partial treatment of the Uracil-DNA-glycosylase (UDG) enzyme to confine DNA damage to the ends of ancient DNA molecules (Rohland et al., 2015). Such “UDG-half” libraries allow us to minimize errors in the aligned genetic sequence data while also maintaining terminal DNA misincorporation patterns needed for DNA damage-based authentication. Library preparation included double indexing by adding unique 8-mer index sequences at both P5 and P7 Illumina adapters. After shallow shotgun sequencing for screening, we enriched libraries of 195 individuals with ≥ 0.1% reads mapped on the human reference genome (hs37d5; GRCh37 with decoy sequences) for approximately 1.24 million informative nuclear SNPs (“1240K”) by performing an in-solution capture using oligonucleotide probes matching for the target sites (Mathieson et al., 2015). In addition, eight samples (see Table S5; Table S4) were also selected and built into single-stranded DNA (ssDNA) sequencing libraries for comparison. Single-end 75 base pair (bp) or paired-end 50 bp sequences were generated for all shotgun and captured libraries on the Illumina HiSeq 4000 platform following manufacturer protocols. Output reads were demultiplexed by allowing one mismatch in each of the two 8-mer indices.

### QUANTIFICATION AND STATISTICAL ANALYSIS

#### Sequence data processing

Short read sequencing data were processed by an automated workflow using the EAGER v1.92.55 program (Peltzer et al., 2016). Specifically, in EAGER, Illumina adapter sequences were trimmed from sequencing data and overlapping sequence pairs were merged using AdapterRemoval v2.2.0 (Schubert et al., 2016). Adapter-trimmed and merged reads with 30 or more bases were then aligned to the human reference genome with decoy sequences (hs37d5) using BWA aln/samse v0.7.12 (Li and Durbin, 2009). A non-default parameter “-n 0.01” was applied. PCR duplicates were removed using dedup v0.12.2 (Peltzer et al., 2016). Based on the patterns of DNA misincorporation, we masked the first and last two bases of each read for UDG-half libraries and 10 bases for non-UDG single-stranded libraries, using the trimbam function in bamUtils v1.0.13 (Jun et al., 2015), to remove deamination-based 5’ C>T and 3’ G>A misincorporations. Then, we generated pileup data using samtools mpileup module (Li and Durbin, 2009), using bases with Phred-scale quality score ≥ 30 (“-Q30”) on reads with Phred-scale mapping quality score ≥ 30 (“-q30”) from the original and the end-masked BAM files. Finally, we randomly chose one base from pileup for SNPs in the 1240K capture panel for downstream population genetic analysis using the pileupCaller program v1.2.2 (https://github.com/stschiff/sequenceTools). For C/T and G/A SNPs, we used end-masked BAM files, and for the others we used the original unmasked BAM files. For the eight ssDNA libraries, we used end-masked BAM files for C/T SNPs, and the original BAM files for the others.

In cases where more than one sample was genetically analyzed per individual, we compared the amount of human DNA between samples. For pairs of petrous bone and teeth, human DNA was higher in the petrous bone in 8 of 13 individuals, and higher in the teeth of 5 of 13 individuals (Table S5). In addition, intra-individual sample variation was high, as evidenced by the high variance observed between paired tooth samples (Table S5). Finally, in a comparison of dsDNA and ssDNA libraries, ssDNA libraries yielded a higher endogenous content in 7 of 8 library pairs. All data from paired samples were merged prior to further analysis.

Of the 225 new individuals analyzed, 18 failed to yield sufficient human DNA (>0.1%) on shotgun screening (Table S4) and a further 6 individuals failed to yield at least 10,000 SNPs after DNA capture (Table S4). Both were excluded from downstream population genetic analysis.

#### Data quality authentication

To confirm that our sequence data consist of endogenous genomic DNA from ancient individuals with minimal contamination, we collected multiple data quality statistics. First, we tabulated 5’ C>T and 3’ G>A misincorporation rate (Fig. S2) as a function of position on the read using mapDamage v2.0.6 (Jónsson et al., 2013). Such misincorporation patterns, enriched at the ends due to cytosine deamination in degraded DNA, are considered as a signature of the presence of ancient DNA in large quantities (Sawyer et al., 2012). Second, we estimated mitochondrial DNA contamination for all individuals using the Schmutzi program (Renaud et al., 2015). Specifically, we mapped adapter-removed reads to the revised Cambridge Reference Sequence of the human mitochondrial genome (rCRS; NC_012920.1), with an extension of 500 bp at the end to preserve reads passing through the origin. We then wrapped the alignment to the circular reference genome using circularmapper v1.1 (Peltzer et al., 2016). The contDeam and schmutzi modules of the Schmutzi program were successively run with the world-wide allele frequency database from 197 individuals, resulting in estimated mitochondrial DNA contamination rates for each individual (Table S4). Last, for males, we also estimated the nuclear contamination rate (Table S4) based on X chromosome data using the contamination module in ANGSD v0.910 (Korneliussen et al., 2014). For this analysis, an increased mismatch rate in known SNPs compared to that in the flanking bases is interpreted as the evidence of contamination because males only have a single copy of the X chromosome and thus their X chromosome sequence should not contain polymorphisms. We report the Method of Moments estimates using the “method 1 and new likelihood estimate”, but all the other estimates provide qualitatively similar results.

Ten individuals were estimated to have >5% DNA contamination (mitochondrial or X) or uncertain genetic sex (Table S4); these individuals were excluded from downstream population genetic analysis.

#### Genetic sex typing

We calculated the genetic sequence coverage on the autosomes and on each sex chromosome in order to obtain the ratio between the sex chomosome coverage and the autosome coverage. For 1240K capture data, we observe females to have an approximately even ratio of X to autosomal coverage (X-ratio of ∼0.8) and a Y-ratio of 0, and males to have approximately half the coverage on X and Y as autosomes (∼0.4). Genetic sex could be determined for a total of 224 individuals, of which 100 were female and 124 were male (Table S6).

#### Uniparental haplogroup assignment

We called mitochondrial consensus sequence from the Schmutzi output using the log2fasta program in the Schmutzi package, with quality threshold of 10. We then assigned each consensus sequence into a haplogroup (Table S6) using the HaploGrep 2 v2.1.19 (Weissensteiner et al., 2016). For the Y haplogroup assignment, we took 13,508 Y chromosome SNPs listed in the ISOGG database and made a majority haploid genotype call for each male using pileupCaller (with “-m MajorityCalling” option). We assigned each individual into a haplogroup (Table S6) using a patched version of the yHaplo program (Poznik, 2016) downloaded from https://github.com/alexhbnr/yhaplo. This version takes into account high missing rate of aDNA data to prevent the program from stopping its root-to-tip haplogroup search prematurely at an internal branch due to missing SNP and therefore assigning a wrong haplogroup. We used “--ancStopThresh 10” following the developer’s recommendation. Haplogroup assignments are shown in Fig. S3.

#### Estimation of genetic relatedness

To evaluate the relatedness within our dataset, we calculated pairwise mismatch rate of haploid genotypes on automosomes across all individuals. The pairwise mismatch rate for each pair of individuals, is defined as the number of sites where two individuals have different alleles sampled divided by the total number of sites that both individuals have data. The pairwise mismatch rate between unrelated individuals is set as the baseline and the coefficient of relationship is inversely linear to the baseline pairwise mismatch rate. More detailed description can be found in the Supplemental Materials of (Jeong et al., 2018).

A total of 15 first or second degree genetic relationships were observed across the dataset (Table S7), of which 10 date to the Xiongnu era. Additionally, in one case, a tooth and petrosal bone thought to belong to one individual (AT-871) were later discovered to belong to two different individuals (OLN001.A and OLN001.B). In another case, two teeth (AT-728 and AT-729) thought to belong to different individuals were found to originate from the same individual (TUK001/TAV008).

#### Data filtering and compilation for population genetic analysis

To analyse our dataset in the context of known ancient and modern genetic diversity, we merged it with previous published modern genomic data from i) 225 worldwide populations genotyped on the Human Origins array (Jeong et al., 2019; Lazaridis et al., 2014), ii) 300 high-coverage genomes in the Simons Genome Diversity Project (“SGDP”) (Mallick et al., 2016), and currently available ancient genomic data across Eurasian continent (Allentoft et al., 2015; de Barros Damgaard et al., 2018; Damgaard et al., 2018; Fu et al., 2014, 2016; Haak et al., 2015; Haber et al., 2017; Harney et al., 2018; Jeong et al., 2016, 2018; Jones et al., 2015; Kılınç et al., 2016; Lazaridis et al., 2016, 2017; Mathieson et al., 2015, 2018; McColl et al., 2018; Narasimhan et al., 2019; Raghavan et al., 2014, 2015; Rasmussen et al., 2010, 2014, 2015; Sikora et al., 2019; Unterländer et al., 2017; Yang et al., 2017). We obtained 1,233,013 SNP sites (1,150,639 of which on autosomes) across our dataset when intersecting with the SGDP dataset, and 597,573 sites (593,124 of which on autosomes) when intersecting with the Human Origins array.

#### Analysis of population structure and relationships

We performed principal component analysis (PCA) on the merged dataset with the Human Origins data using the smartpca v16000 in the Eigensoft v7.2.1 package (Patterson et al., 2006). Modern individuals were used for calculating PCs (Fig. S4), and ancient individuals were projected onto the pre-calculated components using “lsqproject: YES” option (Fig. 2; Fig. S5). To characterize population structure further, we also calculated *f3* and *f4* statistics using qp3Pop v435 and qpDstat v755 in the admixtools v5.1 package (Patterson et al., 2012). We added “*f4mode: YES*” option to the parameter file for calculating *f4* statistics. We used DATES (Narasimhan et al., 2019) for dating admixture in the different ancient population groups.

#### Admixture modeling using qpAdm

For modelling admixture and estimating ancestry proportions, we applied qpWave v410 and qpAdm v810 in the the admixtools v5.1 package (Patterson et al., 2012) on the merged dataset with the SGDP data to maximise resolution. To model the target as a mixture of the other source populations, qpAdm utilizes the linearity of *f4* statistics, i.e. one can find a linear combination of the sources that is symmetrically related to the target in terms of their relationship to all outgroups in the analysis. qpAdm optimizes the admixture coefficients to match the observed f4 statistics matrix, and reports a *p*-value for the null hypothesis that the target derives their ancestry from the chosen sources that are differently related to the outgroups (i.e., when *p*<0.05, the null hypothesis is rejected so that the target is different from the admixture of chosen sources given the current set of outgroups). The chosen outgroups in qpAdm needs to be differentially related to the sources such that a certain major ancestry is “anchored” in the test, which is rather heuristic. We used qpWave to test the resolution of a set of outgroups for distinguishing major ancestries among Eurasians, as well as the genetic cladility between populations given a set of outgroups. We used a set of eight outgroup populations in our study: Central African hunter-gatherers Mbuti.DG (n=5), indigenous Andamanese islanders Onge.DG (n=2), Taiwanese Aborigines Ami.DG (n=2), Native Americans Mixe.DG (n=3), early Holocene Levantine hunter-gatherers Natufian (n=6) (Lazaridis et al., 2016), early Neolithic Iranians Iran_N (n=8) (Lazaridis et al., 2016; Narasimhan et al., 2019), early Neolithic farmers from western Anatolia Anatolia_N (n=23) (Mathieson et al., 2015), and a Pleistocene European hunter-gatherer from northern Italy Villabruna (n=1) (Fu et al., 2016).

To evaluate potential sex bias (Fig. S6), we applied qpAdm to both the autosomes (default setting) and the X chromosome (adding “chrom:23” to the parameter file) for comparing the difference in the estimated ancestry porportions. For a certain ancestry, we calculated sex-bias Z score using the proportion difference between P_A_ and P_X_ divided by their standard errors (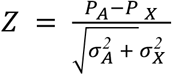, where *σ*_*A*_ and *σ*_*X*_ are the corresponding jackknife standard errors, as previously performed in (Mathieson et al., 2018). Therefore a positive Z score suggests autosomes harbor a certain ancestry more than X chromosomes do, indicating male-driven admixture. A negative Z score, in contrast, suggests female-driven admixture.

#### Phenotypic SNP analyses

We examined 49 SNPs in 17 genes (Table S8) known to be associated with phenotypic traits or with positive selection in Eurasia (Jeong et al., 2018). Given the low coverage of ancient DNA data, we focused on five of these genes and calculated the likelihood of allele frequency for SNPs in each ancient population based on the counts of reads covering on the SNP following a published strategy (Mathieson et al., 2015). In the allele frequency calculation, we classified all ancient individuals before Middle/Late Bronze Age into a single group, and kept three genetic groups during MLBA (Khövsgöl_LBA, Altai_MLBA, Ulaanzuukh), two genetic groups during Iron Age (Chandman_IA, SlabGrave), one group for Xiongnu, one group for Early Medieval and one group for Late Medieval. We calculated allele frequency at five loci (Table S8) that are associated with lactase persistence (*LCT*/*MCM6*), skin pigmentation (*OCA2, SLC24A5*), alcohol metabolism (*ADH1B*), and epithelial phenotypes including shovel-shaped incisor (*EDAR*) (Fig. 4).

## DATA AND CODE AVAILABILITY

All newly reported sequencing data are available from the European Nucleotide Archive, accession number PRJEB35748. 1240K genotype data are available on the Edmond Max Planck Data Repository under the link: https://edmond.mpdl.mpg.de/imeji/collection/2ZJSw35ZTTa18jEo.

## Supplementary Information

Supplementary Text

Materials and Methods

Figure S1. Geographic and ecological features in Mongolia

Figure S2. Selected DNA damage patterns in ancient individuals

Figure S3. Uniparental haplogroup assignments by group

Figure S4. PCA of present-day Eurasian populations used as background for Figs. 2 and S5

Figure S5. Genetic structure of Mongolia through time

Figure S6. Sex-bias Z scores by evaluating the differences of WSH-/Iranian-/Han-related ancestry on the autosomes and the X chromosome

Figure S7. Genetic ancestry changes in chronological order across all newly reported genetic groups

Figure S8. Genetic changes in the Eastern Steppe across time characterized by qpAdm with all individuals indicated

Figure S9. Outgroup *f3*-statistics for the pre-Bronze Age to Middle Bronze Age groups in the Eastern Steppe

Figure S10. Testing cladality of the four ANA populations using f4-statistics

Figure S11. Testing cladality of Afanasievo and Chemurchek using f4-statistics

Figure S12. Dating admixture in prehistoric individuals

Figure S13. Ancestry covariance in prehistoric individuals

Figure S14. Breakdown of the geographic-genetic correlation in Xiongnu

Figure S15. Comparing genetic homogeneity between ancient Mongol individuals and seven present-day Mongolic-speaking populations using qpWave

Table S1. Summary of archaeological sites and individuals by time period

Table S2. Overview of archaeological sites

Table S3. Overview of individuals and samples

Table S4. Overview and preservation assessment of DNA libraries and datasets

Table S5. Comparison of endogenous DNA % in paired samples and libraries

Table S6. Genetic sex and uniparental haplogroups

Table S7. Genetic relatives

Table S8. Phenotypic SNPs

Table S9. Identified population groups based on genetic clusters for all ancient individuals analysed in this study (n=214)

Table S10. Ancient populations used as ancestry proxies in the qpWave/qpAdm modeling

Table S11. List of 345 groups used for the population genetic analyses in this study

Table S12. Direct AMS radiocarbon dates for individuals in this study

Table S13. Genetic cladality and two-way admixture test in Ancient Northeast Asian (ANA) populations

Table S14. Genetic cladality and two-way admixture test for groups archaeologically affiliated with the Afanasievo culture

Table S15. 2-way and 3-way admixture modelling for the groups archaeologically affiliated with the Middle Bronze Age Chemurchek culture Dali_EBA, a published contemporaneous group from eastern Kazakhstan

Table S16. Genetic cladality test and 2-way admixture modelling for Late Bronze Age individuals

Table S17. 3-way admixture modelling for Early Iron Age groups

Table S18. Testing different Iranian proxies in TianShanSaka in 3-way admixture model

Table S19. qpAdm modelling results on Xiongnu

Table S20. Individual modelling results for individuals from the Xiongnu period

Table S21. Admixture modelling results for early Medieval individuals

Table S22. Admixture modeling results for late Medieval individuals

Table S23. Individual modelling results for individuals from the Mongol period

Table S24. Admixture modeling results from autosomes and X chromosome

References

## Notes

https://edmond.mpdl.mpg.de/imeji/collection/2ZJSw35ZTTa18jEo

