## Supplementary Materials for "A dynamic 6,000-year genetic history of Eurasia’s Eastern Steppe"

**This PDF file includes:**

**Supplementary Text**

6.6 Data filtering and compilation for population genetic analysis ……….…………....19
6.7 Analysis of population structure and relationships ………..……………………......19
6.8 Admixture modeling using qpAdm ……………...…………………….……………....20
6.9 Phenotypic SNP analysis ……………….………………………….……………………..20
7 Genetic clustering of ancient individuals into analysis groups …….………………….…..21
7.1 Pre-Bronze Age ……………………………………………..…………….…………21
7.2 Early Bronze Age ……………………………………….……………………….....22
7.3 Middle/Late Bronze Age ……………………………………………………………23
7.4 Early Iron Age …………………………………….…………………….…………...24
7.5 Xiongnu ………………………….…………………….……………….…………….....25
7.6 Early Medieval ………………………………………………………….…………...27
7.7 Late Medieval …………………………………….…………….……………...28

**Supplementary Figures**

Figure S1. Geographic and ecological features in Mongolia ………………………………………...29
Figure S2. Selected DNA damage patterns in ancient individuals …………………………………..30
Figure S3. Uniparental haplogroup assignments by group ……………………………………………..31
Figure S4. PCA of present-day Eurasian populations used as the background for Figs. 2 and S5…..32
Figure S5. Genetic structure of Mongolia through time ……………………………………………..33
Figure S6. Sex-bias Z scores by evaluating the differences of WSH-/Iranian-/Han-related ancestry on the autosomes and the X chromosome ………………………………………34
Figure S7. Genetic ancestry changes in chronological order across all newly reported genetic groups…………………………………………………………………………………..35
Figure S8. Genetic changes in the Eastern Steppe across time characterized by qpAdm with all individuals indicated ……………………………………………………………………..36
Figure S9. Outgroup *f3*-statistics for the pre-Bronze Age to Middle Bronze Age groups in the Eastern Steppe ……………………………………………………………………...37
Figure S10. Testing cladality of the four ANA populations using f4-statistics ……………………....38

**Supplementary Tables (provided as a separate .xlsx file)**

Table S1. Summary of archaeological sites and individuals by time period
Table S2. Overview of archaeological sites
Table S3. Overview of individuals and samples
Table S4. Overview and preservation assessment of DNA libraries and datasets
Table S5. Comparison of endogenous DNA % in paired samples and libraries
Table S6. Genetic sex and uniparental haplogroups
Table S7. Genetic relatives
Table S8. Phenotypic SNPs

Table S9. Identified population groups based on genetic clusters for all ancient individuals analysed in this study (n=214)
Table S10. Ancient populations used as ancestry proxies in the qpWave/qpAdm modeling
Table S11. List of 345 groups used for the population genetic analyses in this study
Table S12. Direct AMS radiocarbon dates for individuals in this study
Table S13. Genetic cladility and two-way admixture test in Ancient Northeast Asian (ANA) populations
Table S14. Genetic cladility and two-way admixture test for groups archaeologically affiliated with the Afanasievo culture
Table S15. 2-way and 3-way admixture modelling for the groups archaeologically affiliated with the Middle Bronze Age Chemurchek culture Dali_EBA, a published contemporaneous group from eastern Kazakhstan
Table S16. Genetic cladility test and 2-way admixture modelling for Late Bronze Age individuals
Table S17. 3-way admixture modelling for Early Iron Age groups
Table S18. Testing different Iranian proxies in TianShanSaka in 3-way admixture model
Table S19. qpAdm modelling results on Xiongnu
Table S20. Individual modelling results for individuals from the Xiongnu period
Table S21. Admixture modelling results for early Medieval individuals
Table S22. Admixture modeling results for late Medieval individuals
Table S23. Individual modelling results for individuals from the Mongol period
Table S24. Admixture modeling results from autosomes and X chromosome

**References for SI citations**………………..……….……………………….…..……..…………44

### Supplemental Text

### 1. Geography and ecology of Mongolia

Mongolia is located in Inner Asia between Russia and China, and it encompasses most of the Eurasian Eastern Steppe (Fig. 1). Mongolia has 21 aimags (provinces) and can be divided into ten geographic regions (Fig. S1a) with distinct ecological (Fig. S1b) and cultural features (Taylor et al., 2019). For example, far north Mongolia borders Siberia and includes both high mountain and mountain-taiga ecological zones, and it is the only aimag where reindeer pastoralism is practiced. North Mongolia is dominated by forest-steppe, but also contains mountain-taiga and steppe zones; cattle and yak pastoralism is particularly productive here, and Bulgan province is renowned for its horse pastoralism. The Altai region represents an extension of the Altai mountains from Russia into Mongolia and consists of a patchwork of environments including high mountains, valleys, and lakes, and ranging from forest steppe to desert steppe as the region stretches from north to south; pastoral economy in the Altai is mixed and differs by local environmental conditions. South Mongolia is dominated by the Gobi Desert, and it borders central and southeast Mongolia, which are largely characterized by desert-steppe; camel pastoralism is found throughout these regions. East Mongolia is a large expansive steppe zone that stretches to northeastern China. Today, mining is important in eastern Mongolia, as well as cattle, sheep, goat, and horse pastoralism.

### 2. Overview of Mongolian archaeology

Mongolian prehistory extends back more than 40,000 years, with documented sites ranging from the Upper Paleolithic to the present day. During nearly all of this time, lifeways in Mongolia have been nomadic, either supported by hunting, fishing and gathering or by pastoralism. The short-term and ephemeral nature of nomadic camp sites makes them difficult to identify on the landscape, and wind deflation has further reduced the visibility and preservation of many domestic sites. Only during the Bronze Age, with the sudden appearance of stone mounds and other burial features, do sites become more conspicuous and the archaeology better attested. As such, knowledge of Mongolian prehistory is strongly biased towards the past five millennia. The archaeology of Mongolia can be divided into 7 main periods: (1) pre-Bronze Age, prior to 3500 BCE; (2) Early Bronze Age, 3500-1900 BCE; (3) Middle/Late Bronze Age, 1900-900 BCE; (4) Early Iron Age, 900-300 BCE; (5) Xiongnu, 200 BCE to 100 CE; (6) Early Medieval, 100-850 CE, and (7) Late Medieval, 850-1650 CE. A brief summary of each period, as well as details for the sites included in this study, are provided below and in Data Tables S1, S2, and S6.

#### *2.1 Pre-Bronze Age (prior to 3500 BCE)*

The early archaeological record of Mongolia is poorly understood, particularly with respect to human remains and burials. While occasional finds provide direct evidence of *Homo sapiens* in Mongolia as far back as the Early Upper Paleolithic (Devièse et al., 2019), only a small handful of intentionally buried skeletons have been recovered prior to the end of the 4th millennium BCE. Although early and middle Holocene-era (10,000-3500 BCE) features and burials have been referred to as “Mesolithic,” “Neolithic,” or “Eneolithic” (Hanks, 2010), there is no direct evidence for domestic animals or a food-producing economy at any of these localities, although pottery was in wide use by the mid-Holocene (Janz et al.,

2017). Defining traits of most pre-Bronze Age burials in Mongolia include the absence of surficial construction features and the inclusion of artifacts made from animal remains such as deer and marmot (Eregzen, 2016). Two individuals in this study date to this pre-Bronze Age period in Mongolia. The first, dating to ca. 4600 BCE, is from Tamsagbulag (SOU) at the extreme eastern end of Mongolia and consists of a crouched pit burial, which may be associated with Chinese Neolithic cultures (https://edmond.mpdl.mpg.de/imeji/collection/2ZJSw35ZTTa18jEo). The second, dating to ca. 3700 BCE, is from Erdenemandal (ERM) in the Arkhangai province of north Mongolia (https://edmond.mpdl.mpg.de/imeji/collection/2ZJSw35ZTTa18jEo). Like the Tamsabulag burial, it lacked stone features and consisted of a simple crouched pit burial beneath a shallow earthen mound. The burial was recovered at a depth of 2.3 m and the grave mound appears to be part of a larger cemetery. In addition to these two pre-Bronze burials from Mongolia, we also analyzed four pre-Bronze individuals from the site of Fofonovo (FNO) near Lake Baikal in Buryatia, Russia (Lbova et al., 2008). All were buried in simple pit burials partially flexed (only their legs bent) as individuals, or sometimes several persons together. People at Fofonovo, like many others around Lake Baikal, were interred with many burial goods, including an array of bone and stone beads, neck pieces, chipped stone blades and points, bone harpoons, and pottery. Among these items were fragments or worked ornaments from wild boar, sable, and hawk.

Although pre-Bronze Age material from Mongolia is sparse, recent excavations in neighboring regions provide important context. In southern Russia, excavations by the Baikal Archaeological Project (BAP) at three sites (Lokomotiv, Shamanka II, and Ust'-Ida I) have enabled characterization of the Lake Baikal Neolithic Kitoi (5200-4200 BCE) and Isakovo (4000-3000 BCE) mortuary traditions, including genome sequencing of 14 of these hunter-gatherers (de Barros Damgaard et al., 2018) (Baikal_EN; also characterized in later studies as East Siberian Hunter Gatherers, ESHG) (Narasimhan et al., 2019). The genomes of six hunter-gatherers dating to ca. 5700 BCE are also available from the site of Devil's Gate (Sikora et al., 2019; Siska et al., 2017), a Neolithic cave site on the border between Russia and Korea. These individuals, separated by 2500 km, share a similar ancestry to each other and to modern Tungusic speakers in the lower Amur Basin, who we refer to as Ancient Northeast Asians (ANA).

#### *2.2 Early Bronze Age (ca. 3500-1900 BCE)*

Two major cultural phenomena associated with monumental mortuary architecture have been described in Mongolia during the Early Bronze Age (EBA): Afanasievo and Chemurchek. Both exhibit features linking them to ruminant pastoralism and to cultures further west.

**Afanasievo (3150-2750 BCE).** Beginning ca. 3150 BCE and persisting until ca. 2750 BCE, stone burials belonging to the Afanasievo culture type have been recovered from the Khangai Mountains in central Mongolia and the Altai Mountains of western Mongolia. These features contain the earliest direct evidence for domestic livestock (sheep/goat and cattle) in Mongolia, and are the oldest stone mounds recognized in Mongolia. They are defined by a circular, flat stone structure bounded by upright stones (https://edmond.mpdl.mpg.de/imeji/collection/2ZJSw35ZTTa18jEo). Internal burials are found with legs flexed inside an internal pit. In addition to domestic animal remains, these features sometimes contain metal artifacts (often reported as bronze, but not systematically distinguished from copper) and apparent deconstructed cart objects (Kovalev and Erdenebaatar, 2009). Similar features with a square burial morphology but otherwise identical construction, are sometimes labelled as Chemurchek (no remains from such features are included in this study). Recent analysis of proteins in human dental calculus from Afanasievo burials directly demonstrates the utilization of ruminant dairy products and the presence of domestic animals in the Afanasievo economy (Wilkin et al., 2019). We analyzed individuals from two Afanasievo sites in this study: Khuurai Gobi (KUR) and Shatar Chuluu (SHT).

**Chemurchek (2750-1900 BCE).** The Chemurchek culture (also called Hemtseg, Qiemu'erqieke, Shamirshak), spans the period between 2750 BCE-1900 BCE (Taylor et al., 2019). These features are found in western Mongolia and adjoining regions of bordering countries, including the Dzungar Basin in Xinjiang, eastern Kazakhstan, and parts of the Russian Altai (Honeychurch, 2017). The culture type is characterized by mounded stone burials with an internal stone cist burial chamber (https://edmond.mpdl.mpg.de/imeji/collection/2ZJSw35ZTTa18jEo). Similar to Afanasievo, burials found in Chemurchek tombs are in a legs-flexed body position. Adjacent to many Chemurchek burial features along the eastern side are anthropomorphic standing stones, sometimes depicted holding a shepherd's crook (https://edmond.mpdl.mpg.de/imeji/collection/2ZJSw35ZTTa18jEo). Inside the burials, artifacts such as stone bowls, bone tools, ceramics, and sometimes metal jewelry, have been recovered. Occasionally, non-funerary ritual structures containing animal remains are also attributed to this culture (Kovalev, 2014). Recent analysis of proteins in human dental calculus from these features confirmed the utilization of ruminant dairy products and the presence of domestic animals in the Chemurchek economy (Wilkin et al., 2019), although available radiocarbon chronology appears to preclude a meaningful exploitation of domestic horses (Taylor et al., 2019). We analyzed individuals from two Chemurchek sites in this study: Yagshiin Huduu (IAG/YAG) and Khundii Gobi (KUM).

Comparative genomic data are available for several contemporaneous sites in neighboring regions, including: (1) Botai, a horse hunter-herder site dating to ca. 3500 BCE in northern Kazakhstan (de Barros Damgaard et al., 2018); (2) multiple sites of Afanasievo ruminant pastoralists dating to ca. 3000-2500 BCE in the Kazakh and Russian Altai-Sayan region (Allentoft et al., 2015; Narasimhan et al., 2019); (3) Dali, a site in southeastern Kazakhstan whose lowest layers contain a woman dating to ca. 2650 BCE but lacking burial context (Narasimhan et al., 2019); (4) Gonur Tepe, a representative Bactria-Margiana Archaeological Complex (BMAC) site in Turkmenistan dating to 2300-1600 BCE (Narasimhan et al., 2019); and (5) three Lake Baikal sites, Ust'-Ida I, Shamanka II and Kurma Xi, associated with the Glazkovo mortuary tradition and dating to ca. 2200-1800 BCE (de Barros Damgaard et al., 2018; Damgaard et al., 2018).

#### *2.3 Middle/Late Bronze Age (ca. 1900-900 BCE)*

The Middle/Late Bronze Age (MLBA) in Mongolia is characterized by the sudden and widespread appearance of monumental mortuary architecture across Mongolia. Primarily taking the form of stone mounds, but also including stone stelae and other features, these Middle and Late Bronze Age structures remain among the most conspicuous features on the landscape even today. Middle and Late Bronze Age burial mound typology is complex and there is scholarly debate and disagreement on how to precisely define and delineate different mortuary types. In this study, we focused on four main burial forms: Mönkhkhairkhan burials, Baitag burials, Deer Stone-Khirigsuurs Complex (DSKC) burials, and Ulaanzuukh/Shape burials. We provide a general overview of these burial types, but acknowledge that not all scholars will agree with all details.

**Mönkhkhairkhan (1850-1350 BCE).** Dating to after the Chemurchek period, ca. 1850-1350 cal. BCE (Taylor et al., 2019), Mönkhkhairkhan burials are like their predecessors concentrated in northern and western areas of Mongolia, and characterized by a flexed leg burial position. Like Afanasievo, these features have a flat surface morphology and a central pit burial (https://edmond.mpdl.mpg.de/imeji/collection/2ZJSw35ZTTa18jEo), and they may be either circular or rectangular in overall shape – although unlike Afanasievo graves they are not constructed with a perimeter of upright slabs. Artifacts recovered from within these features include animal bones (wild, and perhaps domestic), bone jewelry, copper or bronze tools, and bone tools (Clark, 2015; Eregzen, 2016). Burials of this type represent the end of the flexed-leg burial tradition, which was replaced by prone (face-down) and supine traditions in the mid second millennium BCE. Because of the scarcity of these features, very little can be said about the economy of this culture or period, although the presence of wild animal fauna in at least one burial indicates a role for hunting, while connections with Chemurchek and Afanasievo would suggest the presence of domestic livestock. We analyzed individuals from three Mönkhkhairkhan sites in this study: Khukh Khoshuunii Boom (KHU), Shar Gobi-3 (SBG), and Ulaan Goviin Uzuur (UAA).

**Baitag (1050-900 BCE).** Found in a restricted region of western Mongolia, Baitag burials consist of non-mounded, small stone rings constructed from a single layer of small flat stone slabs (https://edmond.mpdl.mpg.de/imeji/collection/2ZJSw35ZTTa18jEo). A central burial pit oriented west-east contains a single individual oriented in a supine position with knees flexed. Unlike Deer Stone/Khirigsuur burials but similar to preceding Altai groups, such as the Mönkhkhairkhan and Chemurchek, the Baitag burials contain various small grave goods, including bronze and stone jewelry. These artifacts share similarities with those included in Karasuk culture graves from the Minusinsk Basin, as well as in burials Xinjiang and Gansu (Sibu culture) in northwestern China (Kovalev and Erdenebaatar, 2009). We analyzed individuals from one Baitag site in this study: Uliastai River Lower Terrace (ULI).

**Deer Stone-Khirigsuur Complex (DSKC) (1350-900 BCE).** This culture comprises three different monumental features - *khirigsuurs*, deer stones, and sagsai-style graves - and is tightly associated with the emergence of horsemanship in the Mongolian Steppe during the late second millennium BCE. In general, DSKC sites are concentrated in the western, northern, and central parts of Mongolia, with only a small number of sites further east (Honeychurch, 2015a). *Khirigsuurs* are large stone mounds, surrounded by an exterior fence that is either circular or rectangular in shape (https://edmond.mpdl.mpg.de/imeji/collection/2ZJSw35ZTTa18jEo). Although their exclusive function as burials is a subject of contention (Wright, 2012), *khirigsuurs* often contain a supine human body (Littleton et al., 2012) and do not typically yield other kinds of artifacts. Deer stones are anthropomorphic standing stones found either independently or co-occurring with *khirigsuurs*. Deriving their name from the common motif of stylized deer, carvings on these stelae also depict belts, weapons, and tools - and occasionally even a human face. Many of the weapons depicted on deer stones are of recognizably Karasuk style, bearing a strong resemblance to bronzes found in tombs in the Minusinsk Basin more than 500 km to the northwest (Honeychurch, 2015b), and the presence of deer stones in nearby Tuva further support the possibility of long-distance interaction between the Karasuk and the DSKC (Honeychurch, 2015b). At many Mongolian *khirigsuurs* and deer stones, smaller stone mounds containing the head, jaw, neck, and hooves of individual horses are found surrounding the eastern perimeter of the monument (https://edmond.mpdl.mpg.de/imeji/collection/2ZJSw35ZTTa18jEo). These horse mounds can range in number from a handful into the hundreds or thousands. Osteological study of DSKC horses reveal their use in transport and likely riding, as well as their sophisticated management as herd animals (Taylor et al., 2015, 2018). Another kind of satellite feature found at DSKC sites, open stone circles, often yield partial remains of sheep, goat, or cattle. Sagsai-style graves are often associated with the DSKC culture. These burials are also referred to as "slope burials" because of their common occurrence on the edge of hillslopes. Alternate names include *Munguntaiga* and even *khirigsuur.* Sagsai burials are similar to *khirigsuurs* in that they also have a supine burial position. They are sometimes (although not always) mounded, and can occasionally have a small external fence. Sagsai burials are either rectangular or circular; however, unlike *khirigsuurs*, these features have four upright corner posts and do not have external satellite mounds (https://edmond.mpdl.mpg.de/imeji/collection/2ZJSw35ZTTa18jEo). Similar to other DSKC features, they are concentrated in western and northern areas of Mongolia.

Radiocarbon modeling dates *khirigsuurs* to between ca. 1350-900 cal BCE, deer stones to ca. 1150-750 cal. BCE, and places the emergence of DSKC horse ritual at ca. 1200 cal BCE (Taylor et al., 2019, 2017). Sagsai-style graves fall within this range (1350-1050 BCE), further strengthening the claim for their affiliation to the DSKC culture sphere (Taylor et al., 2019). It should be noted, however, that these estimates could be influenced by dietary or taphonomic processes. In particular, later dates on deer stones may be influenced by radiocarbon contamination (Zazzo et al., 2019). Dairy proteins preserved in dental calculus demonstrate a pastoral, ruminant dairy-based economy at *khirigsuur* and sagsai sites (Jeong et al., 2018; Wilkin et al., 2019), and one sagsai site to date has also yielded evidence of horse milking (Wilkin et al., 2019). Perhaps buoyed by the innovation or adoption of mounted horseback riding and accompanying changes to the pastoral economy, deer stones and *khirigsuurs* proliferated over an extremely wide geographic range, reaching modern-day Tuva and southern Russia, Kazakhstan, Kyrgyzstan, and northwest China. We analyzed individuals from four DSKC sites in this study: Arbulag Soum (ARS), Berkh Mountain (BER), Uliastai River Lower Terrace (ULI), and Uushigiin Uver (UUS).

**Ulaanzuukh/Shape Burial (1450-1150 BCE).** Beginning in the mid-second millennium BCE, a number of different burial traditions emerged in the southern and eastern regions of Mongolia. United by a common prone or face-down burial position, these groups are sometimes considered a single cultural unit, and other times classified separately as discrete burial types (Honeychurch, 2015b). Ulaanzuukh features (named after the type site in southeastern Mongolia), are non-mounded square or rectangular features with a wall of upright slabs or layered stone and a central pit (https://edmond.mpdl.mpg.de/imeji/collection/2ZJSw35ZTTa18jEo). Shape burials, also called *Tevsh*, ant-shaped, hourglass-shaped, and other names, are similar, but with a waisted hourglass-style edge construction (https://edmond.mpdl.mpg.de/imeji/collection/2ZJSw35ZTTa18jEo). These features are typically made of layered stone, and sometimes with a single edge ringed with upright slabs. Other variations often included within this culture group include D-shaped or stirrup shaped graves with a prone body position. Radiocarbon modeling suggests that Ulaanzuukh features date to ca. 1450-1150 BCE, while shape burials could both predate and postdate this mark – although very few have been reliably dated (Taylor et al., 2019). Burials of this culture often contain apparently domestic livestock remains, including sheep, goat, horse, and cattle (Nelson et al., 2009), although the earliest horses from these features date to only ca. 1250 BCE (Taylor et al., 2017). Recent analysis of proteins in human dental calculus has confirmed the utilization of ruminant dairy products and the presence of domestic animals in the Ulaanzuukh economy (Wilkin et al., 2019). A few bronze knives of Karasuk origin have been found in Ulaanzuukh-Tevsh graves, indicating possible long-distance connections to the Minusinsk basin (Honeychurch, 2015b). We analyzed individuals from two Ulaanzuukh sites in this study: Bulgiin Ekh (BUL) and Ulaanzuukh (ULN). In addition, we also analyzed individuals from four additional Late Bronze Age sites with uncertain or unclassified cultural affiliations: Biluutiin Am (BIL), Khoit Tsenkher (KHI), Uliastai Zastav II (ULZ), and Tsaidam Bag (TSB/TSI).

Comparative genomic data are available for several contemporaneous archaeological sites in neighboring regions, including: (1) four Okunevo sites (Verkhni Askiz, Okunev Ulus, Uybat, Syda 5), dating to 2200-2600 BCE (Allentoft et al., 2015; de Barros Damgaard et al., 2018); (2) five Sintashta sites (Bulanovo, Tanabergen II, Stepnoe VII, and Bol'shekaraganskii, Kamennyi Ambar 5 cemetery), dating to ca. 2200-1800 BCE (Allentoft et al., 2015; Narasimhan et al., 2019); (3) four Central Steppe sites near Krasnoyarsk in western Siberia (Krasnoyarsk Krai, Potroshilovo II, Ust-Bir IV, Chumyash-Perekat-1) dating to 1400-1700 BCE (Narasimhan et al., 2019); (4) three Karasuk sites (Arban I, Sabinka II, and Bystrovka), dating to ca. 1400-1300 BCE (Allentoft et al., 2015).

#### *2.4 Early Iron Age (ca. 900-300 BCE)*

The Early Iron Age cultures of Inner Asia arose during a time of new technological advancements, including the development of composite bows and the beginnings of iron metallurgy used for items like arrows and horse-riding equipment (Honeychurch, 2015a). These cultures include (1) the widespread Slab Grave culture, prevalent in eastern, southeastern, and central Mongolia as well as East Baikal and parts of northern China, and (2) the Uyuk and Pazyryk cultures in the Sayan-Altai and portions of northwestern Mongolia. These latter cultures were part of a broader "Scythian" cultural phenomenon that spread into eastern Kazakhstan and across the Eurasian steppes, and which was related to Saka groups of northern Iran and the Tian Shan mountains. The Saka were an Iranian group broadly associated with the Scythians. Their later (after 200 BCE) military activities in Sogdia, Bactria, and the Tian Shan were recorded by Persian, Greek, and Chinese sources (Beckwith, 2009). Alongside the technological advancements of the Early Iron Age came increased long-distance interactions and the adoption of grain subsistence by cultures such as the Uyuk in areas outside of the central Mongolian Steppe, but not yet by groups like the Slab Grave culture within Mongolia (Ventresca Miller and Makarewicz, 2019).

**Slab Grave (1000-300 BCE).** Beginning around 1000 BCE, a new and burial style known as Slab Grave began appearing in eastern Mongolia. Slab graves are so called because of the large stone slabs used to mark the surface of the burial and to contain the rectangular burial space (hence in Mongolian they are called "square burials") wherein single individuals are interred (Tsybiktarov, 1998). Although occasionally found singly, Slab Grave burials are more typically grouped into small cemeteries (Honeychurch, 2015a). Stone slabs are set upright in the ground, and are thus prominent grave markers (https://edmond.mpdl.mpg.de/imeji/collection/2ZJSw35ZTTa18jEo). The burial pits are quite shallow, and human remains are rarely found complete or in good preservation. Over time, the Slab Grave culture expands northwards into eastern Baikal and westwards into central Mongolia, where it intrudes into former DSKC territory. Some slab graves tear apart the stone structures of *khirigsuurs* to construct the graves, and some even reuse deer stones for standing corner stones or laid-down slabs within the burial pit (Honeychurch, 2015a). Unlike earlier Bronze Age burials, grave goods become more common in Slab Grave burials, consisting primarily of bronze beads, buttons, and small ornaments, as well as horse gear, arrowheads, axes, and knives. Stone, ceramic, and bone artifacts are also found in slab graves, and a few burials contained tripod-shaped pottery similar to those from Inner Mongolia and Manchuria or other non-local grave goods such as turquoise and carnelian beads from Central or South Asia (Honeychurch, 2015a).

Portions of livestock are often set at the edge or just outside of the rectangular burial space. In addition to faunal remains demonstrating the presence of domestic animals in the Slab Grave economy, recent analysis of proteins in human dental calculus has confirmed the utilization of ruminant and horse dairy products (Wilkin et al., 2019). Although the Slab Grave phenomenon emerges out of the former territory of the Ulaanzuuk culture, archaeological evidence for the relationship between these two groups has been ambiguous. Nevertheless, the similarity of bronze artifacts, especially relating to horse gear and weaponry, found at Slab Grave sites to similar artifacts found in the Altai, Tuva, and Minusinsk regions may indicate a continuation of previously established long-distance relationships between these regions (Honeychurch, 2015b). We analyzed individuals from five Slab Grave sites in this study: Bor Bulag (BOR), Morin Tolgoi (MIT), Darsagt (DAR), Shunkhlai Mountain (SHU), and Pesterevo 82 (PTO).

**Uyuk (700-200 BCE).** This Early Iron Age culture centered in the Upper Yenisei River area, in modern-day Tuva, with some extensions into northwestern Mongolia (Murphy, 2003; Savinov, 2002). This culture is also referred to as the Aldy-Bel or Sagly-Bazhy culture, and is known best in Mongolia by the thoroughly excavated site of Chandman Mountain included in this study (Tseveendorj, 1980) (https://edmond.mpdl.mpg.de/imeji/collection/2ZJSw35ZTTa18jEo). Graves were marked by a round pile of stones and are often found in cemeteries of one to two dozen graves. Beneath the stone mounds are large log chambers containing several individuals (often assumed to be kin as they include men, women and children) all laid in partially flexed positions on their sides. Portions of sheep are also often placed in the graves. The Uyuk log chambers resemble similar log architecture constructed by the contemporaneous Pazyryk culture in the Russian Altai and surrounding areas, and both the Uyuk and Pazryk have been associated with the broader Saka culture (Parzinger, 2006). Similar to Slab Graves, recent analysis of proteins in human dental calculus has confirmed the utilization of ruminant and horse milk among those at Chandman Mountain (Wilkin et al., 2019). Isotopic studies have also shown that some Uyuk communities, including at Chandman Mountain, had a significant amount of millet in their diet (Murphy et al., 2013; Ventresca Miller and Makarewicz, 2019; Wilkin et al.). This links them to agropastoralist cultures of the southern steppe and Central Asia, where millet cultivation was widely adopted during the westward spread of the cereal from China to the Caucasus during the second millennium BCE (Ventresca Miller and Makarewicz, 2019). We analyzed individuals from one Uyuk site in this study: Chandman Mountain (CHN).

**Pazyryk (500-200 BCE).** This culture is known mainly for its type site of Pazyryk, whose large tombs contain numerous exotic imports, including silks from China and textiles from Achaemenid Persia (Rudenko, 1970). Pazryk burials are found mostly within the northern Altai areas of Russia, far eastern Kazakhstan (Samashev, 2011) and northwestern Mongolia (Törbat et al., 2009). Similar to Uyuk and other 'Saka' style graves, Pazyryk burials are marked by round piles of stones. Beneath these stone piles, however, most Pazyryk graves have smaller wooden chambers with only one or two persons; their size and burial goods vary greatly, though many of them are accompanied by whole horses laid beside the burial chamber (Kubarev and Shul'ga, 2007) (https://edmond.mpdl.mpg.de/imeji/collection/2ZJSw35ZTTa18jEo). No new Pazyryk individuals were included in this study; however, they are important to consider because the northern Altai practice of whole horse burials later appears in scattered central Mongolia cemeteries of the subsequent Xiongnu period. Genome-wide data from Pazyryk individuals have been previously reported from site of Berel in Altai region of Kazakhstan (Unterländer et al., 2017).

Comparative genomic data are available for several contemporaneous sites in neighboring regions, including: (1) the early Sarmatian site Pokrovka in southwestern Russia, dating to ca. 500-100 BCE (Unterländer et al., 2017), a Scythian individual from the Samara region dating to ca. 300 BCE (Mathieson et al., 2015), and nine Sarmatian sites in southwestern Russia (Chebotarev V, Kamyshevahsky X, Nesvetay II, Nesvetay IV and Tengyz), northern Kazakhstan (Bestamak and Naurzum Necropolis), and the southern Ural region (Cherniy Yar and Temyaysovo) (Damgaard et al., 2018; Krzewińska et al., 2018); (2) the Pazyryk site of Berel in the Altai, dating to ca. 400-200 BCE (Unterländer et al., 2017); and (3) the Saka sites of Borli, Karasjok-1, Karasjok-6, Nazar-2, Sjartas (Zjartas), and Taldy-2 in Kazakhstan (Damgaard et al., 2018), and the sites of Basquiat I, Keden, and Ornek in the Tian Shan (Damgaard et al., 2018); and (4) the Tagar site of Grishkin Log 1 in the Minusinsk Basin (Damgaard et al., 2018). Data from three other potentially relevant sites (the Aldy-Bel site Arzhan 2 in Tuva, dating to ca. 700-500 BCE, and the Zevakino-Chilikta sites Ismailovo and Zevakino in eastern Kazakhstan, dating to ca. 900-600 BCE; (Unterländer et al., 2017) were excluded from analysis due to insufficient genetic coverage for comparison.

#### *2.5 Xiongnu (ca. 200 BCE to 100 CE)*

During the late first millennium BCE, a radically new multi-regional political entity formed in Mongolia, known as the Xiongnu empire. The Xiongnu empire is attested not only by historical records but also by ample archaeological remains throughout Inner Asia (Brosseder and Miller, 2011; Honeychurch, 2015a). For roughly three centuries the Xiongnu ruled from their core realms in central and eastern Mongolia, expanding into western Mongolia, northern China and eastern Baikal, as well as making inroads into more distant regions in Central Asia. Most graves of the Xiongnu period were shaft pits set beneath thick rings of stones on the surface. These burials represent the vast network of regional and local elites and not the "commoner" people of Xiongnu society, whose burials are far less conspicuous, lying under small piles of stones or in unmarked pits. The graves of the uppermost ruling elites of the empire, on the other hand, were constructed on a far grander scale than that of ring graves.

While ring grave structures are found throughout the entire Xiongnu era, prestige accoutrements (and to some degree burial rituals) changed during the course of the empire. According to these changes, we can discern a general division between Early (200-50 BCE) and Late (50 BCE - 100 CE) Xiongnu periods (Miller, 2014). Overall, Xiongnu graves are marked by a dramatic increase in grave goods and furnishings as compared to previous time periods and cultures in Mongolia. As the Xiongnu expanded their empire, they conquered numerous neighboring groups to their east and west as well as subduing their Han Chinese neighbors to the south (Di Cosmo, 2002). They continually traded and warred with Han China, defying the Great Wall boundaries, and held significant sway over the Silk Road kingdoms of Central Asia (Hulsewé, 1979). The findings of exotic items from China, Persia and the Mediterranean attest to these far-flung interactions, with Egyptian-style faience beads in graves of local elites and Roman glass bowls in the tombs of the rulers (Miller and Brosseder, 2017; 2011). The end of the Xiongnu period ca. 100 CE is marked by the widespread decline of Xiongnu power and influence following defeats by the Xianbei in northeastern China and the Han Dynasty of China, although isolated groups from the Xiongnu empire continued to exist in northern China until the 5th century CE.

**Early Xiongnu (200-50 BCE).** Prestige items during the Early Xiongnu period are dominated by large bronze belt pieces; however, burial customs within graves of the Early period varied to a great degree between regions. One example of this occurs at Salkhityn Am cemetery, where rituals of ring graves show a high degree of variation, even including offerings of whole horses that are more typical of Altai elites such as those in Pazyryk graves (Ölziibayar et al., 2019) (https://edmond.mpdl.mpg.de/imeji/collection/2ZJSw35ZTTa18jEo). We analyzed individuals from three Early Xiongnu sites in this study: Astyn Gol (AST), Buural Uul (BAU/BRL/BUU), and Salkhityn Am (SKT).

**Late Xiongnu (50 BCE - 100 CE).** Prestige items in the Late Xiongnu period shift to more iron items, often covered with gold foil or even inlaid with precious stones, and increasingly focused on long-distance exotic materials. At the same time, burial customs in ring graves throughout the empire become more regularized. Most elites were buried in wooden coffins in shaft pits with livestock portions and ceramic vessels set beside the coffin (https://edmond.mpdl.mpg.de/imeji/collection/2ZJSw35ZTTa18jEo). During the Late period, the high ruling Xiongnu elites adopted a radically new form of burial structure. These square tombs were marked on the surface by rectangular stone structures with trapezoidal 'ramp' entryways, their burial pits were extremely deep, and wooden coffins were decorated and nested within larger wooden chambers (https://edmond.mpdl.mpg.de/imeji/collection/2ZJSw35ZTTa18jEo). We analyzed individuals from 26 Late Xiongnu sites in this study: Atsyn Am (ATS), Baruun Mukhdagiin Am (BAM), Baruun Khovdiin Am (BRU), Burkhan Tolgoi (BTO), Chandman Mountain (CHN), Delgerkhaan Uul (DEL), Khanan Uul (DOL), Duulga Uul (DUU); Emeel Tolgoi (EME), Khudgiin Am (HUD), Ikh Tokhoirol (IKT), Il'movaya Pad (IMA), Jargalantyn Am (also called Jargalantyn Khondii; JAA/JAG), Tarvagatain Am (also called Khoit Tsenkher; KHO), Naimaa Tolgoi (NAI), Sant Uul (SAN), Solbi Uul (SOL), Songino Khairkhan (SON), Takhityn Khotgor (TAK), Tavan Tolgoi (TAV), Tevsh Mountain (TEV), Ulaanzuukh (ULN), Ovgont (UVG), Yuroo II (YUR), Tamiryn Ulaan Khoshuu (also called Burkhan Tolgoi; BUR/TMI/TUH/TUK), and Uguumur Uul (UGU).

Comparative genomic data are available for a few contemporaneous sites in neighboring regions, including: (1) two early Xiongnu individuals from Khövsgöl (Hovsgol) province dating to 50-350 BCE (Damgaard et al., 2018); (2) a late Xiongnu royal tomb (DA39.SG) in Arkhangai dating to 80-160 CE (Damgaard et al., 2018).

#### *2.6 Early Medieval (ca. 100-850 CE)*

After the fall of the Xiongnu, Xianbei groups from northeast China pushed into Mongolia, although historical and archaeological evidence for the establishment of large and long-lasting Xianbei polities appears only in northern China, not in Mongolia (Miller, 2016). One individual in this study (TUK001) at the site of Tamiryn Ulaan Khoshuu (Burkhan Tolgoi) dates to the era of Xianbei power in Inner Asia; however, there is no cultural context that could affirm affiliation with the Xianbei or other groups of northeastern China. Instead, recent excavations at this site have yielded artifacts, such as pottery from the Kwarezm oasis cultures near the Aral Sea and coins of the Sassanian Persian empire, that indicate significant interactions with areas in Central Asia and much farther west. In the mid-fourth century, a large polity known as the Rouran purportedly took over all of Mongolia; however, there is little recorded history about the Rouran (Kradin, 2005), and only one grave found so far can be dated to the Rouran era (Li et al., 2018; Nei Menggu zizhiqu wenwu kaogu yanjiusuo and International Institute for the Study of Nomadic Civilizations, 2015). The archaeology of the second to sixth centuries in Mongolia, i.e., the Xianbei and Rouran eras, constitute an extremely new field of research (Odbaatar and Egiimaa, 2018).

The most prominent political entities in the Early Medieval era are the Türk and Uyghur empires, the latter being an immediate dynastic takeover from the former. Numerous burials of the Türk era have been unearthed in Mongolia. By contrast, far fewer Uyghur burials have been identified and excavated to date.

**Türk (550-750 CE).** Göktürkic tribes of the Altai Mountains established a political structure across Eurasia beginning in 552 CE, with an empire that ruled over Mongolia from 581-742 CE (Golden, 1992). A brief period of disunion occurred between 659-682 CE, during which the Chinese Tang dynasty laid claim over Mongolia. One individual from this study (TUM001) was a sacrificial person within the ramp of a Chinese-style tomb in central Mongolia dating (via tomb inscription) to this exact time period. The other Türkic era individuals in this study were excavated from conventional Türkic style graves. Features of the Türk period include numerous stone statues and stone offering boxes across the steppe landscape, while burials are often arranged as small groups of graves or single graves inserted into burial grounds of earlier Bronze to Iron ages. Most elites were interred within wooden coffins as single individuals buried beneath a stone mound, and many were buried with whole horses equipped with riding gear (https://edmond.mpdl.mpg.de/imeji/collection/2ZJSw35ZTTa18jEo). Other burials were in small wooden coffins without whole horses beside them. We analyzed individuals from 5 Türk sites in this study: Nomgonii Khundii (NOM), Shoroon Bumbagar (Türkic mausoleum; TUM), Zaan-Khoshuu (ZAA), Uliastai River Lower Terrace (ULI), and Umuumur uul (UGU).

**Uyghur (750-850 CE)**. In the mid-eighth century, Uyghur tribes from the Upper Yenesei region overthrew the Türk rulers and immediately established a Mongolia-based empire, taking over the Orkhon valley as their capital and establishing a dynasty from 744-840 CE (Mackerras 1972). Most Uyghur period burials excavated to date, including those from the Olon Dov burial ground (OLN) included in this study, lie in the vicinity of the Kharbalgas capital in the Orkhon Valley. Most of the burials excavated were discovered beneath large earthen enclosures that contained ritual structures for venerating the uppermost elites. These conspicuous ritual enclosures occur as single monuments or in small groups, and they are found in several locations throughout the foothills of the nearby the Uyghur capital. These monumental tombs with ramp entries and vaulted brick chambers were likely reserved for the ruling nobility of the Uyghur empire (Odbaatar 2016) (https://edmond.mpdl.mpg.de/imeji/collection/2ZJSw35ZTTa18jEo). One individual in this study (OLN006) was found in a monumental tomb. A second, more modest category of Uyghur burials consists of stone structures placed on the surface, either square or round in shape, that contain multiple individuals (Erdenebat 2016) (https://edmond.mpdl.mpg.de/imeji/collection/2ZJSw35ZTTa18jEo). Dozens of these burials have been documented at Olondov (Erdenebat et al., 2012), and most of the Uyghur individuals in this study are from such graves. One such grave at Olon Dov, grave 19, contained the remains of multiple individuals, six of whom are included in this study. Other scattered examples of single Uyghur graves have been found in Mongolia, and we analyzed one of these (ZAA001) from the site of Zaan-Khoshuu. Although a few large 'royal' complexes have been found elsewhere in central Mongolia, no significant cemeteries outside the capital region have yet been found. We analyzed individuals from two Uyghur sites in this study: Olondov (OLN) and Zaan-Khoshuu (ZAA).

Comparative genomic data are available for contemporaneous sites in neighboring regions, including: (1) Alan sites in North Ossetia-Alania and Alan 51 from the Caucasus (Damgaard et al., 2018); (2) the Rouran site of Khermen Tal site from Arkhanggai, Mongolia (Li et al., 2018).

#### *2.7 Late Medieval (ca. 850-1650 CE)*

This period in Mongolia is dominated mostly by the power struggles of two empires established by the Khitans (907-1125 CE) and the Mongols (1206-1368 CE). Burials from the Khitan era are virtually unknown in Mongolia, whereas numerous graves from the Mongol era have been documented and unearthed. So-called cave burials are known from both periods (Bemmann and Nomguunsüren, 2012), but their human remains were not included in this study.

**Khitan (ca. 900-1100 CE).** After the collapse of the Uyghur empire in Mongolia in 840 CE, the Khitans of northeast China established the powerful Liao Dynasty in 916 CE. Although based in Manchuria, the Khitans conquered and controlled the steppe of present-day Mongolia through a system of garrisons and long walls, deporting people from other conquered regions, such as northern Korea, to Mongolia (Kradin and Ivliev, 2008). The dissolution of the Khitan empire in 1125 CE lead to a power vacuum in Mongolia until the rise of Chinggis Khan in the early 13th century CE. To date, very few Khitan era graves have been found in Mongolia. The site of Ulaan Kherem II (ULA) has yielded one Khitan-era grave (ULA001), and two Khitan-era unmarked graves of a man and woman were also discovered during the excavation of a Xiongnu settlement at Zaan Khoshuu (ZAA) beneath an older collapsed building (Nei Menggu zizhiqu wenwu kaogu yanjiusuo and International Institute for the Study of Nomadic Civilizations, 2015; Ochir et al., 2016). The man, found in a pit within the pit-house, was buried in a simple pit with a quiver and arrows. The woman, found nearby a pit-house, was buried in full dress and placed in a supine position with her head to the northwest inside a wooden coffin, along with pottery of the Khitan era (https://edmond.mpdl.mpg.de/imeji/collection/2ZJSw35ZTTa18jEo). These burials are significantly different in form and structure from other Khitan burials in northern China, where the core of the empire was located. At present, no monumental tombs of high Khitan elites have been found in Mongolia.

**Mongol (ca. 1200-1400 CE).** The home base of the Mongol tribe was in the forest-steppe zone at the Onon and Kerülen (Kherlen) rivers in northeastern Mongolia. From this core region they successfully conquered the Eurasian steppes and most of their sedentary neighbors in the adjacent regions. Historical records indicate that they transferred a large number of defeated people, war captives and slaves all over their growing empire; they also fostered trade, the exchange of knowledge, techniques, and technicians (Allsen, 2015). Mongol burials are typically situated in small groups on flat southern slopes or placed within Bronze and Iron Age cemeteries. They are marked above ground with stones in an irregular, flat, oval or rounded, one-layered setting (https://edmond.mpdl.mpg.de/imeji/collection/2ZJSw35ZTTa18jEo). The pit is normally between 50-150 cm deep, rarely deeper, and very seldom constructed as a niche. Typical Mongol burials contain one person placed in supine position and sometimes in a wooden coffin, with the head to the north. A very characteristic feature of Mongol burials is the inclusion of a tibia from small livestock, mostly sheep, placed near the head and sometimes in a vessel. There are two ideal burial types concerning grave goods: one equipped with bow, arrow, quiver, horse equipment, and belt with attachments, and a second with scissors, a comb, a mirror, beads and a *bogtag* – a long hat made out of birch bark, covered with silk and decorated with golden ornament. Graves of these standard types are spread all over Mongolia, and at present no regional differences have been reported and no monumental burials are known (Erdenebat, 2009; Lkhagvasüren, 2007). The Mongol burials included in this study are of these types, which consist of the burials of local steppe warriors and elites of the Mongol empire. Individuals from the cosmopolitan capital of Karakorum were not sampled in this study.

Historical records mention a large amount of foreign people who migrated, whether for opportunity or by force, into the core steppe regions of the Mongol empire (Allsen, 2015). Given the intriguing results of extreme genetic diversity among local elite constituents for the Xiongnu era, one might expect a similar or even greater diversity during the Mongol era. However, within the core steppe realms, lower local levels of the Mongol empire appear not to have been as open. The supposed mass of incoming foreigners must be sought in other burial contexts, not those of Mongol tradition.

Because no comparative genomic data are available for contemporaneous sites, we compared our Late Medieval data to modern Mongolic speaking populations (Buryat, Khamnegan, Kalmyk, Mongol, Daur, Tu, Mongola) (Jeong et al., 2019; Lazaridis et al., 2014; Patterson et al., 2012).

### 3. Study design and sample selection

Here we present new genome-wide data for 213 ancient individuals from Mongolia and 13 individuals from Buryatia, Russia, which we analyze together with 21 previously published ancient Mongolian individuals (Jeong et al., 2018), for a total of 247 individuals. All new Mongolian individuals, except ERM, were sampled from the physical anthropology collections at the National University of Mongolia and the Institute for Archaeology and Ethnology in Ulaanbaatar, Mongolia. ERM001/002/003 was provided by Jan Bemmann. Russian samples were collected from the Institute for Mongolian, Buddhist, and Tibetan Research as well as the Buryat Scientific Center, Russian Academy of Sciences (RAS).

Together, this ancient Eastern Steppe dataset of 247 individuals originates from 89 archaeological sites (Fig. 1; Table S1) and spans approximately 6,000 years of time (Data Tables S1, S2, S6). High quality genetic data was successfully generated for 214 individuals and was used for population genetic analysis (Table S4). Subsistence information inferred from proteomic analysis of dental calculus has been recently published for a subset of these individuals (n=32; (Wilkin et al., 2019)), and stable isotope analysis of bone collagen and enamel (n=137) is also in progress (Wilkin et al.); together, these data allow direct comparison between the biological ancestry of specific archaeological cultures and their diets, particularly with respect to their dairy and millet consumption.

### 4. Radiocarbon dating of sample materials

A total of 24 new radiocarbon dates were obtained by accelerator mass spectrometry (AMS) of bone and tooth material at the Curt-Engelhorn-Zentrum Archäometrie (CEZA) in Mannheim, Germany (n=22) and the University of Cologne Centre for Accelerator Mass Spectrometry (CologneAMS) (n=2). Selection for radiocarbon dating was made for all burials with ambiguous or unusual burial context and for all individuals appearing as genetic outliers for their assigned period. Uncalibrated direct carbon dates were successfully obtained for all bone and tooth samples (Table S12). An additional 81 previously published radiocarbon dates for individuals in this study were also compiled and analyzed, making the total number of directly dated individuals in this study 105. Dates were calibrated using OxCal v.4.3.2 (Ramsey, 2017) with the r:5 IntCal13 atmospheric curve (Reimer et al., 2013).

Of the 105 total radiocarbon dates analyzed in this study, 27 conflicted with archaeological period designations reported in excavation field notes or previous publications (Table S12). Four burials of uncertain cultural context were successfully assigned to the Middle/Late Bronze Age (BIL001, MIT001) and Late Medieval periods (UUS002, ZAA003). One burial originally assigned to the Late Medieval period was reassigned to the pre-Bronze Age following radiocarbon dating (ERM001), and three burials originally assigned to the Middle/Late Bronze Age were similarly reassigned to the Early Bronze Age (KUM001, KUR001, IAG001). This suggests that early burials may be underreported in the literature because they are mistaken for later graves. Likewise three burials originally classified as Late Medieval were found to be hundreds or thousands of years older, dating to the Early Medieval (TSB001) and Middle/Late Bronze Age (ULZ001, TSI001) periods. Although some highly differentiated burial forms can be characteristic of specific locations and time periods, simple burial mounds also exist for all periods and - lacking distinctive features - they can be difficult if not impossible to date without radiometric assistance.

In addition to early burials being mistaken for later ones, late burials were also misassigned to earlier periods. For example, three burials originally assigned to the Middle/Late Bronze Age were determined to date to the Early Iron Age (DAR001), Xiongnu (ULN004), and Late Medieval (SHU001) periods, and two Early Iron Age (CHN010, CHN014), six Xiongnu (TUK001, UGU001, DUU002, BRL001, BAU001, DEE001), and two Early Medieval (ULA001, ZAA005) graves were likewise reassigned to later periods following radiocarbon dating. Part of the difficulty in correctly assigning archaeological period to later burials relates to the frequent reuse of earlier graves and cemeteries by populations from later periods. The site reports of several Xiongnu excavations noted burial intrusions, displaced burials, and other indications of burial disturbance and reuse. However, evidence of burial reuse may also be subtle and easily overlooked. As such, we recommend great care in making cultural or temporal assignments at multi-period cemeteries or for any burials showing evidence of disturbance.

### 5. Procedures for ancient DNA recovery and sequencing

#### *5.1 Sampling*

Sampling was performed on a total of 169 teeth and 75 petrosal bones from fragmented crania originating from 225 individuals (Table S3). For 14 individuals, both a tooth and a petrosal bone were sampled (Table S5). For three individuals, two teeth were sampled, and for one individual, two teeth and one petrosal bone were sampled (Table S5). For Mongolian material, whole teeth and petrosal bone (except ERM) were collected at the physical anthropology collections of the National University of Mongolia and the Institute for Archaeology and Ethnology under the guidance and supervision of Erdene Myagmar and S. Ulziibayar. Petrosal and tooth material from ERM were provided by Jan Bemmann. For Russian material, whole teeth alongside petrosal bone or bone were collected from the Institute for Mongolian, Buddhist, and Tibetan Research as well as the Buryat Scientific Center, Russian Academy of Sciences (RAS). After collection, the selected human skeletal material was transferred to the Max Planck Institute for the Science of Human History (MPI-SHH) for genetic analysis.

#### *5.2 Laboratory procedures for genetic data generation*

Genomic DNA extraction and Illumina double-stranded DNA (dsDNA) sequencing library preparation were performed for all samples in a dedicated ancient DNA clean room facility at the MPI-SHH, following published protocols (Dabney et al., 2013) with slight modifications (Mann et al., 2018). We applied a partial treatment of the Uracil-DNA-glycosylase (UDG) enzyme to confine DNA damage to the ends of ancient DNA molecules (Rohland et al., 2015). Such "UDG-half" libraries allow us to minimize errors in the aligned genetic sequence data while also maintaining terminal DNA misincorporation patterns needed for DNA damage-based authentication. Library preparation included double indexing by adding unique 8-mer index sequences at both P5 and P7 Illumina adapters. After shallow shotgun sequencing for screening, we enriched libraries of 195 individuals with ≥ 0.1% reads mapped on the human reference genome (hs37d5; GRCh37 with decoy sequences) for approximately 1.24 million informative nuclear SNPs ("1240K") by performing an in-solution capture using oligonucleotide probes matching for the target sites (Mathieson et al., 2015). In addition, eight samples (see Table S5; Table S4) were also selected and built into single-stranded DNA (ssDNA) sequencing libraries for comparison. Single-end 75 base pair (bp) or paired-end 50 bp sequences were generated for all shotgun and captured libraries on the Illumina HiSeq 4000 platform following manufacturer protocols. Output reads were demultiplexed by allowing one mismatch in each of the two 8-mer indices.

### 6. Genetic data analysis

#### *6.1 Sequence data processing*

Short read sequencing data were processed by an automated workflow using the EAGER v1.92.55 program (Peltzer et al., 2016). Specifically, in EAGER, Illumina adapter sequences were trimmed from sequencing data and overlapping sequence pairs were merged using AdapterRemoval v2.2.0 (Schubert et al., 2016). Adapter-trimmed and merged reads with 30 or more bases were then aligned to the human reference genome with decoy sequences (hs37d5) using BWA aln/samse v0.7.12 (Li and Durbin, 2009). A non-default parameter "-n 0.01" was applied. PCR duplicates were removed using dedup v0.12.2 (Peltzer et al., 2016). Based on the patterns of DNA misincorporation, we masked the first and last two bases of each read for UDG-half libraries and 10 bases for non-UDG single-stranded libraries, using the trimbam function in bamUtils v1.0.13 (Jun et al., 2015), to remove deamination-based 5' C>T and 3' G>A misincorporations. Then, we generated pileup data using samtools mpileup module (Li and Durbin, 2009), using bases with Phred-scale quality score ≥ 30 ("-Q30") on reads with Phred-scale mapping quality score ≥ 30 ("-q30") from the original and the end-masked BAM files. Finally, we randomly chose one base from pileup for SNPs in the 1240K capture panel for downstream population genetic analysis using the pileupCaller program v1.2.2 (https://github.com/stschiff/sequenceTools). For C/T and G/A SNPs, we used end-masked BAM files, and for the others we used the original unmasked BAM files. For the eight ssDNA libraries, we used end-masked BAM files for C/T SNPs, and the original BAM files for the others.

In cases where more than one sample was genetically analyzed per individual, we compared the amount of human DNA between samples. For pairs of petrous bone and teeth, human DNA was higher in the petrous bone in 8 of 13 individuals, and higher in the teeth of 5 of 13 individuals (Table S5). In addition, intra-individual sample variation was high, as evidenced by the high variance observed between paired tooth samples (Table S5). Finally, in a comparison of dsDNA and ssDNA libraries, ssDNA libraries yielded a higher endogenous content in 7 of 8 library pairs. All data from paired samples were merged prior to further analysis.

Of the 225 new individuals analyzed, 18 failed to yield sufficient human DNA (>0.1%) on shotgun screening (Table S4) and a further 6 individuals failed to yield at least 10,000 SNPs after DNA capture (Table S4). Both were excluded from downstream population genetic analysis.

#### *6.2 Data quality authentication*

To confirm that our sequence data consist of endogenous genomic DNA from ancient individuals with minimal contamination, we collected multiple data quality statistics. First, we tabulated 5' C>T and 3' G>A misincorporation rate (Fig. S2) as a function of position on the read using mapDamage v2.0.6 (Jónsson et al., 2013). Such misincorporation patterns, enriched at the ends due to cytosine deamination in degraded DNA, are considered as a signature of the presence of ancient DNA in large quantities (Sawyer et al., 2012). Second, we estimated mitochondrial DNA contamination for all individuals using the Schmutzi program (Renaud et al., 2015). Specifically, we mapped adapter-removed reads to the revised Cambridge Reference Sequence of the human mitochondrial genome (rCRS; NC_012920.1), with an extension of 500 bp at the end to preserve reads passing through the origin. We then wrapped the alignment to the circular reference genome using circularmapper v1.1 (Peltzer et al., 2016). The contDeam and schmutzi modules of the Schmutzi program were successively run with the world-wide allele frequency database from 197 individuals, resulting in estimated mitochondrial DNA contamination rates for each individual (Table S4). Last, for males, we also estimated the nuclear contamination rate (Table S4) based on X chromosome data using the contamination module in ANGSD v0.910 (Korneliussen et al., 2014). For this analysis, an increased mismatch rate in known SNPs compared to that in the flanking bases is interpreted as the evidence of contamination because males only have a single copy of the X chromosome and thus their X chromosome sequence should not contain polymorphisms. We report the Method of Moments estimates using the "method 1 and new likelihood estimate", but all the other estimates provide qualitatively similar results.

Ten individuals were estimated to have >5% DNA contamination (mitochondrial or X) or uncertain genetic sex (Table S4); these individuals were excluded from downstream population genetic analysis.

#### *6.3 Genetic sex typing*

We calculated the genetic sequence coverage on the autosomes and on each sex chromosome in order to obtain the ratio between the sex chromosome coverage and the autosome coverage. For 1240K capture data, we observe females to have an approximately even ratio of X to autosomal coverage (X-ratio of ~0.8) and a Y-ratio of 0, and males to have approximately half the coverage on X and Y as autosomes (~0.4). Genetic sex could be determined for a total of 224 individuals, of which 100 were female and 124 were male (Table S6).

#### *6.4 Uniparental haplogroup assignment*

We called mitochondrial consensus sequence from the Schmutzi output using the log2fasta program in the Schmutzi package, with quality threshold of 10. We then assigned each consensus sequence into a haplogroup (Table S6) using the HaploGrep 2 v2.1.19 (Weissensteiner et al., 2016). For the Y haplogroup assignment, we took 13,508 Y chromosome SNPs listed in the ISOGG database and made a majority haploid genotype call for each male using pileupCaller (with "-m MajorityCalling" option). We assigned each individual into a haplogroup (Table S6) using a patched version of the yHaplo program (Poznik, 2016) downloaded from https://github.com/alexhbnr/yhaplo. This version takes into account high missing rate of aDNA data to prevent the program from stopping its root-to-tip haplogroup search prematurely at an internal branch due to missing SNP and therefore assigning a wrong haplogroup. We used "--ancStopThresh 10" following the developer's recommendation. Haplogroup assignments are shown in Fig. S3.

#### *6.5 Estimation of genetic relatedness*

To evaluate the relatedness within our dataset, we calculated pairwise mismatch rate of haploid genotypes on automosomes across all individuals. The pairwise mismatch rate for each pair of individuals, is defined as the number of sites where two individuals have different alleles sampled divided by the total number of sites that both individuals have data. The pairwise mismatch rate between unrelated individuals is set as the baseline and the coefficient of relationship is inversely linear to the baseline pairwise mismatch rate. More detailed description can be found in the Supplemental Materials of (Jeong et al., 2018).

A total of 15 first or second degree genetic relationships were observed across the dataset (Table S7), of which 10 date to the Xiongnu era. Additionally, in one case, a tooth and petrosal bone thought to belong to one individual (AT-871) were later discovered to belong to two different individuals (OLN001.A and OLN001.B). In another case, two teeth (AT-728 and AT-729) thought to belong to different individuals were found to originate from the same individual (TUK001/TAV008) .

#### *6.6 Data filtering and compilation for population genetic analysis*

To analyse our dataset in the context of known ancient and modern genetic diversity, we merged it with previous published modern genomic data from i) 225 worldwide populations genotyped on the Human Origins array (Jeong et al., 2019; Lazaridis et al., 2014), ii) 300 high-coverage genomes in the Simons Genome Diversity Project ("SGDP") (Mallick et al., 2016), and currently available ancient genomic data across Eurasian continent (Allentoft et al., 2015; de Barros Damgaard et al., 2018; Damgaard et al., 2018; Fu et al., 2014, 2016; Haak et al., 2015; Haber et al., 2017; Harney et al., 2018; Jeong et al., 2016, 2018; Jones et al., 2015; Kılınç et al., 2016; Lazaridis et al., 2016, 2017; Mathieson et al., 2015, 2018; McColl et al., 2018; Narasimhan et al., 2019; Raghavan et al., 2014, 2015; Rasmussen et al., 2010, 2014, 2015; Sikora et al., 2019; Unterländer et al., 2017; Yang et al., 2017). We obtained 1,233,013 SNP sites (1,150,639 of which on autosomes) across our dataset when intersecting with the SGDP dataset, and 597,573 sites (593,124 of which on autosomes) when intersecting with the Human Origins array.

#### *6.7 Analysis of population structure and relationships*

We performed principal component analysis (PCA) on the merged dataset with the Human Origins data using the smartpca v16000 in the Eigensoft v7.2.1 package (Patterson et al., 2006). Modern individuals were used for calculating PCs (Fig. S4), and ancient individuals were projected onto the pre-calculated components using "lsqproject: YES" option (Fig. 2; Fig. S5). To characterize population structure further, we also calculated *f3* and *f4* statistics using qp3Pop v435 and qpDstat v755 in the admixtools v5.1 package (Patterson et al., 2012). We added "*f4mode: YES*" option to the parameter file for calculating *f4* statistics. We used DATES (Narasimhan et al., 2019) for dating admixture in the different ancient population groups.

#### *6.8 Admixture modeling using qpAdm*

For modelling admixture and estimating ancestry proportions, we applied qpWave v410 and qpAdm v810 in the the admixtools v5.1 package (Patterson et al., 2012) on the merged dataset with the SGDP data to maximise resolution. To model the target as a mixture of the other source populations, qpAdm utilizes the linearity of *f4* statistics, i.e. one can find a linear combination of the sources that is symmetrically related to the target in terms of their relationship to all outgroups in the analysis. qpAdm optimizes the admixture coefficients to match the observed f4 statistics matrix, and reports a *p*-value for the null hypothesis that the target derives their ancestry from the chosen sources that are differently related to the outgroups (i.e., when $p<0.05$, the null hypothesis is rejected so that the target is different from the admixture of chosen sources given the current set of outgroups). The chosen outgroups in qpAdm needs to be differentially related to the sources such that a certain major ancestry is "anchored" in the test, which is rather heuristic. We used qpWave to test the resolution of a set of outgroups for distinguishing major ancestries among Eurasians, as well as the genetic cladility between populations given a set of outgroups. We used a set of eight outgroup populations in our study: Central African hunter-gatherers Mbuti.DG (n=5), indigenous Andamanese islanders Onge.DG (n=2), Taiwanese Aborigines Ami.DG (n=2), Native Americans Mixe.DG (n=3), early Holocene Levantine hunter-gatherers Natufian (n=6) (Lazaridis et al., 2016), early Neolithic Iranians Iran_N (n=8) (Lazaridis et al., 2016; Narasimhan et al., 2019), early Neolithic farmers from western Anatolia Anatolia_N (n=23) (Mathieson et al., 2015), and a Pleistocene European hunter-gatherer from northern Italy Villabruna (n=1) (Fu et al., 2016).

To evaluate potential sex bias (Fig. S6), we applied qpAdm to both the autosomes (default setting) and the X chromosome (adding "chrom:23" to the parameter file) for comparing the difference in the estimated ancestry porportions. For a certain ancestry, we calculated sex-bias Z score using the proportion difference between $P_A$ and $P_X$ divided by their standard errors ($Z = \frac{P_A - P_X}{\sqrt{\sigma_A^2 + \sigma_X^2}}$, where $\sigma_A$ and $\sigma_X$ are the corresponding jackknife standard errors, as previously performed in (Mathieson et al., 2018). Therefore a positive Z score suggests autosomes harbor a certain ancestry more than X chromosomes do, indicating male-driven admixture. A negative Z score, in contrast, suggests female-driven admixture.

#### *6.9 Phenotypic SNP analyses*

We examined 49 SNPs in 17 genes (Table S8) known to be associated with phenotypic traits or with positive selection in Eurasia (Jeong et al., 2018). Given the low coverage of ancient DNA data, we focused on five of these genes and calculated the likelihood of allele frequency for SNPs in each ancient population based on the counts of reads covering on the SNP following a published strategy (Mathieson et al., 2015). In the allele frequency calculation, we classified all ancient individuals before Middle/Late Bronze Age into a single group, and kept three genetic groups during MLBA (Khövsgöl_LBA, Altai_MLBA, Ulaanzuukh), two genetic groups during Iron Age (Chandman_IA, SlabGrave), one group for Xiongnu, one group for Early Medieval and one group for Late Medieval. We calculated allele frequency at five loci (Table S8) that are associated with lactase persistence (LCT/*MCM6*), skin pigmentation (*OCA2*, *SLC24A5*), alcohol metabolism (*ADH1B*), and epithelial phenotypes including shovel-shaped incisor (*EDAR*) (Fig. 4).

**7. Genetic clustering of ancient individuals into analysis units**

To further characterize the dynamic changes of the Eastern Steppe gene pools using group-based analyses, we quantitatively examined genetic differences among the analyzed individuals in combination with their temporal, archeological, and geographic information. We first obtained an approximate map of population structure by observing the position of ancient individuals on the PCA calculated from 2,077 present-day Eurasian individuals. PC1 separates geographically eastern and the western populations, PC2 captures the internal variations in eastern Eurasians, and PC3 captures variations in western Eurasians, thus allowing us to characterize an overall pattern of genetic changes through time and helping us to formulate explicit hypotheses regarding the genetic relationships between groups and individuals. Second, we computed outgroup-*f3* and symmetric-*f4* statistics to (1) quantify genetic similarity between individuals/groups falling together on PCA and (2) explore populations whose ancestry through admixture may have contributed to the differences observed between pairs of groups. Third, we identified representative ancient populations to serve as proxies for five distinct ancestries that we then further investigate (Table S10). We changed the specific ancestry proxy for our test groups based on the temporal and archeological records accordingly. Using these ancestry proxies, we performed a formal admixture modeling using qpWave/qpAdm, which tests difference between the target and a combination of the proxies (i.e. an admixture model) with regard to their genetic affinity to outgroups. We applied the same admixture models for test groups belonging to the same time/culture/geography category to compare them in a straightforward manner (Fig. 3; Fig. S6). In the following paragraphs, we describe each of the genetics-based analysis groups reported in our dataset, as well as the principles we applied to model their genetic ancestry using qpAdm.

***7.1 Pre-Bronze Age***

- New genetic groups: eastMongolia_preBA(1), centralMongolia_preBA(1), and Fofonovo_EN(4)
- Published genetic groups: DevilsCave_N(6), and Baikal_EN(9)

Our dataset adds three Ancient Northeast Asian (ANA)-related genetic groups before the start of the Bronze Age in eastern Eurasia. During this period, we observe the wide distribution of this ANA ancestry from Lake Baikal to the Russian Far East, spanning more than 2,000 kilometers. As Baikal_EN has been modeled to have ~10% Ancient North Eurasian (ANE) ancestry, we also investigated the possible genetic contribution from ANE in our pre-Bronze Age Mongolian and Baikal groups using Botai, AG3, MA1 and West_Siberia_N separately as ancestry proxies. We find ANE-related ancestry appears in centralMongolia_preBA and Fofonovo_EN only to a minor extent; ANE ancestry is not present in eastMongolia_preBA, which is instead characterized by only ANA-related ancestry (Table S13).

#### *7.2 Early Bronze Age*

- New genetic groups: Afanaseivo_Mongolia(2), Afanasievo_KUR001(1), Chemurchek_Altai(2), and Chemurchek_KUM001(1)
- Published genetic groups: Afanaseivo(23), Okunevo_EMBA(19), and Baikal_EBA(5)

Our dataset adds two main genetic groups during Early Bronze Age - Afanasievo_Mongolia and Chemurchek_Altai. We group two individuals from Shatar Chuluu site (SHT001, SHT002) into Afanasievo_Mongolia as both are archaeologically classified into the Afanasievo cultural context and genetically indistinguishable from Afanasievo individuals from the Russian Altai-Sayan region (Table S14, Figure S9; S11). One female Afanasievo individual from Altai (KUR001) shows mainly ANA-related ancestry with small proportions of ancestry related to Afanasievo_Mongolia (Table S14, Figure S9; S11), and consequently we have grouped her separately as Afanasievo_KUR001. She appears to be an individual of mixed local ANA and migrant Afanasievo ancestry who was buried in a culturally Afanasievo manner.

We group two individuals from Yagshiin Huduu site (IAG001, YAG001) into Chemurchek_Altai as both are archaeologically classified to the Chemurchek cultural context and cluster together on PCA, providing the first genomic investigation of the Chemurchek culture. We observed that Chemurchek_Altai has the highest genetic affinity to ANE-related groups (e.g., Botai) and secondary affinity to Iranian-related groups (Figure S9; S11). We tested Afanasievo as a potential ancestral source given the geographic overlap and similar burial posture between Afanasievo and Chemurchek (Taylor et al., 2019), however the 2-way model with Afanasievo as one of the two sources fails (Table S15). The model also fails when using Okunovo (a neighboring group contemporaneous with Chemurchek that succeeds the Afanasievo culture) either as Afanasievo + Okunevo or Okunevo + Iranian (Table S15). Thus, despite some cultural similarities between steppe groups (Afanasievo, Okunevo) and Chemurchek, there is no evidence for the genetic influence from the steppe populations among the Chemurchek individuals analyzed here. To further investigate the Iranian-related ancestry among the Chemurchek, we tested four published groups from the BMAC genetic cluster (Gonur1_BA, Bustan_BA, Dzharkutan1_BA, and Sappali_Tepe_BA), four Chalcolithic/Bronze Age Iranian groups (Hajji_Firuz_C, Tepe_Hissar_C, Seh_Gabi_C, and Shahr_I_Sokhta_BA1), Eneolithic Turkmenistan and Tajikstan groups (Paikhai_EN, Sarazm_EN, and Tepe_Anau_EN) and Mesolithic Caucasus Hunter-Gatherer (CHG). Interestingly, 2-way models consisting of Botai + BMAC genetic cluster groups adequately model Chemurchek_Altai with ~40% ancestry proportion from the latter, while preceding Eneolithic Turkmenistan/Tajikistan groups do not (Table S15). Spatiotemporally more distant groups, such as Chalcolithic Iranian groups or CHG, also adequately model Chemuchek with similar ancestry proportion (Table S15). We find that Chemurchek_Altai has the closest genetic affinity to Dali_EBA (Fig S9), an individual dating to ca. 2650 BCE with poor burial context from southeastern Kazakhstan who has admixed ANE-Iranian ancestry (see (Narasimhan et al., 2019)). Applying the same 2-way admixture models using Dali_EBA for comparison, we found that Dali_EBA also requires an additional Iranian-related ancestry but in a smaller proportion. In addition to the main Chemurchek genetic cluster, one female Chemurchek individual (KUM001) from the Altai shows high genetic affinity to ANA groups with only small ancestral proportions from Chemurchek (Table S15, Figure S9; S11). Because she appears to be an individual of mixed local ANA and some Chemurchek_Altai ancestry who was buried in a culturally Chemurchek manner, we therefore classified her into a separate group: Chemurchek_KUM001.

#### *7.3 Middle and Late Bronze Age*

- New genetic groups: Altai_MLBA(7), Ulaanzuukh_SlabGrave(11/16), UAA001(1), KHI001(1), UUS001(1), KHU001(1) and TSI001(1)
- Published genetic groups: Khövsgöl_LBA(17), ARS017(1), ARS026(1), Sintashta_MLBA(37), Krasnoyarsk_MLBA(18)

Our dataset adds two main genetic groups in the Eastern Steppe during the MLBA - Altai_MLBA and Ulaanzuukh_SlabGrave - to the previously published Khövsgöl_LBA from northern Mongolia (Jeong et al., 2018). Our new data substantially expand the geographic scope of genetically characterized MLBA populations in Mongolia, and reveal an overall picture of the population structure of the MLBA Eastern Steppe. The Altai_MLBA group contains seven individuals from the Altai-Sayan region (BER002, BIL001, ULZ001, ARS026, SBG001, ULI001, ULI003), who are admixed between the Western Steppe gene pool associated with Srubnaya/Sintashta/Andronovo cultures ("steppe_MLBA") and the one associated with Khövsgöl_LBA/Baikal_EBA. Although the ancestry proportion estimates within this group vary along a cline, the Altai_MLBA represents the formation of a gene pool incorporating a substantial genetic influx from Western Steppe herders. Thus we classified them into one genetic group despite their archaeological and cultural differences (Mönkhkhairkhan, Baitag, and DSKC burial types). This also explains the genetic profile of one outlier from Khövsgöl_LBA (ARS026), who now genetically falls within the Altai_MLBA group. Of note, one member of this group, ULZ001, is found not in the Altai, but in far eastern Mongolia.

The other genetic cluster - Ulaanzuukh_SlabGrave, contains 11 individuals with Ulaanzuukh burial type (BUL001, BUL002, ULN001, ULN002, ULN003, ULN005, ULN006, ULN007, ULN009, ULN010, ULN015) and 5 individuals with Slab Grave burials (see below), from eastern Mongolia. They all are classified into one single genetic group given their strong genetic homogeneity with ANA (Table S16) and the geographic links between the two. This clustering of Ulaanzuukh and Slab Grave confirms previous archaeological hypotheses that the Slab Grave culture likely emerged out of the Ulaanzuukh gene pool. This genetic cluster also explains another Khövsgöl_LBA outlier - ARS017, who now genetically falls within the Ulaanzuukh_SlabGrave group, as well as a single individual with unknown burial type from central Mongolia, TSI001, who also falls into this cluster. Of note, one male Mönkhkhairkhan individual (KHU001) also has a large proportion of ancestry from Ulaanzuukh_SlabGrave in addition to his main genetic component from Baikal_EBA (Table S16). Together, the individuals ARS017, TSI001, and KHU001 suggest contact with the Ulaanzuukh_SlabGrave group in northern, central Mongolia, even though these individuals were buried according to local burial customs. Overall, this Ulaanzuukh_SlabGrave genetic cluster is a continuation of the ANA easternMongolia_preBA gene pool (represented by SOU001) of 3,000 years earlier.

We also identified three outliers which do not fall into any of the three genetic clusters described above. UAA001 (Mönkkhairkhan) from the Altai is well-fitted with 3-way admixture model using Afanasievo, Baikal_EBA and Gonur1_BA (Table S16), despite the fact that they date to ~1500 years after the

Afanasievo culture. KHI001 (unclassified culture) from the Altai, is well-fitted with 3-way admixture model using Sintatash, Baikal_EBA and Gonur1_BA (*p*-value=0.056, Table S16), presenting minor genetic component from Gonur1_BA. Alternatively, KHI001 can also be modeled as a 2-way admixture between Afanasievo and Khövsgöl_LBA (*p*-value=0.117, Table S16); however, this model has a lower priority than the former model. UUS001 (DSKC) from Khövsgöl province is well-fitted with 3-way model using Sintashta, eastMongolia_preBA and Gonur1_BA (Table S16). Given the temporal discordance between UUS001 and the eastMongolia_preBA individual (~3,000 years), it is more likely that the admixing partner for UUS001 was related to the Ulaanzuukh cluster; Ulaanzuukh shares a high degree of ancestry with eastMongolia_preBA and is contemporaneous with the UUS001 individual, and some Ulaanzuukh individuals plot very close to the eastMongolia_preBA individual - SOU001 in PCA.

***7.4 Early Iron Age***

- New genetic groups: Chandman_IA(9), Ulaanzuukh_SlabGrave(5/16),
- Published genetic groups: Tagar(8), CentralSaka(6), TianShanSaka(10), Kazakhstan_Berel_IA(2; Pazyryk culture)

Our dataset adds two main genetic groups during EIA - one represented by Ulaanzuukh_SlabGrave and the other represented by the site of Chandman Mountain associated with the Uyuk culture (Chandman_IA). In addition to the 11 Ulaanzuukh burials described above, four Slab Grave individuals (BOR001, DAR001, MIT001, SHU001) from eastern Mongolia also presented a homogeneous genetic profile with Ulaanzuuk and thus were merged into the Ulaanzuukh_SlabGrave analysis group (Table S17). Interestingly, PTO001, a Trans-Baikal individual who is also archaeologically classified as Slab Grave, has a genetic profile that matches other Slab Grave individuals from eastern Mongolia, and we also merged PTO001 into the Ulaanzuukh_SlabGrave genetic cluster. The genetic profile of PTO001 is consistent with an archaeologically described expansion of the Slab Grave culture into the Baikal region during EIA (Losey et al., 2017).

The contemporaneous Chandman_IA from the Altai-Sayan region in western Mongolia has a genetic profile that matches the preceding Altai_MLBA cline. Since all individuals are from a single site and cluster together on PCA, we group them into a single analysis unit ("Chandman_IA"). Here, we use the Andronovo-associated dataset Krasnoyarsk_MLBA as the representative central steppe_MLBA group for admixture modelling because it is geographically closest to our test EIA groups. We first tested a 2-way admixture model of Krasnoyarsk_MLBA + Baikal_EBA, but it failed to adequately model the Chandmand_IA cluster, as did Karsnoyarsk_MLBA + Khövsgöl_LBA. Further changing the steppe_MLBA source from Karsnoyarsk_MLBA to Sintashta_MLBA did not rescue the 2-way admixture model. We then attempted a 3-way admixture model by adding Iranian-related ancestry as the third source, using a BMAC group from the Gonur Tepe site (Gonur1_BA) as a proxy. Using Krasnoyarsk_MLBA as the Steppe proxy, we observed 51.3% of Steppe, 42.2% of Baikal_EBA and 6.5% of Iranian ancestry in Chandman_IA (Table S17).

Because it is a priori quite unlikely due to a long-distance migration from Bactria/Iran specific to Chandman_IA, we next applied the same 3-way models of Krasnoyarsk_MLBA/Sintashta_MLBA+Baikal_EBA+Gonur1_BA to four Iron Age central Asian groups

(Tagar from Minusinsk Basin, Central Saka from central Kazakhstan, Kazakhstan_Berel_IA from eastern Kazakhstan, and Tian Shan Saka from Kyrgyzstan) and also to the Final Bronze Age group Karasuk. We observed that Iranian-related ancestry proportions range from ~7-28% in the tested Iron Age groups, while not required for Karasuk. In particular, the Tian Shan Saka, geographically closest to the Gonur Tepe site, has the highest amount of estimated Iranian-related ancestry. Because of cultural connections between the Uyuk of Chandman_IA and the Saka generally (see section 2.4 above), it is possible that Saka and related groups in Tian Sian, Fergana and Transoxiana/Turan (such as the sampled Tian Shan Saka) are the proximal source of the Iranian ancestry in the Iron Age groups further to the north, such as Chandman_IA. To narrow down the spatiotemporal origin of this Iranian-related ancestry, we tested 3-way models using alternative Iranian-related groups as the proxy in the Tian Shan Saka: (1) three other post-BMAC groups (Bustan_BA, Dzharkutan1_BA, and Sappali_Tepe_BA) that fall into the BMAC genetic cluster with Gonur1_BA (Narasimhan et al., 2019), (2) Shahr_I_Sokhta_BA1 from the southeastern corner of Iran, (3) three Chalcolithic Iranian groups (Hajji_Firuz_C, Tepe_Hissar_C, Seh_Gabi_C), (4) two Iron Age groups from Pakistan (Katelai_IA, Loebanr_IA), (5) Eneolithic groups from Turkmenistan such as Geoksyur_EN, Parkhai_EN and Tepe_Anau_EN, and (6) Sarazm_EN fromwestern Tajikistan. All Iranian-ancestry proxies mentioned above except Hajji_Firuz_C and Seh_Gabi_C from the Zagros provide a well-fitted 3-way model (Table S18). Therefore, for the Iron Age Eastern Steppe, genetic data alone can only narrow down the source of the Iranian ancestry to a broad region east of the Caspian Sea. Taken in context, though, we propose that this ancestry likely arrived via a local contact around the Transoxiana/Sogdiana region (i.e., the border between Kazakhstan, Uzbekistan and Kyrgyzstan).

For the prehistoric genetic groups described above, we used DATES to estimate the date of admixture between Western ancestry sources (WSH or the Iranian-related groups) and local ancestry sources (i.e., Khovsgol_LBA or Baikal_EBA) (Fig. S12, Fig. S13). As shown in Fig. S12, the estimated admixture date between Sintashta and Baikal_EBA for the Karasuk and Tagar is consistent with the admixture date observed in Altai_MLBA - at around 3,500 BP. For the Central Saka, Pazyryk (Kazakstan_Berel_IA) and Uyuk (Chandman_IA), the admixture date is estimated to be a few centuries later, and the most recent admixture date is estimated for the Saka from Tian Shan. Notably, we find that the estimated admixture dates between Gonur1_BA and Baikal_EBA in the Iron Age groups are roughly consistent with the admixture dates for Sintashta (Fig. S12). However, because we are using a method designed for dating a 2-way admixture on what is best modeled as 3-way admixture in our study, we caution that these admixture dates should be interpreted with care.

#### *7.5 Xiongnu Empire*

- New genetic groups: earlyXiongnu_west(6), earlyXiongnu_rest(6), SKT007(1), lateXiongnu(24), lateXiongnu_sarmatian(13), lateXiongu_han(8)
- Published genetic groups: Xiongnu_WE(2), Xiongnu_royal(1, DA39.SG), Han_2000BP(2)

Our dataset reveals a great deal of previously uncharacterized genetic diversity during the Xiongnu period. For individual modelling, we tested every possible combination of five main ancestries: Steppe (Krasnoyarsk_MLBA, Sintashta, Srubnaya, Sarmatian, Chandman_IA), Gonur1_BA, Khövsgöl_LBA, Ulaanzuukh_SlabGrave, and Han. Considering the low resolution of individual modelling, we report selected working models that work for many individuals belonging to the same time period and archaeological context and that reflect qualitative trends observed in PCA. We observed that Iron Age Chandman_IA is a good Steppe ancestry proxy for many Xiongnu individuals, but there are also many who have western Eurasian ancestry in higher proportion than that of Chandman_IA. These individuals with high western Eurasian ancestry proportion show strong affinity to the Iranian-related ancestry that cannot be explained by the earlier Late Bronze Age steppe groups (e.g. Krasnoyarsk_MLBA, Sintashta_MLBA or Srubnaya). Instead, Gonur1_BA or Iron Age Sarmatian fit better with the genetic profile required. Also, a few individuals fall into the eastern Eurasian cline along PC2 and are explained as a combination of the eastern Eurasian gene pools, Ulaanzuukh_SlabGrave and present-day Han Chinese, without contribution from western Eurasian sources (Table S19). We used high-coverage whole genome sequences of present-day Han Chinese ("Han.DG"; n=4) as a proxy for the ancestry component that is currently broadly distributed across northern China and distinct from the component represented by Ulaanzuukh_SlabGrave further to the north. This is to achieve statistical power in our admixture modeling given that there are to date very few available ancient genomes that reflect this ancestry component. This is due to the fact that ancient China, Korea, Japan, and Southeast Asia remain mostly unsampled. We fully acknowledge the genetic diversity present within contemporary Han Chinese populations, and do not intend to claim by our admixture modeling a specific connection between the ancient populations within our study and present-day ethno-cultural identities.

For the group-based qpAdm modelling, we split Xiongnu into two categories based on their age - early Xiongnu and late Xiongnu. We further split early Xiongnu into two subgroups, earlyXiongnu_west (SKT010, SKT001, SKT003, SKT009, SKT008, AST001) and earlyXiongnu_rest (JAG001, SKT002, SKT004, SKT005, SKT006, SKT012), based on their individual modelling results, leaving out one individual outlier - SKT007 (Khövsgöl_LBA-like). The two previously published Xiongnu individuals grouped as "Xiongnu_WE" show a similar genetic profile to earlyXiongnu_rest, are dated to the early Xiongnu period, and are from the same valley as the two early Xiongnu sites (SKT and AST) in our dataset (Table S19). For the late Xiongnu, we summarized their individual modelling results in Table S20. Based on the individual modeling results, we set up three subgroups within late Xiongnu individuals to highlight key demographic processes and to use them for specific analyses such as sex-biased gene flow. First, we assigned 24 of 47 individuals into the main lateXiongnu group (BTO001, CHN010, DEL001, DOL001, IMA001-IMA008, JAA001, KHO006, KHO00, SAN001, SOL001, TEV002, TEV003, TUK003, UGU004, UGU011, ULN004, UVG001; Table S20); this group is well modeled as a mixture of two main Iron Age clusters, Chandman_IA+Ulaanzuuk_SlabGrave ($p=0.316$; 76.6±0.8% from Ulaanzuuk_SlabGrave). Another 13 individuals have more western Eurasian ancestry than Chandman_IA and thus require a different western Eurasian source. Two of them (NAI002, BUR001) are explained by Chandman_IA+Gonur1_BA, a model for earlyXiongnu_west, but the remaining 11 need Sarmatian contribution, including three that are cladal to Sarmatian (BUR003, TM001, UGU010). Taken all 13 individuals as a group (lateXiongnu_sarmatian; BRL002, BUR001-BUR004, DUU001, HUD001, NAI001, NAI002, TMI001, UGU005, UGU006, UGU010), we infer a major contribution from a Sarmatian-related source into this group (75.7±2.8%; Table S20). On the other hand, we grouped eight individuals (ATS001, BAM001, BRU001, EME002, SON001, TUH001, TUH002, YUR001) into the third group lateXiongnu_han based on their affinity to Han Chinese and other East Asian populations that Ulaanzuuk_SlabGrave cannot explain (37.2±10.6% from Han.DG; Table S20). The previously published Xiongnu_royal individual shows substantial Han-related ancestry (Table S19), similar to our lateXiongnu_han group. Further, the late Xiongnu individual YUR001 is an extreme East Asian outlier, who genetically resembles "Han_2000BP", two Han empire soldiers recovered from a mass grave near a Han fortress in the southern Gobi (Damgaard et al., 2018). These two groups, lateXiongnu_sarmatian and lateXiongnu_han, robustly support influxes of new ancestries both from the west and the east that were not previously observed in early Xiongnu or earlier populations. We left two individuals out of grouping, due to their unusual ancestry profiles: TAK001 mostly resembles Khövsgöl_LBA, and TUK002 is modeled as Chandman_IA+Ulaanzuuk_SlabGrave_Gonur1_BA (Table S20). In contrast to the strong east-west genetic division among Bronze Age Eastern Steppe populations through the end of the Early Iron Age, the Xiongnu period is characterized by an extreme degree of genetic diversity and heterogeneity that does not have any obvious geographic correlation (Fig. S14).

#### *7.6 Early Medieval*

- New genetic groups: TUK001(1), earlyMed_Türk(7), TUM001(1), earlyMed_Uyghur(12), OLN007(1)

Our dataset adds two main genetic groups during early Medieval in Mongolia - earlyMed_Türk and earlyMed_Uyghur. We tested every possible combination of four main ancestries - Steppe (Sarmatian, Alan), Gonur1_BA, Ulaanzuukh_SlabGrave, and Han for each individual. The genetic contribution from Iranian-related ancestry becomes even more prominent in Türkic and Uyghur individuals, as seen from well-fitted models using the Alan - an Iranian pastoral population from the Caucasus (Table S21). Overall, the Türkic and Uyghur individuals in this study show a high degree of genetic diversity, as seen in their wide scatter across PC1 in Fig. 2. TUK001(250-383 CE), the earliest early Medieval individual in our dataset from a Xiongnu site with a post-Xiongnu occupation, has the highest western Eurasian affinity. This individual is distinct from Sarmatians, and likely to be admixed between Sarmatians and populations with BMAC/Iranian-related ancestry (Table S1). Among the Türkic period individuals, TUM001 is a genetic outlier with mostly East Asian (Han_2000BP-like) ancestry. This individual was buried together with a knife and two dogs within the ramp of a Türkic era mausoleum. The mausoleum's stone epitaph indicates that it was constructed for a diplomatic emissary of the Pugu tribe who was allegiant to the Chinese Tang Empire. His cremated remains were found within the tomb; TUM001 was likely this emissary's servant (Ochir et al., 2013).

With respect to Uyghur burials, many consist of collective graves, and it has been suggested that such graves may contain the remains of kin groups (Erdenebat, 2016). We examined one such collective grave (grave 19) at the site of Olon Dov; however, of the six individuals analyzed in grave 19, there were no first degree relatives (parent-offspring pairs or sibling), and only two individuals (OLN002 and OLN003) exhibited a second degree (avuncular, grandparent-grandchild, or half-sibling) relationship. One Uyghur individual (OLN007) had markedly higher proportions of Han-related East Asian ancestry that cannot be explained by Ulaanzuuk_SlabGrave, and therefore grouped separately from the other earlyMed_Uyghur individuals (Table S21).

#### *7.7. Late Medieval*

- New genetic groups: lateMed_Khitan(3), lateMed_Mongol(61), SHU002(1)

Our dataset adds two main genetic groups during late Medieval in Mongolia: lateMed_Khitan and lateMed_Mongol. We used the same modelling strategy as used for the early Medieval period, and additionally explored the genetic cladality between every individual from the Mongol period and from modern Mongolic-speaking populations via qpWave (Table S23, Fig. S15). Relatively few Khitan individuals (n=3) were available for analysis, but all show high ANA-related ancestry (Table S22). Mongol-era individuals (n=61) are genetically more diverse and are cladal with modern Mongolic-speaking populations (Table S22, Fig. S15). SHU002 is single individual dated to late Medieval period however without recognizable Mongol-like burial feature. Overall, Mongol period individuals characterized by a remarkable decrease in Western Eurasian ancestry compared to the preceding 1,600 years. They are best modeled as a mixture of ANA-like and East Asian-like ancestry sources, with only minor Western genetic ancestry.

### Supplementary Figures

**a**

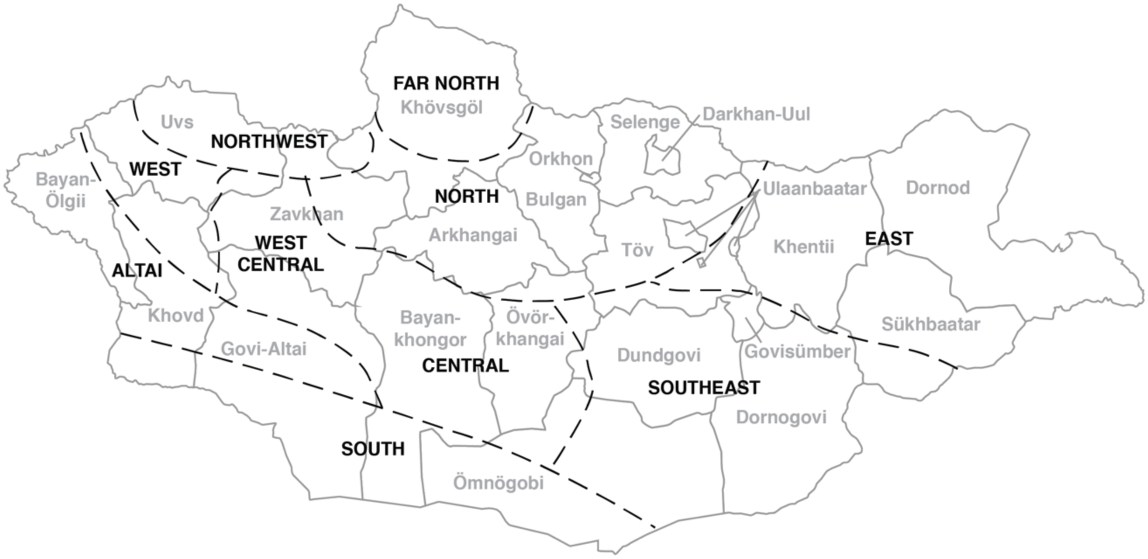

**b**

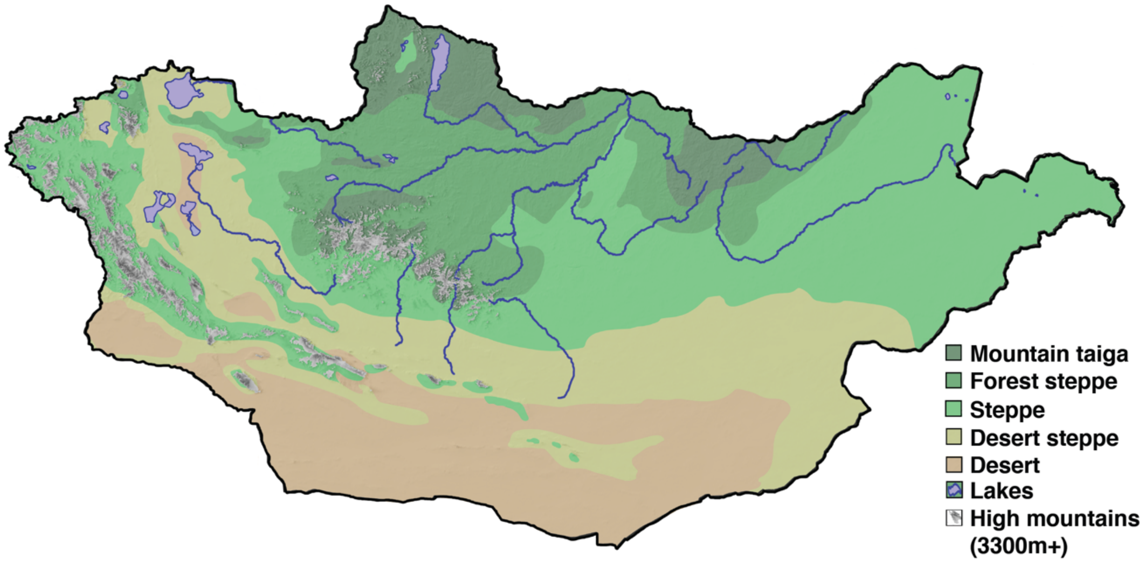

**Figure S1. Geographic and ecological features in Mongolia.** **a**, Mongolian regions and aimags (provicnes). Aimags are indicated by gray lines and text. Regions are indicated by black dashed lines and text following the definitions of Taylor et al. (2019). **b**, Ecological zones of Mongolia. Map produced using QGIS software (v3.6) with ecological data from Dorjgotiv (2004).

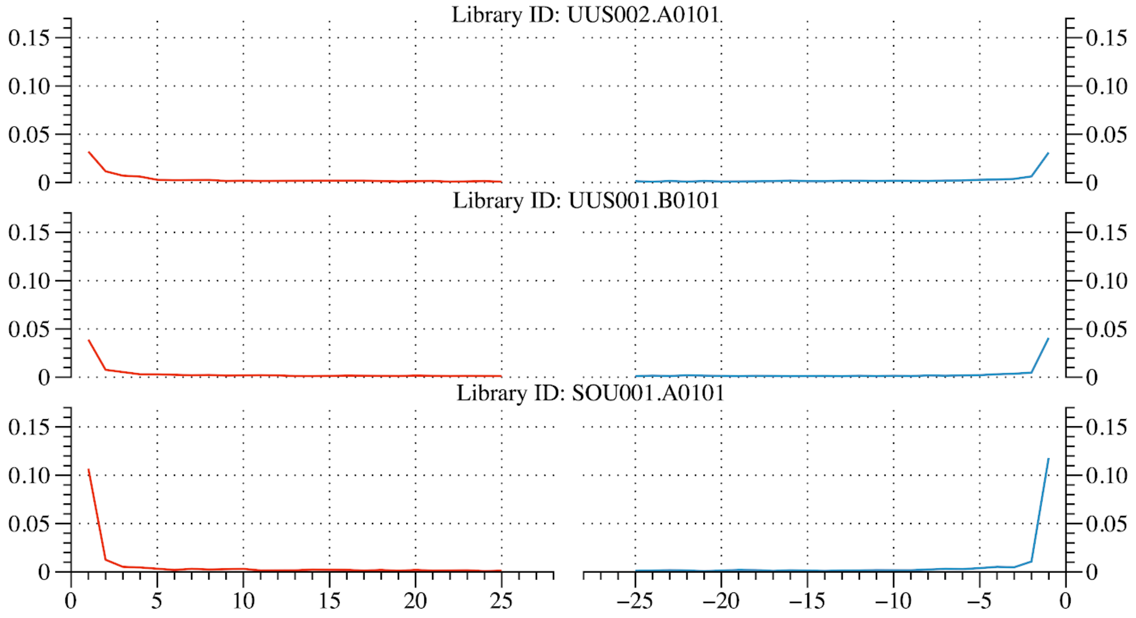

**Figure S2. Selected DNA damage patterns in ancient individuals.** Three individuals were selected as representative of the dataset: SOU001, Pre-Bronze Age; UUS001, DSKC culture, MLBA; UUS002, Mongol empire, Late Medieval. The level of DNA damage is measured by the rate of cytosine deamination-based misincorporation of bases as a function of position on reads. Red and blue lines represent C>T and G>A misincorporations, respectively. Data are shown for dsDNA UDG-half libraries. As expected from UDG-half treated libraries, damage is restricted to the first (C>T) and last (G>A) few bases. The observed damage patterns confirm that these individuals have typical patterns of DNA damage for archaeological samples.

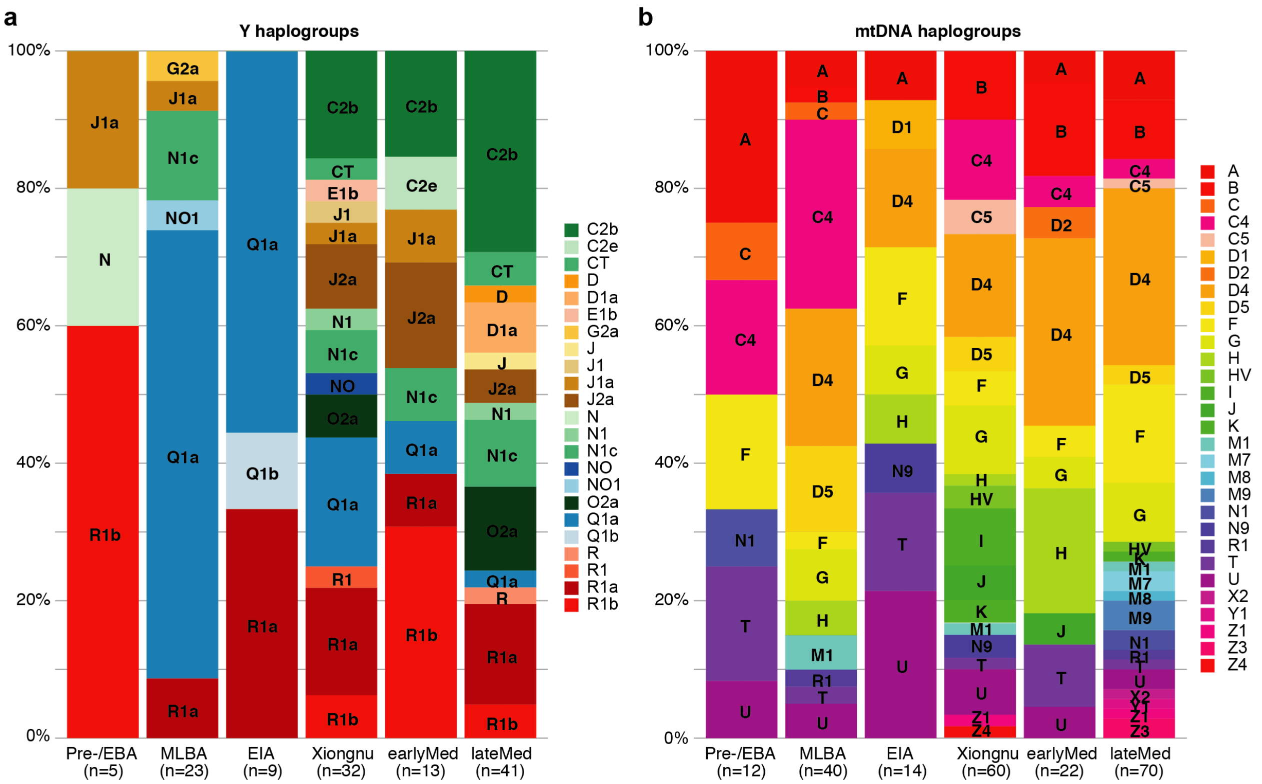

**Figure S3. Population structure from uniparentally inherited markers. a**, Distribution of Y haplogroups across each period. **b**, Distribution of mitochondrial haplogroups across each period.

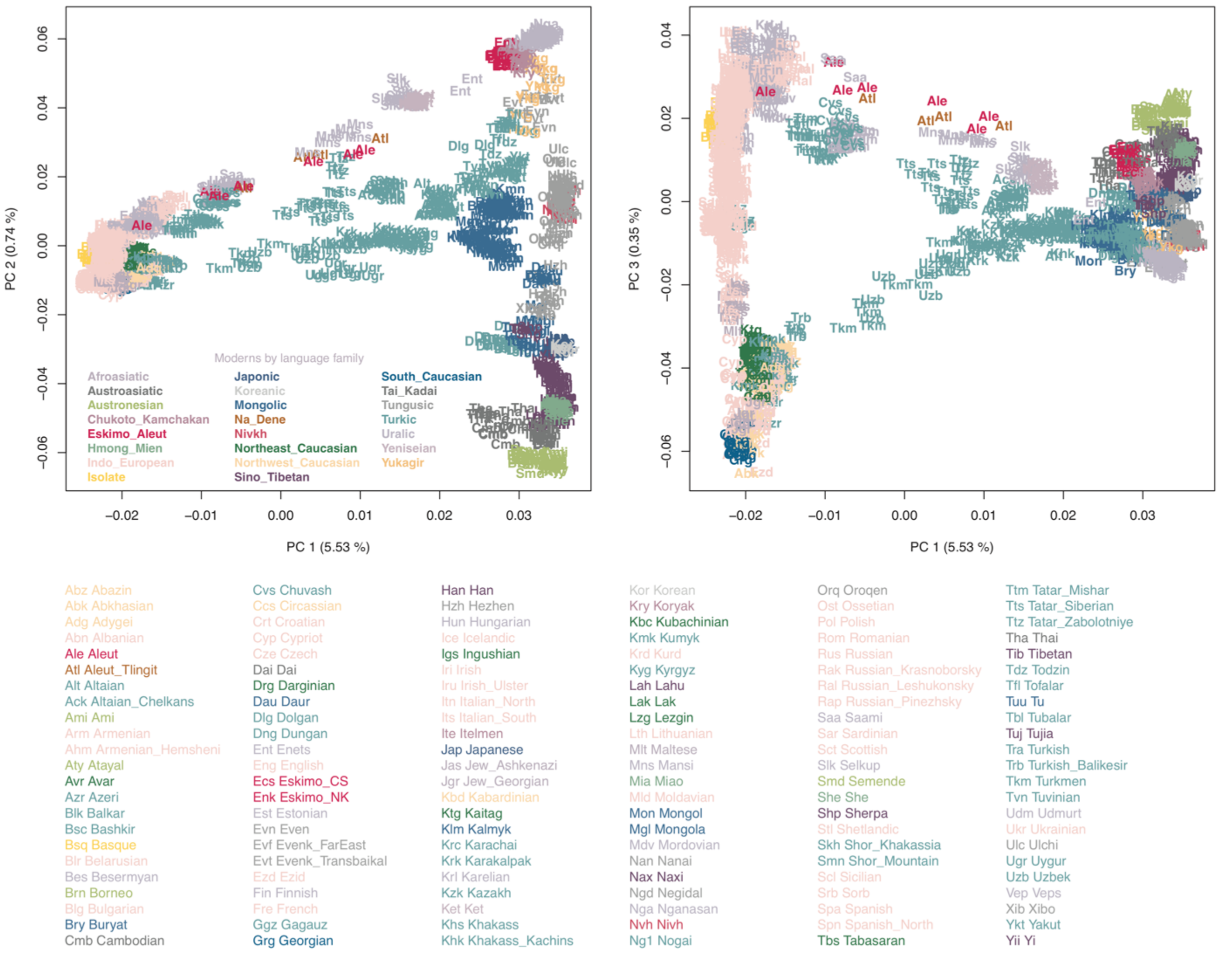

**Figure S4. PCA of present-day Eurasian populations used as the background for Fig. 2 and Fig. S5.** Here we show the population labels for the 2,077 Eurasian individuals used for calculating PCs and plotted as grey dots in Figure 2. Each three-letter code in the plot represents a single individual. Population IDs matching to the three-letter codes are listed at the bottom.

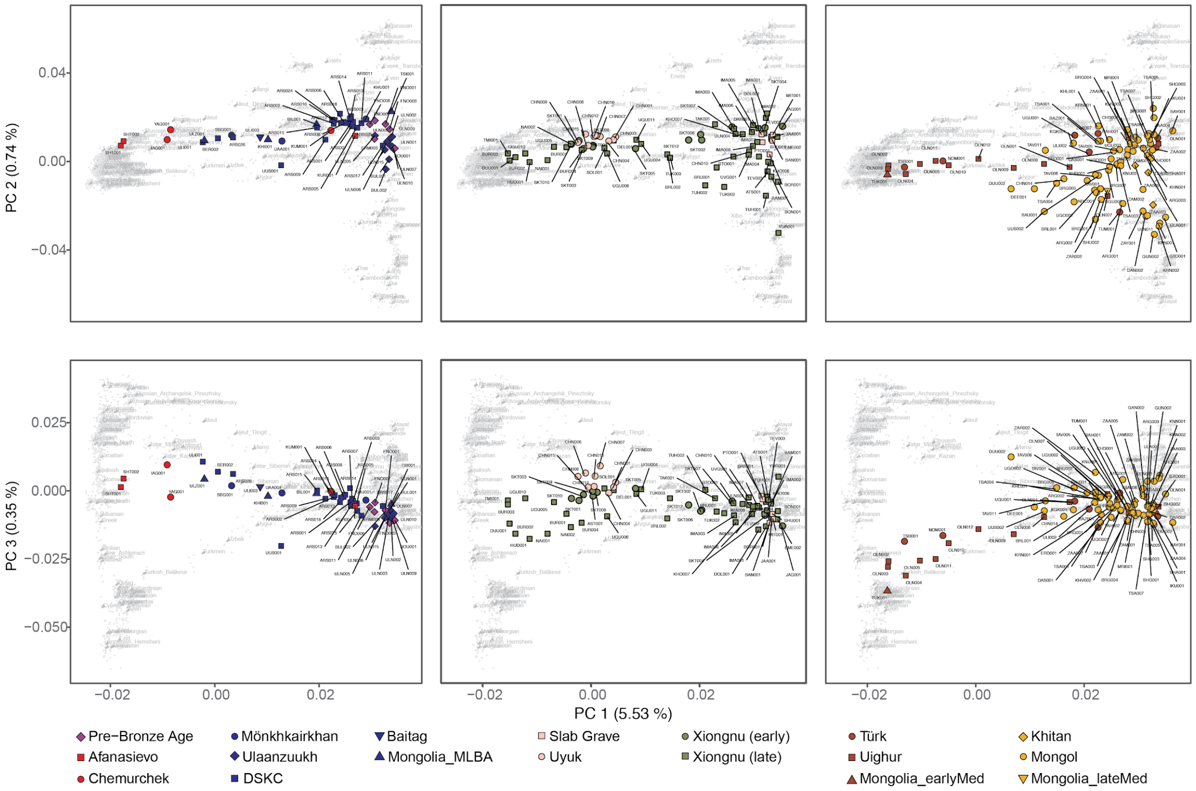

**Figure S5. Genetic structure of Mongolia through time. Principal component analysis (PCA) of ancient individuals (n=214) from three major periods projected onto contemporary Eurasians (gray symbols). a-c**, PC1 vs. PC2; **d-f**, PC1 vs PC3. **a,d**, Pre-Bronze to Late Bronze Age; **b,e**, Early Iron Age to Xiongnu; **c,f**, Early Medieval to Late Medieval. Projection and axis variance corresponds to Figure 2. Population labels are positioned over the mean coordinate across individuals belonging to each population.

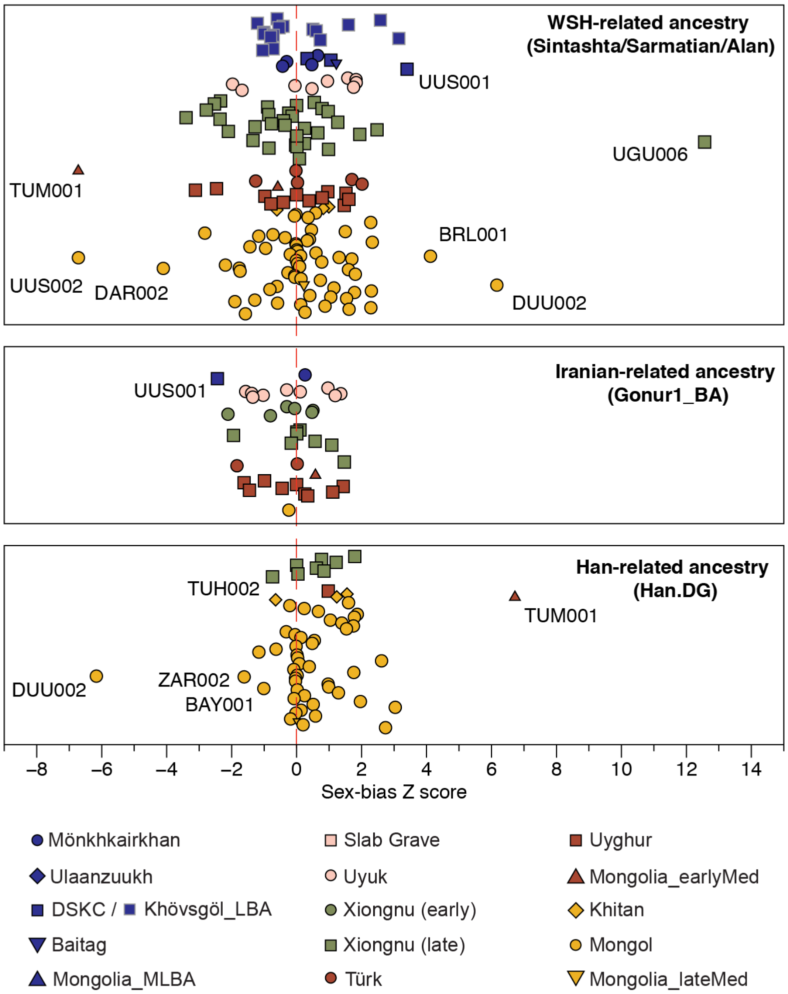

**Figure S6. Sex-bias Z scores by evaluating the differences of WSH-/Iranian-/Han-related ancestry on the autosomes and the X chromosome.** We calculated Z-score for every ancient individual who has genetic admixture with any of the three ancestries. Positive scores suggest more WSH-/Iranian-/Han-related ancestry on the autosomes, i.e., male-driven admixture.

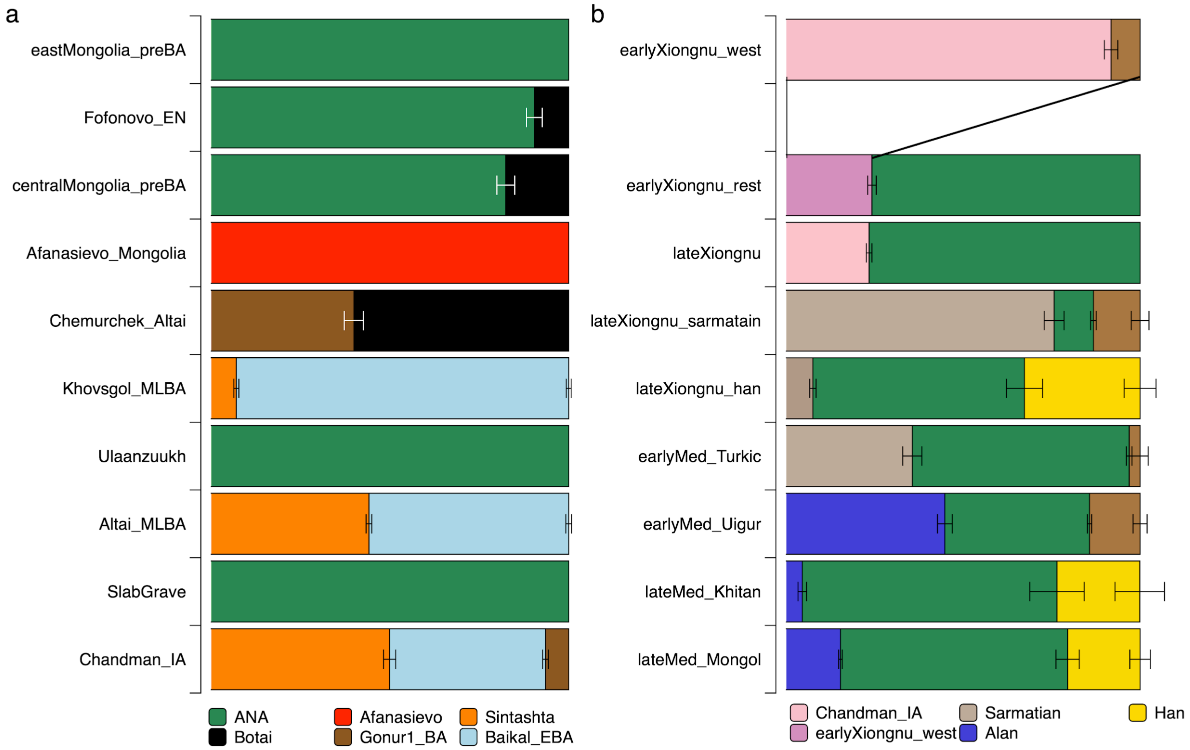

**Figure S7. Genetic ancestry changes in chronological order across all newly reported genetic groups.** We show the well-fitted modelling results in the grouped-based population genetics analyses for (**a**) prehistoric periods and (**b**) historic periods. The number of individuals in each group can be found on the right. Raw ancestry proportions and standard error estimates can be found in Tables S13-S22.

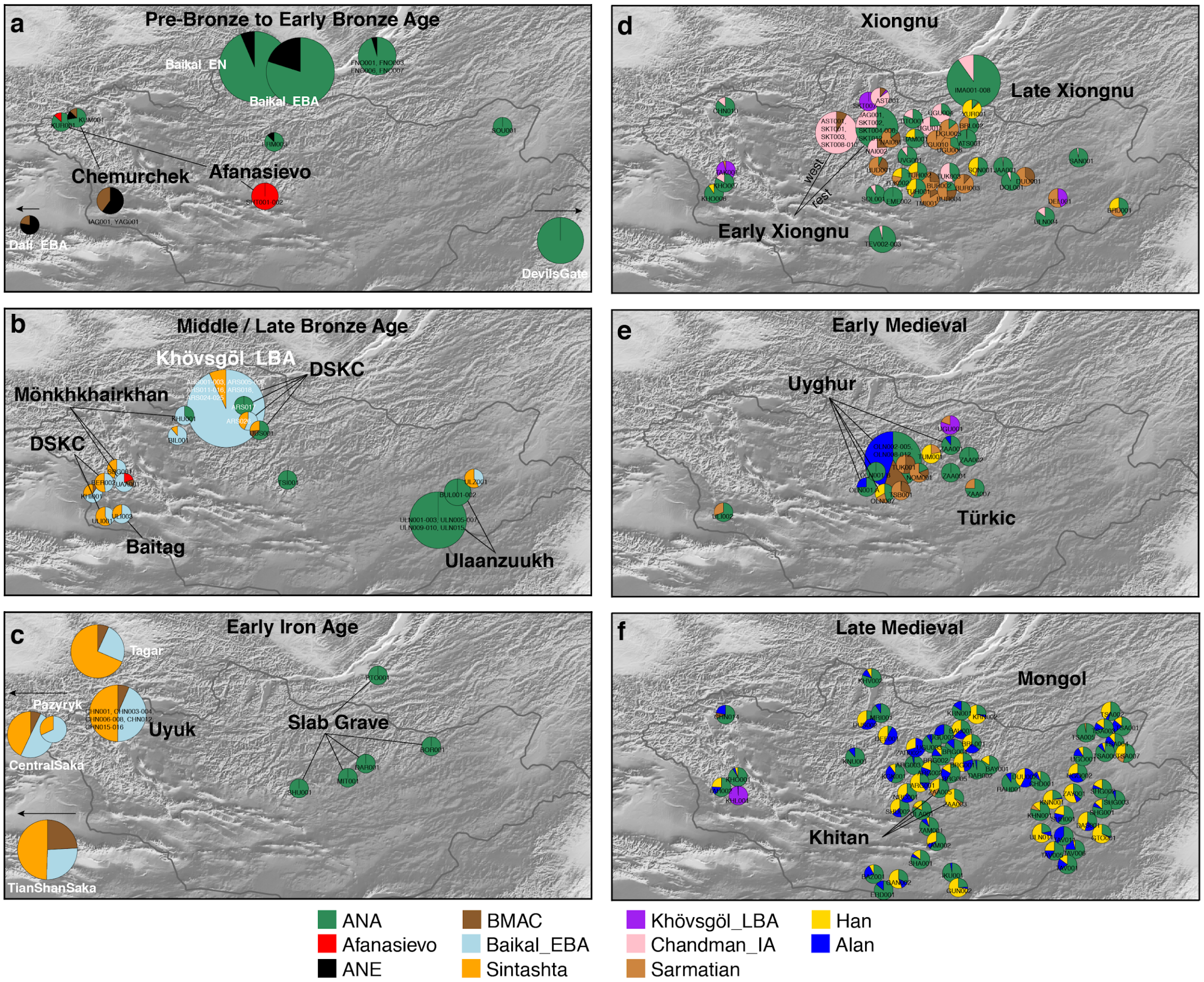

**Figure S8. Genetic changes in the Eastern Steppe across time characterized by qpAdm with all individuals indicated.** **a**, Pre-Bronze through Early Bronze Age; **b**, Middle/Late Bronze Age; **c**, Early Iron Age; **d**, Xiongnu period; **e**, Early Medieval; **f**, Late Medieval. Modeled ancestry proportions are indicated by sample size-scaled pie charts, with ancestry source populations shown below. Cultural groups are indicated by bold text. For panels **d-f**, individuals are Late Xiongnu, Türkic, and Mongol, respectively, unless otherwise noted. Previously published reference populations are noted with white text; all others are from this study. Populations beyond the map borders are indicated by arrows. Burial locations have been jittered to improve visibility of overlapping individuals. Zoom in to see individual labels. Here we report results from admixture models that include all ancestry components required to explain historic late Medieval individuals as a group for unbiased cross comparison between individuals. Individual results with simpler admixture models can be found in Table S23.

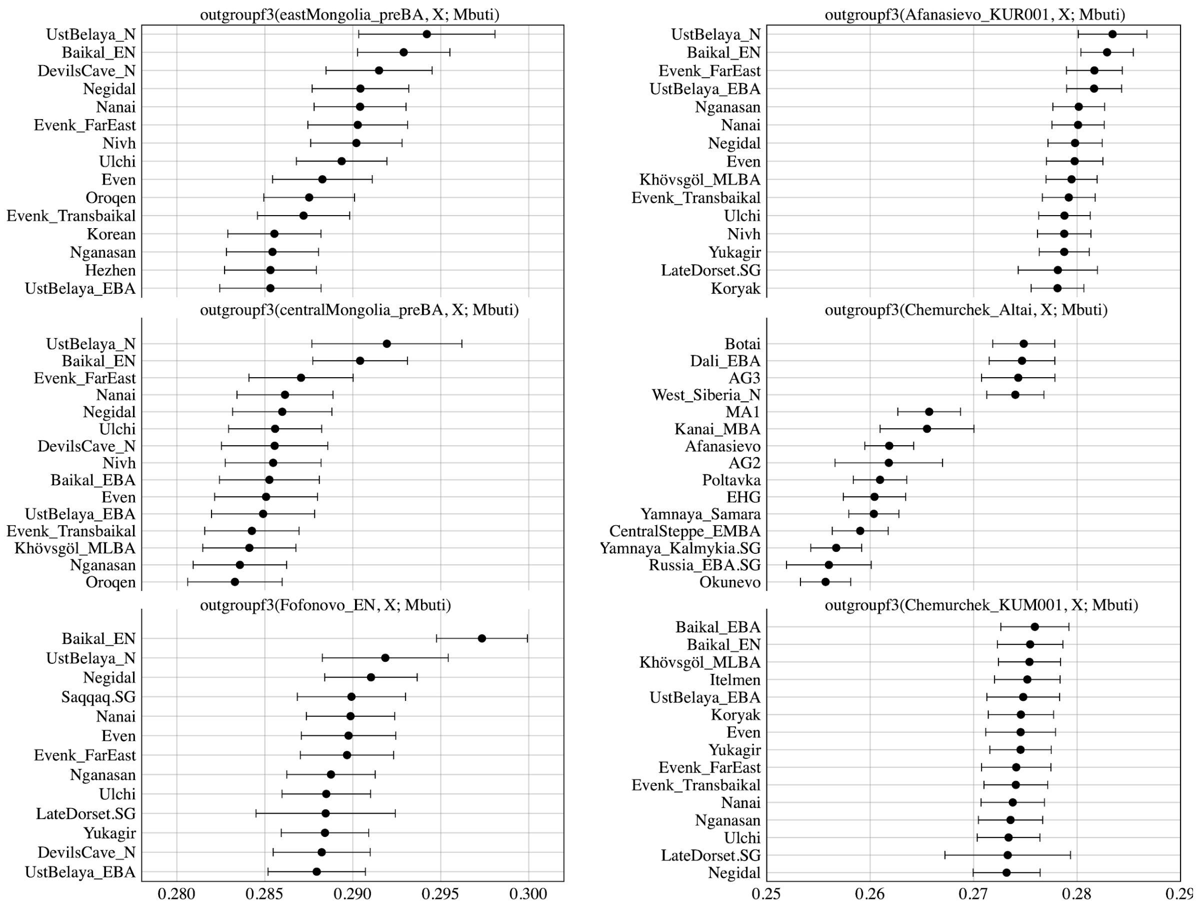

**Figure S9. Outgroup *f3*-statistics for the pre-Bronze Age to Early Bronze Age groups in the Eastern Steppe.** We show top 15 outgroup *f3*-statistics of the form *f3*(Target, world-wide; Mbuti out of 345 ancient and present-day populations for the six target groups: eastMongolia_preBA, centralMongolia_preBA, Fofonovo_EN, Afanasievo_KUR001, Chemurchek_Altai and Chemurchek_KUM001. Horizontal bars represent ±1 standard error (SE) calculated by 5 cM block jackknifing.

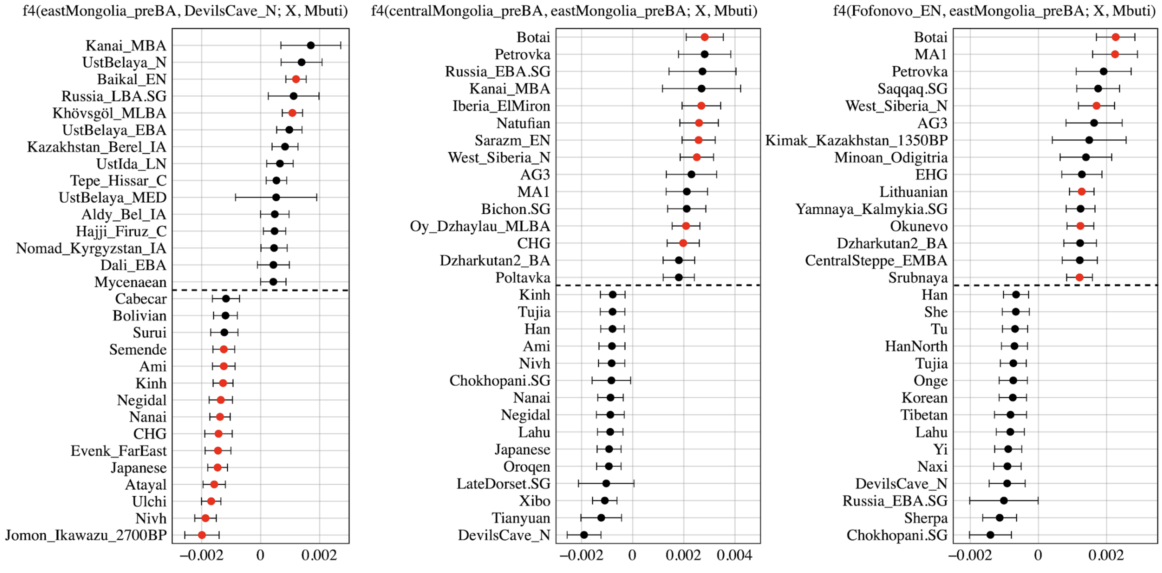

**Figure S10. Testing cladality of the four ANA populations using *f4*-statistics.** We show top and bottom 15 symmetric *f4*-statistics of the form *f4*(ANA1, ANA2; world-wide, Mbuti) out of 345 ancient and present-day populations for the four ANA-related target groups: eastMongolia_preBA, centralMongolia_preBA, Fofonovo_EN, DevilsCave_N. Horizontal bars represent ±1 standard error (SE) calculated by 5 cM block jackknifing. *f4*-statistics with Z-score>3 are highlighted in red.

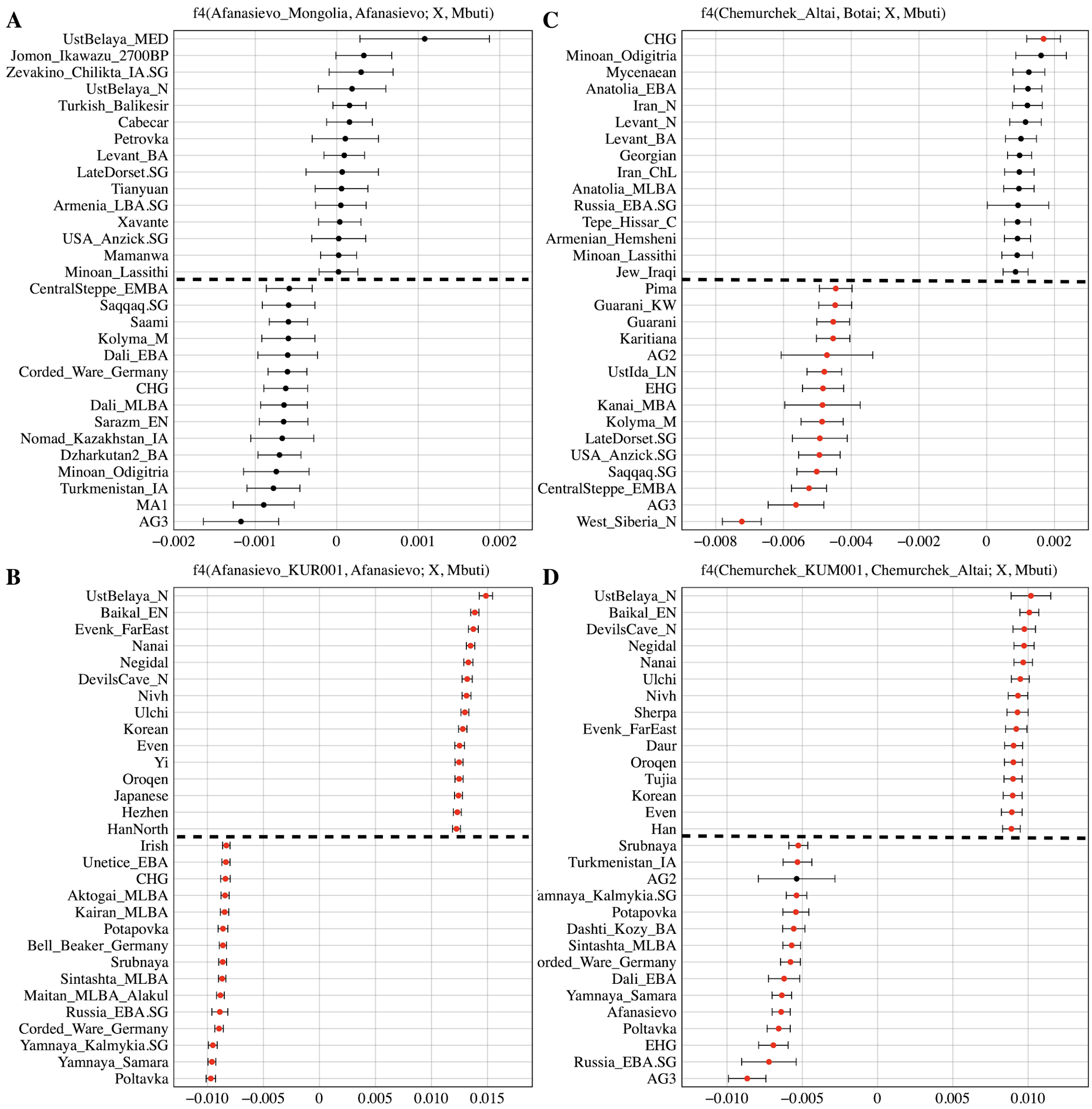

**Figure S11. Testing cladality of Afanasievo and Chemurchek using *f4*-statistics**

We show top and bottom 15 symmetric *f4*-statistics for the four target groups— Afanasievo_Mongolia, Afanasievo_KUR001, Chemurchek_Altai and Chemurchek_KUM001—in the form (**A**) *f4*(Afanasievo_Mongolia, Afanasievo; world-wide, Mbuti), (**B**) *f4*(Afanasievo_KUR001, Afanasievo; world-wide, Mbuti), (**C**) *f4*(Chemurchek_Altai, Botai; world-wide, Mbuti), (**D**) *f4*(Chemurchek _KUM001, Chemurchek_Altai; world-wide, Mbuti), out of 345 ancient and present-day populations. Horizontal bars represent ±1 standard error (SE) calculated by 5 cM block jackknifing. *f4*-statistics with Z-score>3 are highlighted in red.

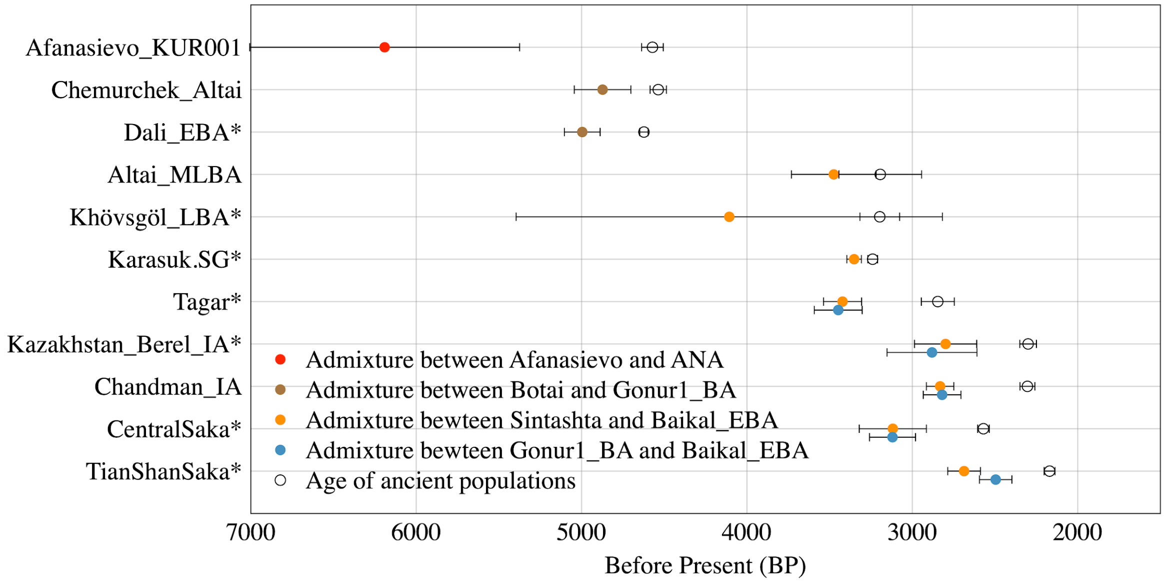

**Figure S12. Dating admixture in prehistoric individuals.** We estimated admixture dates using the DATES program and converted it by adding the age of each ancient population (mean value of the center of the 95% confidence interval of calibrated $^{14}C$ dates) and assuming 29 years per generation. Horizontal bars associated with the admixture dates (colored circles) are estimated by the square root of summing the variance of DATES estimate using leave-one-chromosome-out jackknifing method and the variance of the $^{14}C$ date estimate, assuming that the two quantities are independent. Published groups are marked with asterisk*.

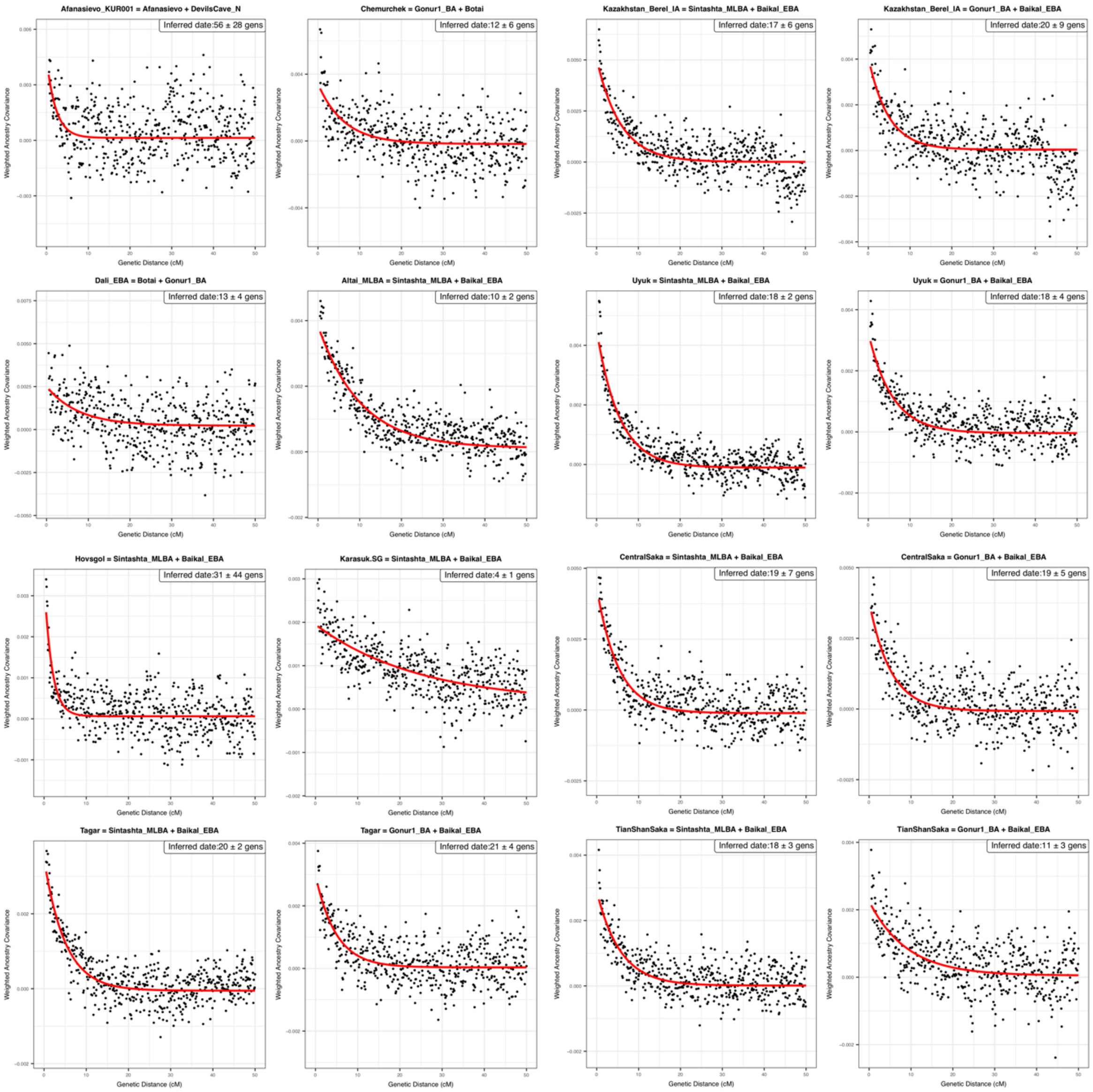

**Figure S13. Ancestry covariance in prehistoric individuals.** We show the weighted ancestry covariance (y-axis) calculated from DATES which is expected to decay exponentially along genetic distance (x-axis) with a decay rate indicating the time since admixture, and fitted exponential curves (shown in red line) for every admixture model summarized in Figure S12 above. We start the fit at genetic distance at 0.45 centiMorgans, and estimate standard error by a weighted block jackknife removing one chromosome in each run.

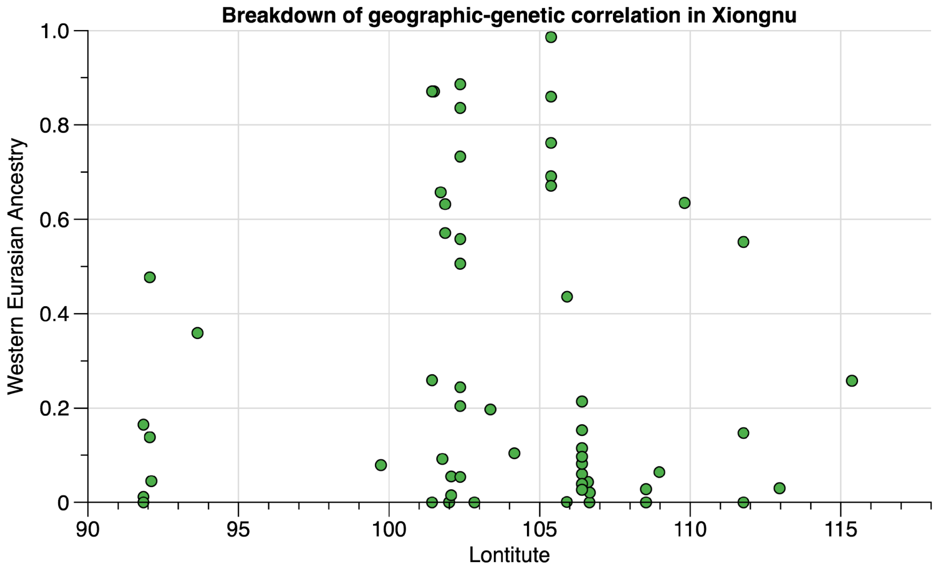

**Figure S14. Breakdown of the geographic-genetic correlation in Xiongnu.** We show the proportions of West Eurasian ancestry on all individuals/groups from Xiongnu era (y axis) versus the longitude of archaeological site they come from (x axis). The raw numbers of individual estimates can be found in Table S20 for models using Sarmatian as the western Eurasian source. Unlike MLBA/EIA individuals (Fig. 3), Xiongnu individuals from more western sites do not have higher proportion of western Eurasian ancestry than those from eastern sites.

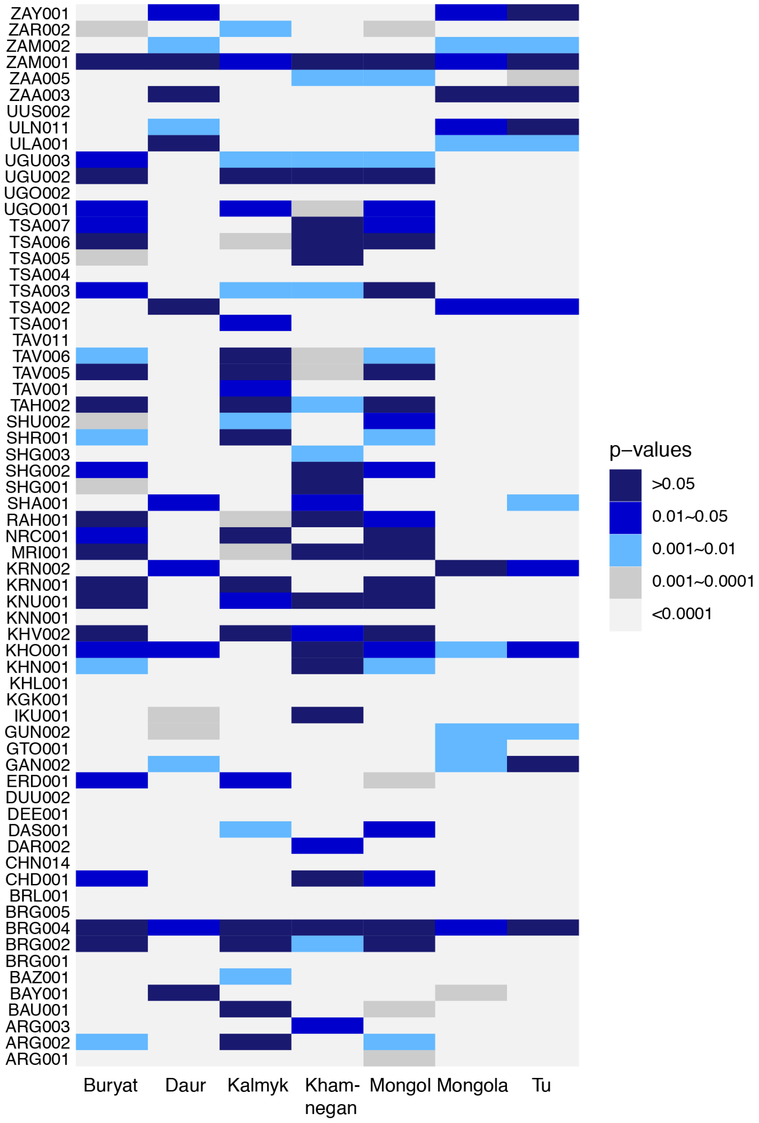

**Figure S15. Comparing genetic homogeneity between ancient Mongol individuals and 7 present-day Mongolic-speaking populations using qpWave.** We report the *p*-value for every individual-based qpWave {ancient Mongol individual; Mongolic group} using seven modern Mongolic-speaking populations: Buryat, Daur, Kalmyk, Khamnegan, Mongol, Mongola, and Tu in the Human Origins dataset. When the *p*-value from qpWave is >0.05, it suggests that the ancient individual on the y axis is genetically indistinguishable from the modern Mongolic-speaking population shown on the x axis. Smaller *p*-values indicate that the ancient individual is significantly different from the modern group.
